# A non-canonical RNA-binding domain of the Fragile X protein, FMRP, elicits translational repression independent of mRNA G-quadruplexes

**DOI:** 10.1101/2022.01.10.475703

**Authors:** MaKenzie R. Scarpitti, Julia E. Warrick, Evelyn L. Yoder, Michael G. Kearse

**Affiliations:** The Biomedical Sciences Graduate Program, The Ohio State University, Columbus, OH 43210 USA; Department of Biological Chemistry and Pharmacology, The Ohio State University, Columbus, OH 43210 USA; Center for RNA Biology, The Ohio State University, Columbus, OH 43210USA

## Abstract

Loss of functional fragile X mental retardation protein (FMRP) causes fragile X syndrome, the leading form of inherited intellectual disability and the most common monogenic cause of autism spectrum disorders. FMRP is an RNA-binding protein that controls neuronal mRNA localization and translation. FMRP is thought to inhibit translation elongation after being recruited to target transcripts via binding RNA G-quadruplexes (G4s) within the coding sequence. Here, we directly test this model and report that FMRP inhibits translation independent of mRNA G4s.

Furthermore, we found that the RGG box motif together with its natural C-terminal domain forms a non-canonical RNA-binding domain (ncRBD) that is essential for translational repression. The ncRBD elicits broad RNA binding ability and binds to multiple reporter mRNAs and all four homopolymeric RNAs. Serial deletion analysis of the ncRBD identified that the regions required for mRNA-binding and translational repression overlap but are not identical. Consistent with FMRP stalling elongating ribosomes and causing the accumulation of slowed 80S ribosomes, transcripts bound by FMRP via the ncRBD co-sediment with heavier polysomes and were present in puromycin-resistant ribosome complexes. Together, this work identifies a ncRBD and translational repression domain that shifts our understanding of how FMRP inhibits translation independent of mRNA G4s.

## INTRODUCTION

Loss of functional fragile X mental retardation protein (FMRP) causes fragile X syndrome (FXS) (1-4), the leading form of inherited intellectual disability and the most common monogenic cause of autism spectrum disorders (ASD) (5). FXS affects around one in 4,000 males and one in 8,000 females (5). Approximately one third of FXS patients are also diagnosed with ASD (6). The vast majority of FXS cases is caused by a CGG trinucleotide repeat expansion in the 5′ untranslated region (UTR) of the *FMR1* gene. The expanded repeat is subsequently hypermethylated which causes transcriptional silencing of the locus (1-4). As a result, most FXS patients express little to no functional FMRP.

FMRP is an RNA-binding protein (RBP) with multiple RNA-binding domains (RBDs) including three K homology (KH) domains (KH0, KH1, and KH2) and a positively-charged RGG box motif (7-9). In general, KH domains canonically bind to short stretches of RNA to provide binding specificity to particular transcripts (10-14). However, the KH domains of FMRP are unable to bind strongly to any combination of five ribonucleotides (15). *In vitro* selection experiments have found that the KH2 domain of FMRP does have high affinity for an artificial RNA pseudoknot (e.g., Δ kissing complex 2; Δkc2) (16). Reports using *in vitro* selection, filter binding, and fluorescence anisotropy have concluded that the RGG box motif of FMRP has preference for RNA G-quadruplexes (G4s), and in particular an artificial RNA molecule called Sc1 that harbors a G4 (15,17). Using these multiple binding domains and motifs, FMRP is thought to bind mRNAs with higher order RNA structure (e.g., pseudoknots and G4s). The importance of the KH domains and RGG box motif in FMRP function is highlighted by independent point mutations found in rare FXS cases. Enigmatically, a G266E mutation in the KH1 domain (18), an I304N mutation in the KH2 domain (19), and a guanine insertion that causes a frameshift early within the RGG box motif (20) all cause FXS. This suggests that FMRP may have multiple functions within neurons dictated by interactions between specific RBDs and mRNA sequences or structures.

FMRP is known to regulate translation and, in alignment with this idea, it is found to primarily bind the coding sequence (CDS) of mRNAs in mouse brain tissue (21). Most previous reports (21-25), but not all (26-28), suggest that FMRP represses translation of its target mRNAs by stalling bound elongating ribosomes. Thus, at least one facet of the FXS phenotype is thought to result from aberrant and unregulated protein synthesis of dendritic mRNAs at synapses. Because FMRP mutants that lack the RGG box motif do not repress translation (22), it is commonly postulated that FMRP binds to target mRNAs at RNA G4s and subsequently blocks translation elongation. In this model, the G4-bound FMRP acts as a physical roadblock and directly contacts the elongating ribosome near the A site to sterically hinder delivery of aminoacyl-tRNAs to the ribosome.

Here we directly test the ability of FMRP to inhibit translation of G4-lacking and G4-containing mRNAs. Our data show that FMRP represses translation independent of mRNA G4s and that the RGG box works synergistically with the positively-charged C-terminal domain (CTD) to form a non-canonical RNA-binding domain (ncRBD) that is critical for repression. The RGG+CTD ncRBD is able to bind all four homopolymeric RNA sequences with slightly less preference for homopolymeric A RNA, providing FMRP the ability to target multiple mRNAs.

Through scanning deletion analysis of the ncRBD, we found that the residues required for mRNA binding and translational repression overlap but are not identical. Consistent with inhibiting translation post-initiation, our data show that FMRP harboring the ncRBD, but not a mutant that is missing the ncRBD, promotes accumulation of inhibited reporter mRNA on heavy polysomes and forms puromycin-resistant ribosome complexes. Taken together, our data indicate that FMRP harbors a small RNA-binding element that extends from the annotated RGG box motif that overlaps with a translational repression domain to stall and slow elongating ribosomes independent of mRNA G4s.

## MATERIALS AND METHODS

### Plasmids

The nLuc coding sequence from pNL1.1 (Promega) was analyzed by Quadruplex forming G-Rich Sequences (QGRS) Mapper (https://bioinformatics.ramapo.edu/QGRS/index.php) and manually codon optimized to eliminate predicted G4 motifs. The final nLuc coding sequence harboring the P2A ribosome skipping motif and human β-globin 5′ UTR was synthesized by Integrated DNA Technologies and cloned into pcDNA3.1(+). The G4 sequence and larger 5′ UTR (three human β-globin 5′ UTR sequences in tandem) were inserted using the Q5 Site-Directed Mutagenesis Kit (NEB # E0552S). pcDNA3.1(+)/mEGFP was a kind gift from Jeremy Wilusz (Baylor College of Medicine). pCRII/FFLuc, which contains the FFLuc coding sequence from pGL4.13 (Promega) downstream from the T7 RNA polymerase promoter, was previously described (29).

An *E. coli* optimized coding sequence for human FMRP (isoform 1) was designed and synthesized by Genscript, and then subcloned into pET His6 MBP TEV LIC cloning vector (1M), which was a gift from Scott Gradia (Addgene plasmid # 29656), through ligation-independent cloning (LIC) using Novagen’s LIC-qualified T4 DNA polymerase (Sigma # 70099-M) as described by Q3 Macrolab (http://qb3.berkeley.edu/macrolab/). The His6-tag was deleted from the N-terminus and inserted at the C-terminus. The NT-hFMRP sequence included a P451S mutation to prevent ribosome stalling at a poly-proline stretch and formation of truncated recombinant protein, as previously described (30). Point mutations and deletions were achieved using the Q5 Site-Directed Mutagenesis Kit. To be as consistent as possible across the previous literature, we refer to the RGG box motif as to the minimal region identified by Darnell and colleagues that bound Sc1 RNA with highest affinity (17).

All plasmids were propagated in TOP10 *E. coli* (Thermo Fisher # C404006), purified using the PureYield Plasmid Miniprep or Midiprep Systems (Promega # A1222 and A2495), and validated by Sanger sequencing at The Ohio State University Comprehensive Cancer Center Genomics Shared Resource (OSUCCC GSR). Nucleotide sequences of the reporters and recombinant proteins are provided in the Supplemental Material.

### Reporter mRNA *in vitro* transcription

All nLuc plasmids were linearized with XbaI and purified using a Zymo DNA Clean & Concentrator 25 (Zymo Research # D4065). pcDNA3.1(+)/mEGFP was linearized with PspOMI. pCRII/FFLuc was linearized with HindIII. DNA was transcribed into mRNA which was co-transcriptionally capped with the Anti-Reverse Cap Analog (ARCA) 3′-O-Me-m7G(5′)ppp(5′)G (NEB # S1411) using the HiScribe T7 High Yield RNA Synthesis Kit (NEB # E2040). Our standard 10 μL reactions used 0.5 μg of linear plasmid template and an 8:1 ARCA:GTP ratio.

Reactions were incubated at 30°C for 2 hrs, then incubated with 20 U of DNaseI (NEB # M0303S) at 37°C for 15 min, and then purified using a Zymo RNA Clean & Concentrator 25 (Zymo Research # R1018). Reporter mRNA was eluted in 75 μL RNase-free water, aliquoted in single use volumes, and stored at -80°C. Reporter mRNA integrity was confirmed by denaturing formaldehyde agarose gel electrophoresis and ethidium bromide visualization. We routinely found the 30°C incubation resulted in less observable truncated products than incubation at 37°C and did not significantly affect yield for our purposes.

### Recombinant protein expression and purification

All recombinant proteins were expressed in Rosetta 2(DE3) *E. coli* (Sigma # 71397-4) using MagicMedia *E. coli* Expression Medium (Thermo Fisher # K6803) supplemented with 50 μg/mL kanamycin and 35 μg/mL chloramphenicol for auto-induction. A 5 mL starter culture in LB media supplemented with 50 μg/mL kanamycin, 35 μg/mL chloramphenicol, and 1% glucose (w/v) was inoculated with a single colony and grown overnight at 37°C, 250 rpm. 1 mL of a fresh overnight starter culture was then used to inoculate 50 mL of room temperature MagicMedia and incubated for 48 hrs at 18°C, 160 rpm in a 250 mL baffled flask. After auto-induction, cultures were pelleted and stored at -20°C for purification later. Recombinant proteins were purified using a dual affinity approach, first using the C-terminal His6-tag, then the N-terminal MBP-tag. Cell pellets were resuspended and lysed with BugBuster Master Mix (Sigma # 71456) using the recommended 5 mL per 1 g wet cell pellet ratio for 10 min at room temperature with gentle end-over-end rotation (10-15 rpm). Lysates were placed on ice and kept cold moving forward. Lysates were cleared by centrifugation for 20 min at 18,000 rcf in a chilled centrifuge (4°C). Lysates were then incubated with HisPur Cobalt Resin (Thermo Fisher # 89965) in a Peirce centrifugation column (Thermo # 89897) for 30 min at 4°C with gentle end-over-end rotation. Columns were centrifuged in a pre-chilled (4°C) Eppendorf 5810R for 2 min at 700 rcf to eliminate the flow through and then were washed 5X with two resin-bed volumes of ice-cold Cobalt IMAC Wash Buffer (50 mM Na_3_PO_4_, 300 mM NaCl, 10 mM imidazole; pH 7.4) in a pre-chilled (4°C) Eppendorf 5810R for 2 min at 700 rcf. His-tagged proteins were then eluted in a single elution step with two resin-bed volumes of ice-cold Cobalt IMAC Elution Buffer (50 mM Na_3_PO_4_, 300 mM NaCl, 150 mM imidazole; pH 7.4) by gravity flow. Eluates were then incubated with Amylose resin (NEB # E8021) in a centrifugation column for 2 hrs at 4°C with gentle end-over-end rotation. Columns were washed 5X with at least two bed-volumes of ice-cold MBP Wash Buffer (20 mM Tris-HCl, 200 mM NaCl, 1 mM EDTA; pH 7.4) by gravity flow. MBP-tagged proteins were then eluted by a single elution step with two resin-bed volumes of ice-cold MBP Elution Buffer (20 mM Tris-HCl, 200 mM NaCl, 1 mM EDTA, 10 mM maltose; pH 7.4) by gravity flow. Recombinant proteins were then desalted and buffer exchanged into Protein Storage Buffer (25 mM Tris-HCl, 125 mM KCl, 10% glycerol; pH 7.4) using a 7K MWCO Zeba Spin Desalting Column (Thermo Fisher # 89892) and, if needed, concentrated using 10K MWCO Amicon Ultra-4 (EMD Millipore # UFC803024). Recombinant protein concentration was determined by Pierce Detergent Compatible Bradford Assay Kit (Thermo Fisher # 23246) with BSA standards diluted in Protein Storage Buffer before aliquoting in single use volumes, snap freezing in liquid nitrogen, and storage at -80°C.

### mRNA folding, mRNP formation, and *in vitro* translation

*In vitro* translation was performed in the dynamic linear range as previously described but adapted to translate mRNPs (29). 28 nM *in vitro* transcribed reporter mRNA in RNA folding buffer (10 mM Tris-HCl, 100 mM KCl; pH 7.4) was heated for 5 min at 70°C then gradually cooled for 30 min at room temperature on the bench. 5 mM Mg(OAc)_2_ (final) was then added, gently mixed, and allowed to cool for an additional 30 min on the bench. In a total of 4 μL, 28 fmol of folded reporter mRNA was mixed with 0-10 picomol of recombinant protein and 100 picomol of UltraPure BSA (Thermo Fisher # AM2618) on ice for 1 hr. UltraPure BSA stock was diluted in protein storage buffer and its addition was necessary to prevent non-specific binding of the reporter mRNA to the tube. For *in vitro* translation reactions, 6 μL of a Rabbit Reticulocyte Lysate (RRL) master mix was added to each 4μL mRNP complex. 10 μL *in vitro* translation reactions were performed in the linear range using 2.8 nM mRNA in the Flexi RRL System (Promega # L4540) with final concentrations of reagents at 30% RRL, 10 mM amino acid mix minus leucine, 10 mM amino acid mix minus Methionine, 0.5 mM Mg(OAc)_2_, 100 mM KCl, 8 U murine RNase inhibitor (NEB # M0314), 0-1 μM recombinant protein, and 10 μM UltraPure BSA. Reactions were incubated for 30 min at 30°C, terminated by incubation on ice and diluted 1:5 in Glo Lysis Buffer (Promega # E2661). 25 μL of prepared Nano-Glo reagent (Promega # N1120) was mixed with 25 μL of diluted reaction and incubated at room temperature for 5 min in the dark (with gentle shaking during the first minute), and then read on a Promega GloMax Discover Multimode Microplate Reader.

IC_50_ measurements were calculated using non-liner regression analysis following the [inhibitor] versus normalized response analysis in GraphPad Prism 9.1.2

### Western blot of *in vitro* translation reactions

10 μL translation reactions were performed with 2.8 nM mEGFP mRNA (folded) and 1 μM WT NT-hFMRP as described above with nLuc mRNA. 40 μL of 2X reducing SDS sample buffer (Bio-Rad # 1610737) was then added and heated at 70°C for 15 min. 10 μL was then separated by standard Tris-Glycine SDS-PAGE (Thermo # XP04200BOX) and transferred on to 0.2 μm PVDF membrane (Thermo # 88520). Membranes were then blocked with 5% (w/v) non-fat dry milk in TBST (1X Tris-buffered saline with 0.1% (v/v) Tween 20) for 30 min at room temperature before overnight incubation with primary antibodies in TBST at 4°C. After three 10 min washes with TBST, membranes were incubated with HRP-conjugated secondary antibody in TBST for 1 hr at room temperature and then washed again with three 10 min washes with TBST. Chemiluminescence was performed with SuperSignal West Pico PLUS for GFP (Thermo # 34577) and with SuperSignal West Femto Maximum Sensitivity Substrate (Thermo # 34095) for tubulin. Blots were imaged using an Azure Sapphire Biomolecular Imager. Rabbit anti-GFP (Cell Signaling # 2956S) was used at 1:1,000. Mouse anti-tubulin (Sigma # T9026-2ML) was used at 1:1,000. HRP-conjugated goat anti-rabbit IgG (H+L) (Thermo # 31460) was used at 1:60,000 for GFP and HRP-conjugated goat anti-mouse IgG (H+L) (Thermo # 31430) was used at 1:10,000 for tubulin.

### Denaturing PAGE, native PAGE, and nucleic acid staining

Denaturing TBE-Urea 6% PAGE gels (Thermo Fisher # EC68652BOX) were run with 1X TBE-Urea Sample Buffer (Thermo Fisher # LC6876) and 1X TBE Running Buffer (Thermo Fisher # LC6675). Gels were pre-run at 180 volts (constant) for 20 minutes, then samples were loaded and run at 180 volts (constant) for 3 hrs. Native TBE 6% PAGE gels (Thermo Fisher # EC62652BOX) were run with 1X Hi-Density TBE Sample Buffer (Thermo Fisher # LC6678) and 1X TBE Running Buffer. Gels were pre-run at 180 volts (constant) for 1 hr, then samples were loaded and run at 180 volts (constant) for 3 hrs. Total RNA was stained with 1X SYBR Green II RNA Gel Stain (Thermo Fisher # S7568) diluted in milliQ water for 10 min in the dark. G4s were selectively stained with 0.1 mg/mL N-methyl mesoporphyrin IX (NMM) (Frontier Scientific # NMM58025MG) in milliQ water for 10 min in the dark. Stained gels were imaged on a Bio-Rad GelDoc Go Gel Imaging System using the SYBR Green setting. NMM stock was made at 10 mg/mL in DMF and stored in single use aliquots at -20°C. For Native TBE PAGE, the mRNA was folded as described above. Due to the different sensitivity, 100 ng and 2,000 ng were loaded for staining with SYBR Green II and NMM, respectively.

### Electrophoretic mobility shift assays

For EMSAs with mRNAs to assess mRNP formation, 4% PAGE gels made with 0.5X TBM (45 mM Tris, 45 mM borate, 2.5 mM MgCl_2_) and acrylamide/bis-acrylamide, 37.5:1 (2.7% crosslinker) were poured between glass plates and allowed to polymerize for at least 1 hr. Gels were pre-run for 20 min at 100 volts (constant) with 0.5X TBM as the running buffer. In 18 μL, 0.4 picomol of folded reporter mRNA was mixed with 20 picomol of recombinant protein and 200 picomol of UltraPure BSA (Thermo Fisher # AM2618) in binding buffer (10 mM Tris-HCl, 100 mM KCl, 5 mM Mg(OAc)_2_; pH 7.4) on ice for 1 hr. 2 μL of 20% Ficoll 400 (Sigma # F5415-25ML) was then added and the entire sample was loaded immediately. After loading, gels were run for 45 min at 100 volts (constant) at room temperature, stained with 1X SYBR Green II RNA Gel Stain, and visualized as described above.

For EMSAs with 5’ FAM-labeled RNA oligos, 4% PAGE gels were made as described above with 0.5X TBM and acrylamide/bis-acrylamide, 37.5:1. 5’ FAM-labeled RNA olgios were first diluted in nuclease-free water and not heated. In 7 μL, 0.2 picomol of 5’ FAM-labeled RNA oligo was mixed with 22 picomol of recombinant protein and 150 picomol of UltraPure BSA in binding buffer (10 mM Tris-HCl, 100 mM KCl, 5 mM Mg(OAc)_2_; pH 7.4) on ice for 1 hr in the dark. 2 μL of 20% Ficoll 400 was then added, and the entire sample was loaded immediately. For the FAM-U(G)17 RNA oligo, 0.4 picomol per reaction was used due to FAM being slightly quenched by proximal guanosines (a known caution provided by the RNA oligo manufacture). Gels were pre-run for 20 min at 100 volts (constant) with 0.5X TBM as the running buffer.

Samples were loaded and gels were run for 40 min at 100 volts (constant) at room temperature in the dark. Gels in glass plates were then directly imaged using an Azure Sapphire Biomolecular Imager. The 5′ FAM-labeled RNA oligo sequences are provided in the Supplemental Data. A single uridine was added as a spacer between the 5’ FAM label and the polymeric guanosines to avoid quenching and was kept for consistency in the other labeled polymeric RNA oligos. We found flexible linkers did not help further avoid quenching by proximal guanosines.

### Fluorescence polarization assays

In 100 μL, 5.5 picomol of 5’ FAM-labeled RNA oligo was mixed with 300 picomol of recombinant protein and 2,000 picomol of UltraPure BSA in binding buffer (10 mM Tris-HCl, 100 mM KCl, 5 mM Mg(OAc)_2_; pH 7.4) on ice for 1 hr in the dark. 30 μL of each reaction was then added to non-binding half-area black 96-well plate (Corning # 3993) and fluorescence polarization was measured on a Tecan Spark equipped with an enhanced fluorescence module.

### Sucrose gradient ultracentrifugation and RT-qPCR

*In vitro* translation reactions were scaled up 10-fold to 100 μL but were performed identically as described above. After 30 min at 30°C, reactions were kept on ice, diluted two-fold with ice-cold Polysome Dilution Buffer (10 mM Tris-HCl, 140 mM KCl, 10 mM MgCl_2_, 1 mM DTT, 100 μg/mL cycloheximide; pH 7.4), and layered on top of a linear 10-50% (w/v) buffered sucrose gradient (10 mM Tris-HCl, 140 mM KCl, 10 mM MgCl_2_, 1 mM DTT, 100 μg/mL cycloheximide; pH 7.4) in a 14 mm × 89 mm thin-wall Ultra-Clear tube (Beckman # 344059) that was formed using a Biocomp Gradient Master. Gradients were centrifuged at 35K rpm for 120 min at 4°C in a SW-41Ti rotor (Beckman) with maximum acceleration and no brake using a Beckman Optima L-90 Ultracentrifuge. Gradients were subsequently fractionated into 0.9 mL volumes using a Biocomp piston fractionator with a TRIAX flow cell (Biocomp) recording a continuous A_260 nm_ trace. Total RNA was extracted from 400 μL of each fraction (spiked with 0.2 ng exogenous control FFLuc mRNA; Promega # L4561) by adding 600 μL TRIzol (Thermo Fisher # 15596018) and following the manufacturer’s protocol. Glycogen (Thermo Fisher # R0561) was added at the isopropanol precipitation step. The resulting RNA pellet was resuspended in 30 μL nuclease-free water. 16 μL of extracted RNA was converted to cDNA using iScript Reverse Transcription Supermix for RT-qPCR (Bio-Rad # 1708841). cDNA reactions were then diluted 10-fold with nuclease-free water and stored at -20°C or used immediately. RT-qPCR was performed in 15 μL reactions using iTaq Universal SYBR Green Supermix (Bio-Rad # 1725124) in a Bio-Rad CFX Connect Real-Time PCR Detection System with 1.5 μL diluted cDNA and 250 nM (final concentration) primers. For each fraction, nLuc reporter mRNA abundance was normalized to the spiked-in control FFLuc mRNA using the Bio-Rad CFX Maestro software (ΔΔCt method).

Abundance of total signal in each fraction was calculated using *Q*_*n*_ *= 2*^*ΔΔCt*^ and *P = 100 × Q*_*n*_*/Q*_*total*_ as previously described (31). Primers for RT-qPCR can be found in **Supplemental Table S1**.

### Puromycin treatment and low-speed sucrose cushions

nLuc reporter mRNA translation was performed as described above except that translation was limited to 15 min at 30°C. Samples were then placed on ice for 3 min before the addition of 0.1 mM puromycin (final) and further incubation at 30°C for 30 min. Control samples lacking puromycin (water added instead) were kept on ice. Cycloheximide (final concentration of 1.43 mg/mL) was then added to all samples to preserve ribosome complexes on mRNAs and halt puromycin incorporation. In a separate tube, Firefly luciferase (FFLuc) reporter mRNA was translated as described above (3 nM mRNA conditions) for 15 min at 30°C and was terminated by the addition of 1.43 mg/mL cycloheximide (final) and incubation on ice.

nLuc and FFLuc translation reactions were combined on ice and then mixed with an equal volume of ice-cold 2X dilution buffer (40 mM Tris-HCl, 280 mM KCl, 20 mM MgCl_2_, 200 μg/ml cycloheximide, 2 mM DTT; pH 7.4). The entire 40 μl volume was then layered on top of 130 μl of ice-cold 35% (w/v) buffered sucrose (20 mM Tris-HCl, 140 mM KCl, 10 mM MgCl_2_, 100 μg/mL cycloheximide, 1 mM DTT; pH 7.4) in a pre-chilled 7 mm x 20 mm thick-walled polycarbonate ultracentrifuge tubes (Thermo Scientific # 45233) and centrifuged in a S100AT3 rotor at 4°C for 60 min at 50,000 x g (43,000 rpm) in a Sorvall Discovery M120 SE Micro-Ultracentrifuge. The supernatant was then discarded and each pellet was resuspended in 0.5 mL of TRIzol (Thermo Fisher # 15596018). Total RNA was extracted from each pellet following the manufacturer’s protocol with glycogen (Thermo Fisher # R0561) added at the isopropanol precipitation step. The resulting RNA pellet was resuspended in 30 μL nuclease-free water. 16 μL of extracted RNA was converted to cDNA and analyzed by RT-qPCR as described in above. For each sample, nLuc reporter mRNA abundance was normalized to FFLuc mRNA (coding sequence from pGL4.13) using the Bio-Rad CFX Maestro software. See **Supplemental Figure S10** for flowchart. Primers for RT-qPCR can be found in **Supplemental Table S1**.

## RESULTS

### FMRP inhibits translation independent of mRNA G4s

Previous reports have shown that deletion of the RGG box motif from FMRP abolishes translational repression and ribosome binding *in vitro* (22). Dependence of the RGG box motif provided support to others that FMRP must target mRNAs and/or ribosomes by binding to intramolecular RNA G4s (22,24). When tested in isolation, the FMRP RGG box motif has high affinity for G-rich sequences that can form RNA G4 structures (15,17,32,33). Together, these data shaped the leading model that FMRP binds to RNA G4s in the CDS via the RGG box motif and then sterically blocks the A site of an elongating ribosome (22,24). This model has yet to be directly tested as most reporters used by the field harbor predicted G4 sequences and altering the sequence would mutate the reporter protein. To our knowledge, it has yet to be shown experimentally that RNA G4s are present and required in coding sequences for FMRP to inhibit translation.

To directly test this model, we first dual-affinity purified the N-terminally truncated human FMRP (NT-hFMRP) (**Figure 1A, B**) because it is more stable than the full-length isoform and it retains translational repression activity (22,30). We then generated specialized nLuc reporters that either lacked or harbored one of two RNA G4 structures within the CDS (**Figure 1C**).

**Figure 1.**
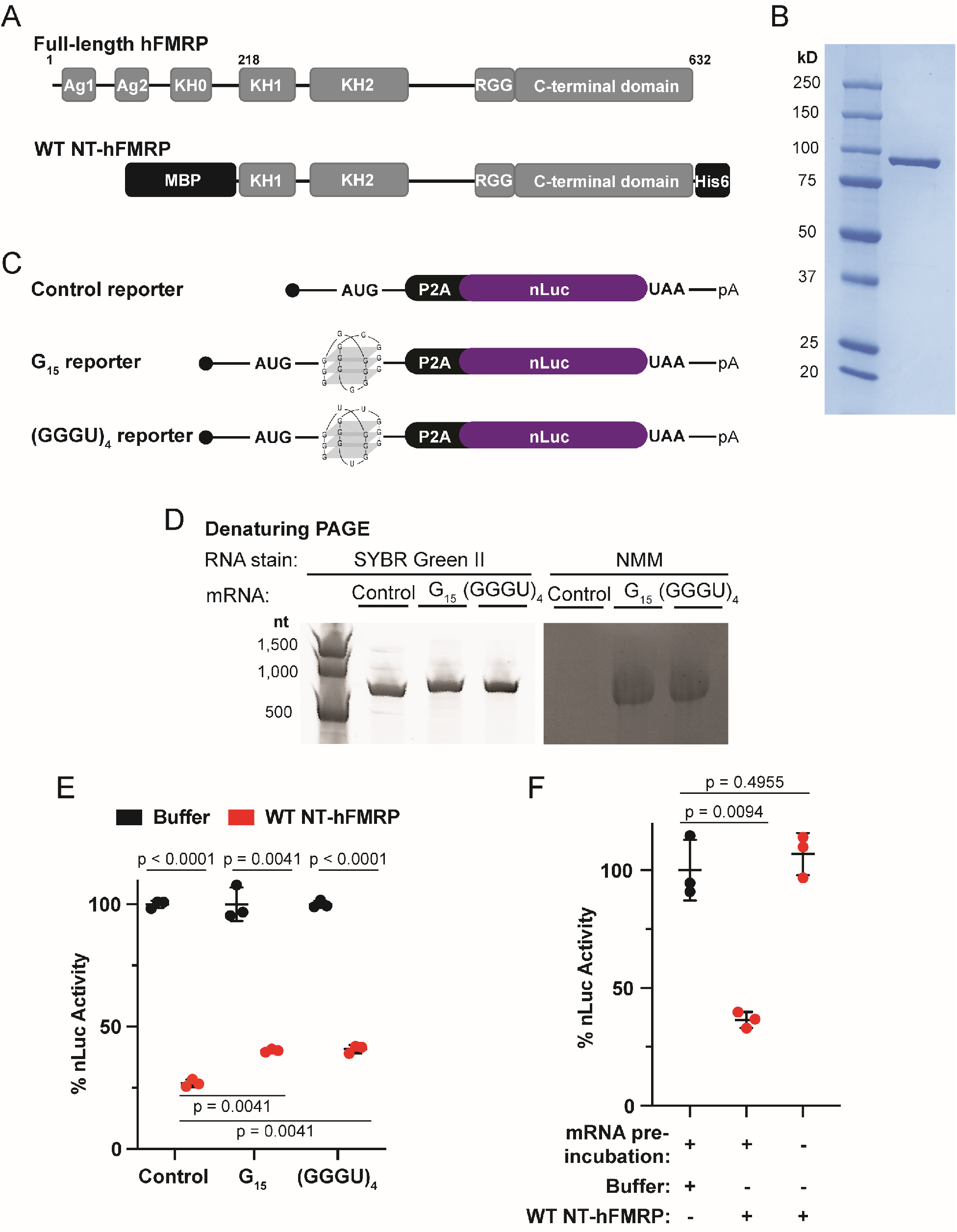
FMRP inhibits translation independent of mRNA G-quadruplexes in the CDS. A) Schematic of full-length (residues 1-632) and MBP- and His6-tagged WT N-terminally truncated human FMRP isoform 1 (NT-hFMRP). The Agenet 1 (Ag1), Agenet 2 (Ag2), and KH0 domains are absent in WT NT-hFRMP. Ag1 and Ag2 are also referred to as Tudor domains in some previous literature. WT NT-hFRMP harbors residues 218-632 of full-length human FMRP isoform 1. B) Coomassie stain of recombinant WT NT-hFMRP. C) Schematic of custom nLuc reporters either lacking a G4 (control reporter) or harboring a G4 in the coding sequence (G_15_ and (GGGU)_4_ reporters). A P2A ribosome skipping motif was included immediately upstream of the nLuc coding sequence to ensure equal nLuc function between reporters. D) Denaturing PAGE of control, G_15_, and (GGGU)_4_ reporters stained for total RNA with SYBR Green II or for G4 structures with NMM. E) *In vitro* translation of control, G_15_, and (GGGU)_4_ reporter mRNA pre-incubated with protein buffer or 1 μM WT NT-hFMRP. Data are shown as mean ± SD. n = 3 biological replicates. Comparisons were made using a two-tailed unpaired t test with Welch’s correction. F) *In vitro* translation of control nLuc reporter mRNA with protein storage buffer as a negative control, with 1 μM WT NT-hFMRP and nLuc mRNA pre-incubated together, and with 1 μM WT NT-hFMRP without a pre-incubation step. Data are shown as mean ± SD. n = 3 biological replicates. Comparisons were made using a two-tailed unpaired t test with Welch’s correction.

Importantly, this nLuc nucleotide sequence was customized to lack predicted G4 structure without altering the amino acid sequence of the reporter protein. We also included a P2A ribosome skipping motif, which releases the nascent peptide but allows the ribosome to stay bound and continue elongation, directly upstream of the nLuc CDS. This allows uniform luciferase detection across all reporters. To experimentally confirm this reporter design, we took advantage of the selective G4 staining properties of N-methyl mesoporphyrin IX (NMM) (34). As expected, the total RNA stain SYBR Green II detected both control and G4-containing nLuc reporter mRNAs (**Figure 1D, left panel**). However, NMM staining only detected the G4 reporter mRNAs (**Figure 1D, right panel**). This same selective staining pattern of NMM for the G4 reporter mRNAs was also seen in native PAGE (**Supplemental Figure S1**). These data support that only the G4 reporter mRNAs form an intramolecular RNA G4 structure (as depicted in **Figure 1C**).

We next tested to what extent recombinant NT-hFMRP represses translation of control and G4 reporter mRNAs when pre-incubated together and translated as an mRNP. If FMRP did in fact require RNA G4s on target transcripts to inhibit translation, we would expect enhanced repression on both the G_15_ and (GGGU)_4_ reporters. However, FMRP repressed translation of the G4 reporter mRNAs marginally less than the control reporter mRNA at 1 μM NT-hFMRP (**Figure 1E**). Identical results were seen with a reporter harboring the Sc1 RNA (**Supplemental Figure S2A-B**). Determining the half-maximal inhibitory concentration (IC_50_) of NT-hFMRP for each mRNA showed that the G_15_ and (GGGU)_4_ reporters were in fact ∼3-fold less sensitive to NT-hFMRP inhibition (**Supplemental Figure S2C**). These data indicate that human FMRP represses translation independent of mRNA G4s in the CDS.

For FMRP to inhibit translation elongation on select mRNAs, it is logical that it must bind target mRNAs before acting on a translating ribosome. High throughput sequencing with crosslinking immunoprecipitation of FMRP from mouse brain tissue revealed that FMRP binds to mRNA predominately in the CDS (21). However, these data are derived from multiple RBDs within FMRP, and it is not yet known which binding sites represent true FMRP translational repression targets (as opposed to mRNA transport or localization). Recombinant FMRP can also bind purified 80S ribosomes near the A site and purified 60S ribosomal subunits alone (22,24,35), raising the possibility that FMRP can directly inhibit the ribosome independent of the mRNA sequence.

To determine if FMRP requires binding to target mRNA first to inhibit translation in our assay, we performed *in vitro* translation assays using different pre-incubation protocols (**Figure 1F**). As a negative control, we programed *in vitro* translation assays with reporter mRNA and protein storage buffer. FMRP was either added directly to the translation reaction immediately before the *in vitro* translation reaction began or was allowed to first form an mRNP with reporter mRNA (which was used in **Figure 1E**). Translational repression was only observed when FMRP was pre-incubated with the nLuc reporter mRNA (**Figure 1F**), demonstrating that FMRP must bind a target mRNA first to inhibit translation. Identical results were seen by Western blot with G4-less mEGFP mRNA (**Supplemental Figure S3**). To further dissect this mechanism, we solely used the mRNA•FMRP pre-incubation strategy and the control nLuc mRNA reporter that lacks an RNA G4 for all remaining experiments.

### The RGG box motif and CTD of hFMRP together, but not independently, inhibit translation

We next sought to identify the critical RNA-binding element in FMRP required for translational repression. FMRP contains at least three canonical RBDs (KH0, KH1, KH2) and a single RGG box motif. FXS patient mutations suggest that multiple regions of FMRP are critical for RNA binding-dependent function and contribute to pathology if mutated. The I304N patient mutation in the KH2 domain abolishes FMRP binding to polysomes in human cells (16). A guanosine insertion (ΔRGG+CTD) within the sequence that encodes the RGG box motif causes a frameshift and results in a truncated FMRP that lacks most of the RGG box motif and the entire C-terminal domain (CTD) (20).

To further define the domain(s) that are critical for translational repression by FMRP, we purified recombinant NT-hFMRP harboring I304N and ΔRGG+CTD mutations (**Figure 2A, B**) and tested their ability to inhibit translation. Multiple attempts were made to purify a KH1 domain G266E mutant (18), but we were unable to recover soluble protein. We then determined the IC_50_ of WT and each mutant NT-hFMRP (**Figure 2C-F**). As expected, the His6-MBP tag alone did not inhibit translation (**Figure 2C**). WT and I304N NT-hFMRP both inhibited translation in our assay, with the I304N mutant having a ∼2-fold more potent IC_50_ than that of the WT isoform (**Figure 2D, E**). This suggests that although the I304N mutation alters FMRP binding to an optimal RNA pseudoknot substrate (i.e., Δkc2) and causes FMRP to dissociate from polysomes in cells (16), the mutation does not interfere with translational repression.

**Figure 2.**
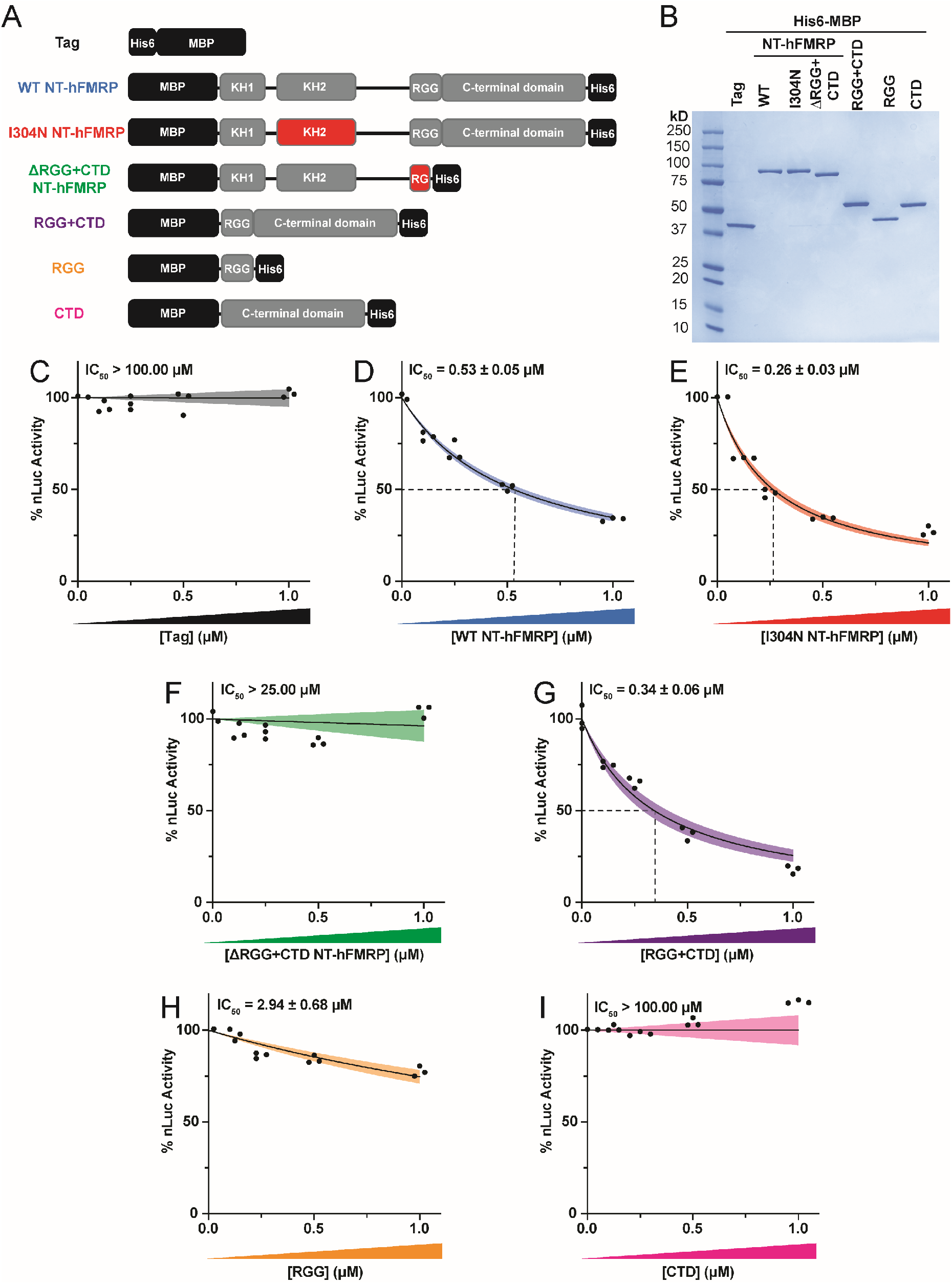
The RGG box motif and CTD together are essential and sufficient to inhibit translation. A) Schematic of recombinant WT and mutant NT-hFMRP. Mutated/truncated domains are highlighted in red. B) Coomassie stain of recombinant proteins. C-I) *In vitro* translation of nLuc mRNA with a titration of recombinant wildtype and mutant NT-hFMRP isoforms. IC_50_ values were determined for the His6-MBP tag (negative control; 633.50 ± infinity μM) (C), WT NT-hFMRP (D), I304N KH2 domain patient-derived mutant NT-hFMRP (E), ΔRGG+CTD mutant NT-hFMRP (25.72 ± infinity μM) (F), the RGG box motif + CTD fusion (G), the RGG box motif alone (H), and the CTD alone (5.75×10^33^ ± infinity μM) (I). n=3 biological replicates. A non-linear regression was used to calculate the IC_50_ and is shown as the line with the 95% confidence interval (CI) included as a watermark. The IC_50_ is reported ± 95% CI.

The ΔRGG+CTD mutant that contains the FXS-patient guanosine insertion did not repress translation (**Figure 2F**). This insertion mutation creates a frameshift in the RGG box motif and results in the encoding of a short novel peptide upstream of a premature termination codon. To validate the loss of translational repression was due to the truncation of the RGG box motif and complete deletion of the CTD rather than the addition of the short novel peptide, we purified a mutant of the NT-hFMRP that completely lacks both the RGG box motif as well as the CTD, which we termed NT-hFMRP ΔRGG+CTD complete (**Supplemental Figure S4A, B**). Full deletion of the RGG+CTD resulted in loss of translational repression (**Supplemental Figure S4C**), which is in alignment with previous reports (22,24). Taken together, the RGG+CTD is essential for translational repression.

To determine if the RGG+CTD was not only essential but also sufficient for translational repression, we purified the RGG box motif and CTD regions both together and separately (**Figure 2A, B**). Robust translational repression was observed with the isolated RGG+CTD region (**Figure 2G**). However, neither the RGG box alone nor the CTD alone effectively inhibited translation (**Figure 2H, I**). Together, these data suggest that the RGG box must be appended to the CTD to inhibit translation and that the RGG+CTD is sufficient for translational repression by FMRP (22,24).

### The RGG+CTD region forms a ncRBD that has broad RNA binding ability

We next sought to determine the RNA binding capability of the RGG+CTD region. We used electrophoretic mobility shift assays (EMSAs) to test the ability of the RGG box motif and CTD, both together and separately, to bind FAM-labeled homopolymeric RNAs. The first identified RGG box motif, belonging to heterogenous nuclear ribonucleoprotein (RNP) U, was found to bind both homopolymeric G and U RNA sequences *in vitro*, with a higher preference for polymeric G RNA (36). In agreement with the FMRP RGG box motif favoring G-rich sequences and G4s (15,37), the RGG box motif alone had some observable binding by EMSA to U(G)_17_ RNA, but little to no binding to U(A)_17_, U(C)_17_, or (U)_18_ RNA (**Figure 3A-E**). Similar results were seen with the positively charged CTD (**Figure 3A-E**). Conversely, the RGG+CTD robustly bound all four homopolymeric RNAs in EMSAs, with slightly less preference for U(A)_17_ RNA (**Figure 3A-E**). These data suggest that the RGG box motif and CTD must be together to form a non-canonical RNA-binding domain (ncRBD) that can bind to a wide range of RNAs.

**Figure 3.**
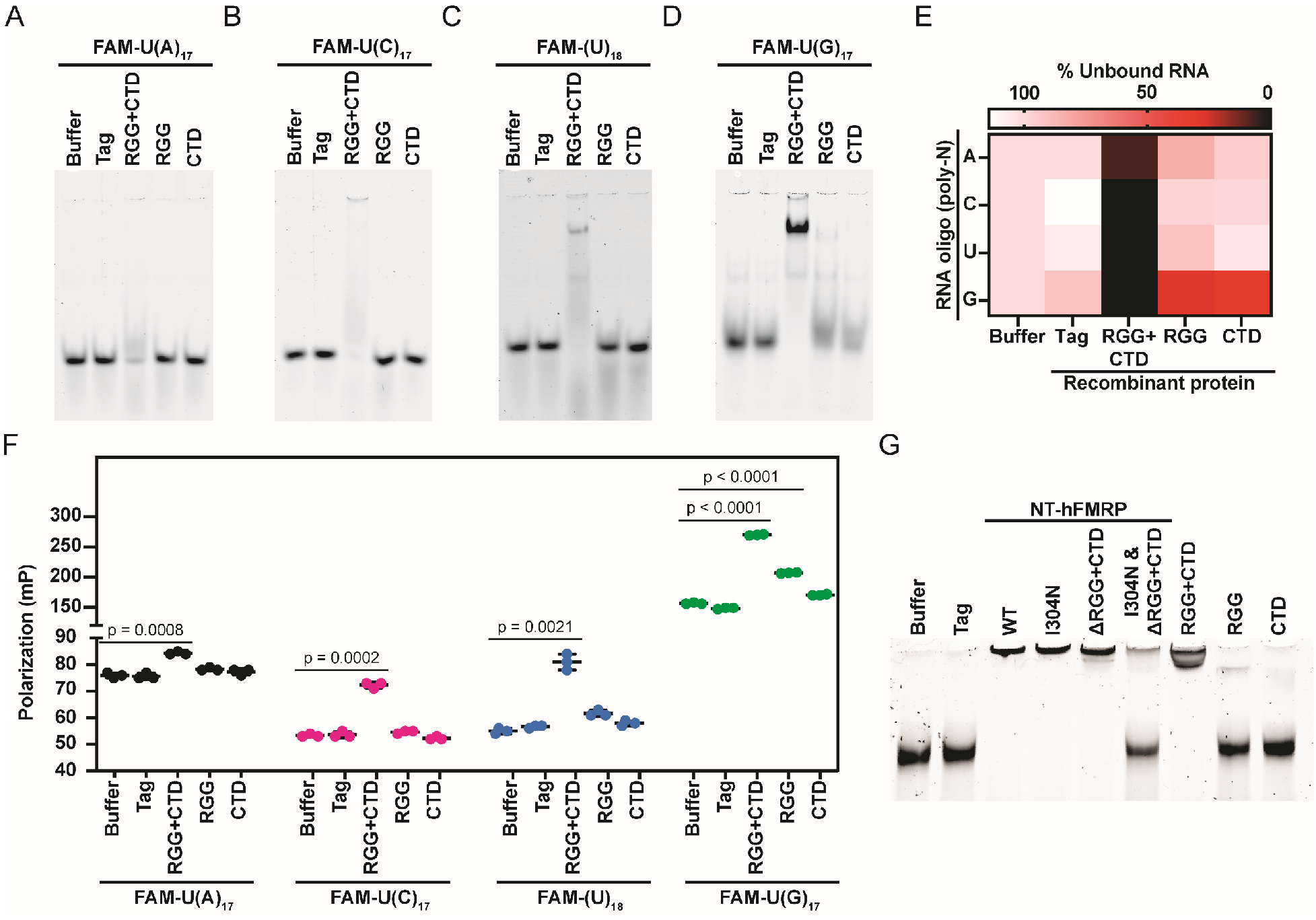
The RGG+CTD non-canonical RBD binds all four homopolymeric RNAs and mRNA. A-D) EMSAs of 5′ FAM-labeled homopolymeric RNA oligonucleotides with the indicated recombinant proteins. Due to the suspected net neutral charge of the RNA oligo•FMRP complex causing the RNP to not enter the gel, the intensity of unbound RNA was quantified and set relative to the protein storage buffer only sample (E). F) Fluorescence polarization of 5′ FAM-labeled homopolymeric RNA with the indicated recombinant protein. n=3 biological replicates. Data are shown as mean ± SD. n = 3 biological replicates. Comparisons were made using a two-tailed unpaired t test with Welch’s correction (p-values for each comparison are listed in **Supplemental Table S2**). G) EMSA of control reporter mRNA incubated with the indicated recombinant protein and stained with SYBR Green II.

It should be noted that we routinely did not see the RNP enter the gel for most of the homopolymeric RNAs tested, suggesting the high positive charge of the RGG+CTD neutralized the negative charge of the RNA oligo. The RGG box motif and CTD have theoretical isoelectric points of 12.1 and 10.0, respectively. Nevertheless, to be as consistent as possible across all the samples, unbound homopolymeric RNA was quantified (**Figure 3E**). It is possible that the U(G)_17_ formed intermolecular G4s, providing a high net negative charge to the complex that facilitated its entry into the gel if RGG+CTD was sub-stoichiometric. As a complementary approach to assess RNA binding that does not depend on an overall negative charge for electrophoresis, we measured the fluorescence polarization of each complex in solution.

Consistent with our EMSA results (**Figure 3A-E**), we observed an increase in polarization of all four FAM-labeled homopolymeric RNAs only when complexed with RGG+CTD (**Figure 3F**).

To further dissect how the ncRBD formed by the RGG+CTD elicits FMRP to inhibit translation, we tested whether recombinant NT-hFMRP WT and each mutant can bind reporter mRNA by EMSA (**Figure 3G**). Remarkably, the ability of WT and each mutant NT-hFMRP to bind reporter mRNA (**Figure 3G**) mirrors the observed translational repression with each mutant (**Figure 2**), except for ΔRGG+CTD. WT and I304N NT-hFMRP, as well as the RGG+CTD fusion, caused a complete gel shift (**Figure 3G**) and were translationally repressive (**Figure 2D, E, G**). RGG box alone and CTD alone did not cause a gel shift and were not robust translational repressors. Unique, ΔRGG+CTD did cause a complete gel shift (**Figure 3G**) but was not translationally repressive (**Figure 2F**). We rationalized that the functional KH domains in ΔRGG+CTD allowed mRNA binding since the ncRBD was absent in this mutant. Indeed, adding the I304N mutation in the KH2 domain to ΔRGG+CTD blocked reporter mRNA binding (**Figure 3G, Supplemental Figure S5**). Identical EMSA results were seen with G4-less mEGFP mRNA (**Supplemental Figure S6**). The decreased binding observed with ΔRGG+CTD when the I304N mutation was added suggests that the KH2 does compete with the ncRBD to bind mRNA. This is also observed in our translational repression assays with the I304N mutant having an ∼2-fold lower IC_50_ compared to WT **(Figure 2D, E**). Together, these data support that the RGG box motif and CTD forms a ncRBD that elicits broad RNA binding.

### Discrete regions of the ncRBD are required for translational repression and mRNA binding

We next asked if mRNA-binding and translational repression are elicited by the same or different regions of the ncRBD. Our pre-incubation data (**Figure 1F**) suggest that mRNA-binding is a prerequisite for FMRP translational repression. Thus, we postulated that if the region of the ncRBD that elicited mRNA-binding and translational repression were the same, we would see a mirrored loss in mRNA-binding and translational repression when mutated. The ncRBD of human FMRP is highly conserved among most vertebrates and is predicted to be largely flexible and disordered (**Supplemental Figure S7**), which hindered our ability to make refined mutations *a priori* as typically is done for canonical RBDs. We generated a series of large and more refined serial deletions of the CTD from the C-terminal end (**Figure 4A**) and identified a sharp decline in translational repression between the RGG+CTD Δ54 and RGG+CTD Δ55 mutants (**Figure 4B**). This transition point between deleting 54 or 55 amino acids from the C-terminus of the ncRBD was also present when tested in the NT-hFMRP isoform that harbors the other canonical RBDs (**Supplemental Figure S8**). NT-hFMRP Δ54 had an IC_50_ of ∼0.88 μM (**Figure 4C**) and NT-hFMRP Δ55 had an IC_50_ of ∼2.29 μM (**Figure 4D**). We attempted to introduce the I304N KH2 domain mutation to offset any altered binding of the truncated C-terminus, but these mutants were insoluble. These data demonstrated that RGG+CTD Δ54 region of the ncRBD represents the minimal repressive element of human FMRP.

**Figure 4.**
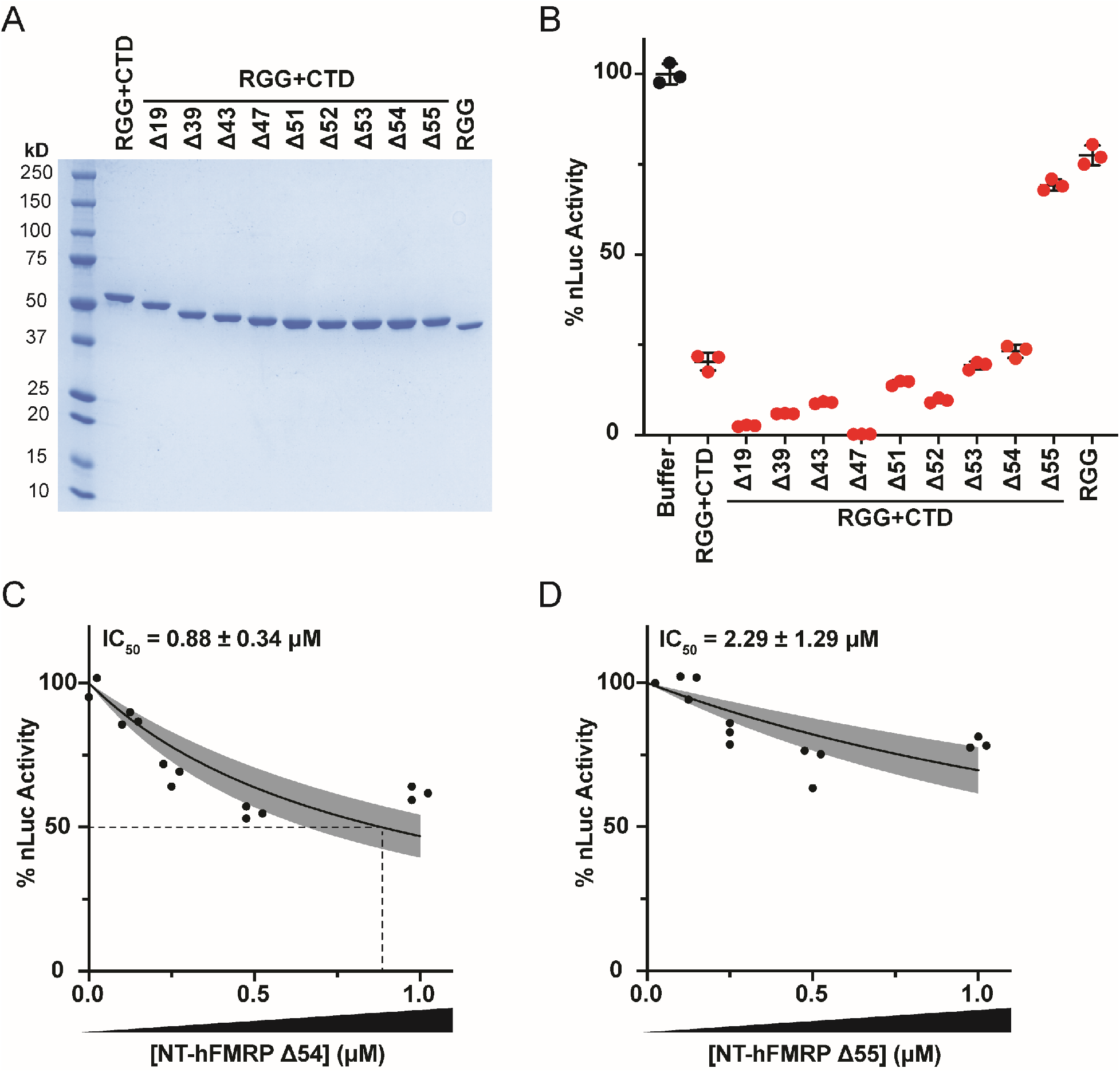
The RGG+CTD Δ54 is required for translational repression. A) Coomassie stain of recombinant proteins with serial truncations to the CTD. The number of amino acids truncated from the CTD is indicated. B) *In vitro* translation of nLuc mRNA with 1 μM CTD truncated recombinant RGG+CTD proteins and protein storage buffer as a negative control. Data are shown as mean ± SD. n = 3 biological replicates. All recombinant protein tested was statistically significant compared to buffer negative control using a two-tailed unpaired t test with Welch’s correction (each comparison had p < 0.01). C, D) *In vitro* translation of nLuc mRNA with a titration of recombinant NT-hFMRP Δ54 (C) and NT-hFMRP Δ55 (D). n=3 biological replicates. A non-linear regression was used to calculate the IC_50_ value for each truncation and is shown as the line with the 95% confidence interval (CI) included as a watermark. The IC_50_ is reported ± 95% CI.

If the change in translational repression between RGG+CTD Δ54 and RGG+CTD Δ55 was due to altered mRNA-binding ability, we would predict a similar drastic change in mRNA binding. However, both RGG+CTD Δ54 and RGG+CTD Δ55 had nearly identical ability to cause a robust gel shift of reporter mRNA by EMSA (**Figure 5A, B**). Further deletional analysis and serial single amino acid truncations (RGG+CTD Δ60, Δ61, Δ62, and Δ63) identified RGG+CTD Δ62 as the key region of the ncRBD for robust mRNA binding (**Figure 5A, B**). Together, these data suggest that the regions of the ncRBD that are responsible for mRNA binding and translational repression overlap but are not identical (**Figure 5C**).

**Figure 5.**
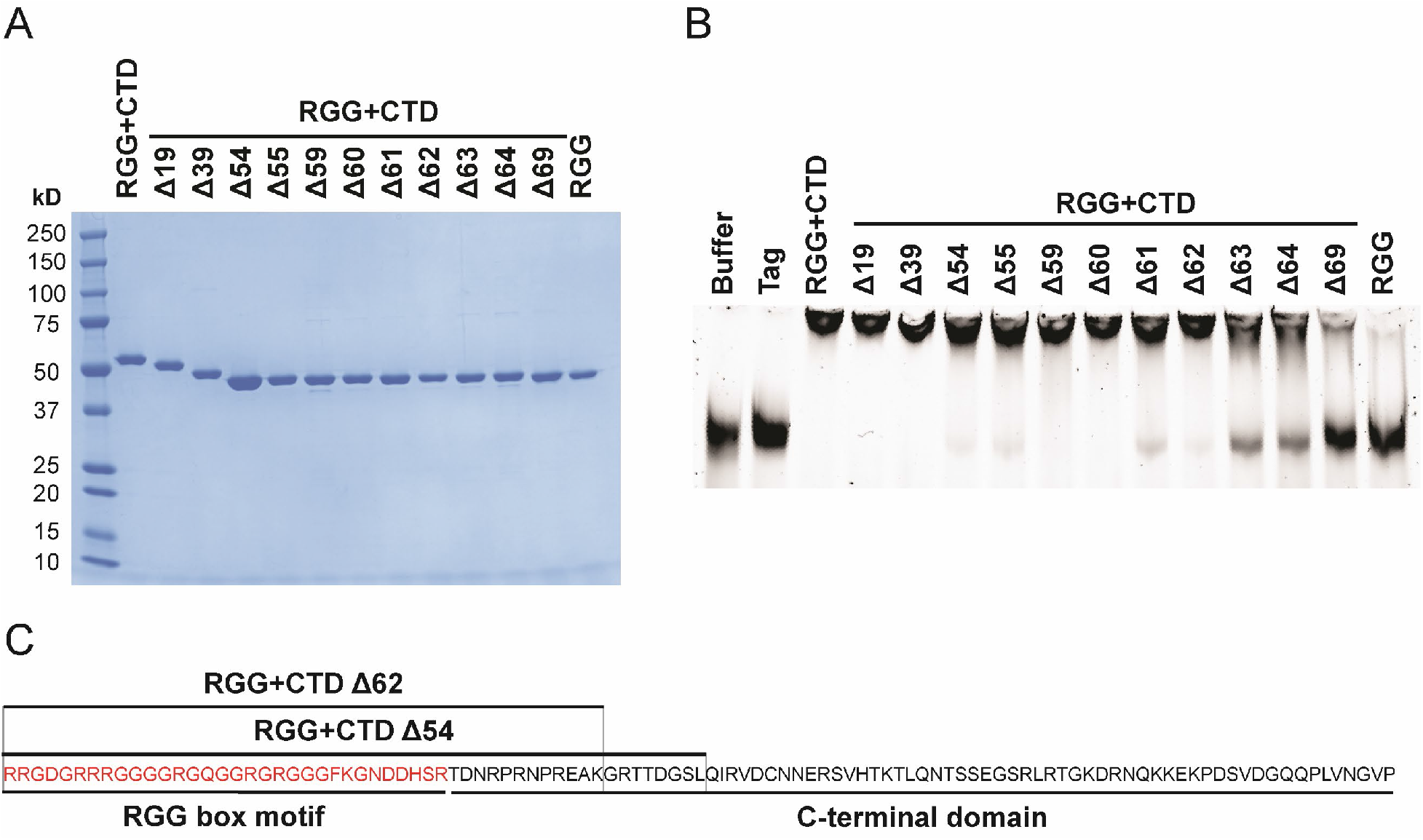
The RGG+CTD Δ62 is the key region of the ncRBD required for robust mRNA-binding. A) Coomassie stain of recombinant proteins with C-terminal truncations to the ncRBD (RGG+CTD). The number of amino acids truncated from the CTD is indicated. B) EMSA of control reporter mRNA incubated with recombinant protein and stained with SYBR Green II. C) Amino acid composition of overlapping regions of the ncRBD that are required for mRNA-binding (RGG+CTD Δ62) and translational repression (RGG+CTD Δ54).

### FMRP inhibits translation post-initiation when bound to mRNA via the ncRBD

FMRP bound to mRNA could directly inhibit translation by either blocking the scanning pre-initiation complex (PIC) in the 5′ UTR or stalling the elongating ribosome in the CDS. In general, scanning PICs are more susceptible and sensitive to obstacles in their path (i.e., RNA structure or bound RBPs) than elongating ribosomes. Most mRNA-bound FMRP is mapped to the CDS of mRNAs *in vivo*, not to the 5′ UTR (21). However, in our assays, it is possible that a portion of recombinant FMRP is bound to the 5′ UTR and is simply blocking the scanning PIC. To confirm that FMRP inhibits translation post-initiation, consistent with previous reports that FMRP slows or stalls elongating ribosomes (21-25), we used the following three distinct strategies.

First, we rationalized that if 5′ UTR-bound recombinant FMRP was blocking scanning PICs, extending the 5′ UTR length would enhance repression. In this case, extending the 5′ UTR would provide FMRP increased opportunity to bind the 5′ UTR instead of the CDS. To achieve this, we mutated the control reporter that harbors the 50 nt human β-globin 5′ UTR and extended the 5′ UTR three-fold by inserting two additional β-globin 5′ UTR sequences (resulting in a 150 nt 5′ UTR). Nevertheless, FMRP inhibited translation of the control and long 5′ UTR reporter mRNAs to similar extents (**Figure 6A, Supplemental Figure S2C**). These data suggest that the predominant mechanism by which FMRP inhibits translation in our assays is not by inhibiting scanning PICs.

**Figure 6.**
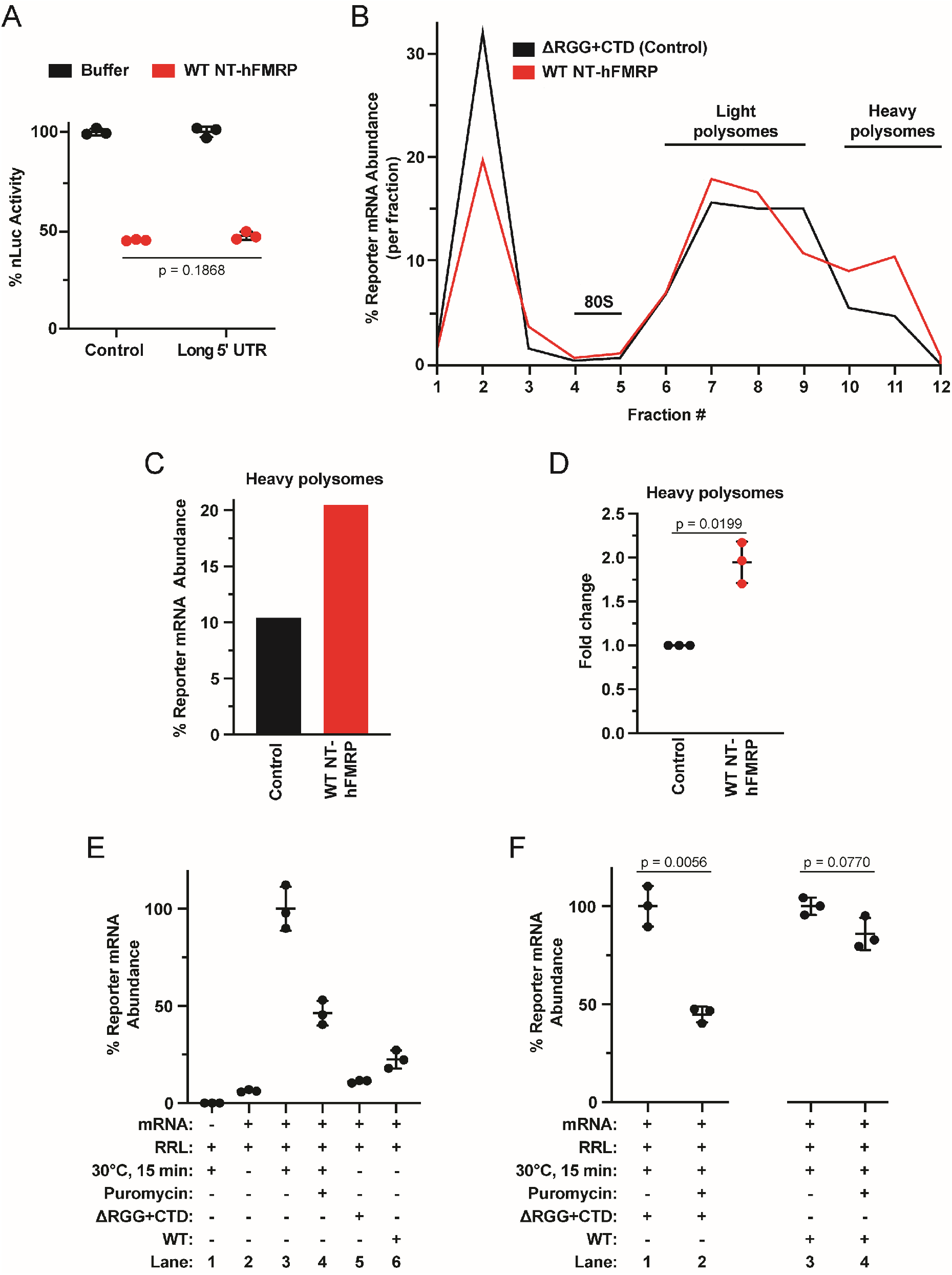
Human FMRP inhibits translation post-initiation when binding mRNA via the ncRBD. A) *In vitro* translation of the control reporter mRNA that harbors the 50 nt human β-globin 5′ UTR and a long 5′ UTR reporter mRNA that harbors three β-globin 5′ UTRs in tandem (150 nt total). mRNPs were formed with protein storage buffer or 1 μM WT NT-hFMRP. Data are shown as mean ± SD. n = 3 biological replicates. Comparisons were made using a two-tailed unpaired t test with Welch’s correction. B) Distribution of nLuc reporter mRNA across sucrose gradients to assess polysome formation when pre-incubated with 1 μM ΔRGG+CTD (Control) or 1 μM WT NT-hFMRP. Abundance of reporter mRNA in each gradient fraction was determined by RT-qPCR. C) Cumulative nLuc abundance in heavy polysomes in fractions 10-12 from 1 μM ΔRGG+CTD (Control) and 1 μM WT NT-hFMRP samples. D) Fold change of nLuc mRNA abundance in heavy polysomes. Data are shown as mean ± SD. n = 3 biological replicates. Comparisons were made using a two-tailed unpaired t test with Welch’s correction. E) Relative quantification of nLuc reporter mRNA pelleted through a 35% (w/v) sucrose cushion after a low-speed centrifugation (see **Supplemental Figure S10**). Lane 1 is a negative control lacking nLuc mRNA. Lane 2 is a negative control containing mRNA in RRL but not incubated at 30°C to start translation. Lane 3 is nLuc mRNA in RRL translated for 15 min at 30°C. Lane 4 is nLuc mRNA in RRL translated for 15 min at 30°C and then incubated with 0.1 mM puromycin (final) for 30 min at 30°C. Lanes 5 and 6 are negative control untranslated reactions of nLuc•FMRP mRNPs formed with 1 μM ΔRGG+CTD (Control) and 1 μM WT NT-hFMRP, respectively, in RRL kept on ice demonstrating poor pelleting without active translation through the low-speed sucrose cushion as described in the Materials and Methods and outlined in **Supplemental Figure S10**). Data are shown as mean ± SD. n = 3 biological replicates. Comparisons were made using a two-tailed unpaired t test with Welch’s correction. F) Relative quantification of nLuc reporter mRNA pelleted through a 35% (w/v) sucrose cushion after a low-speed centrifugation. nLuc•ΔRGG+CTD (Control) and nLuc•WT NT-hFMRP mRNPs were translated and treated with 0.1 mM puromycin (final) before being overlayed on the cushion and low-speed centrifugation. Final concentration of recombinant protein was 1 μM. Data are shown as mean ± SD. n = 3 biological replicates. Comparisons were made using a two-tailed unpaired t test with Welch’s correction.

Second, we used sucrose density gradient ultracentrifugation to confirm the expected distribution of mRNAs with stalled elongating ribosomes. For example, elongation inhibitors (e.g., cycloheximide or emetine) cause an increase in polysomes as they do not prevent initiation, but slow down and stabilize elongating ribosomes on mRNAs (38). This is typically seen as an increase in polysome signal in the heavier fractions of sucrose gradients. If FMRP inhibits translation post-initiation when bound to mRNA via its ncRBD, we would predict that polysomes accumulate on inhibited transcripts only when FMRP harbored the RGG+CTD ncRBD. To test this prediction, we used the ΔRGG+CTD mutant as a negative control as it does not inhibit translation (**Figure 2F**) but still binds the reporter mRNA (**Figure 3G**) in our assays.

After *in vitro* translation and ultracentrifugation, we quantified nLuc reporter mRNA abundance in each fraction. The monosome peak (which is primarily inactive 80S ribosomes native to RRL (39)) was routinely found in fractions 4 and 5 (**Supplemental Figure S9A**), indicating that polysomes would sediment in fractions 6 through 12. Compared to the negative control that lacks the translationally repressive ncRBD, WT NT-hFMRP increased the abundance of nLuc mRNA in the heavy polysomes (**Figure 6B-D, Supplemental Figure S9B-E**). Specifically, nLuc mRNA abundance in the heavy polysomes in fractions 10-12 increased ∼2-fold (**Figure 6C, D, Supplemental Figure S9C, E**). This increase of reporter mRNA at the bottom of the gradient with the heavy polysomes is consistent with accumulation of slowed and stalled ribosomes.

Lastly, we tested the ability of NT-hFMRP to generate ribosome complexes on reporter mRNPs that are resistant to puromycin. FMRP was previously identified by immunogold labeling and electron microscopy to be bound to ribosomes within puromycin-resistant polysomes from mouse brain (21). Puromycin is an aminonucleoside antibiotic and acyl tRNA analog that is incorporated into nascent polypeptides and results in ribosomes releasing both the nascent polypeptide and bound mRNA. Puromycin sensitivity is specific for actively elongating 80S ribosomes that are in an unrotated (classic) state with an empty A site (e.g., during the decoding step of elongation). Slowly elongating ribosomes that have an occupied A site, that are stalled in the rotated (or hybrid) state, or that are inhibited during translocation are resistant to puromycin and stay bound to mRNA. We predicted that NT-hFMRP would stall and slow elongating ribosomes and elicit puromycin-resistance only when harboring the ncRBD. We optimized a low-speed sucrose cushion protocol to be selective for mRNAs only bound by ribosomes to be recovered, allowing us to assay puromycin-resistant ribosomes on nLuc•FMRP mRNPs (**Supplemental Figure S10**). A series of controls demonstrates that nLuc mRNA is detected in the ribosome pellet only after being translated and this detection is strongly prevented when completed translation reactions are treated with 0.1 mM puromycin prior to low-speed centrifugation over a 35% sucrose cushion (**Figure 6E, Lanes 1-4**). Only minor amounts of nLuc mRNA were recovered from untranslated reactions with nLuc•NT-FMRP mRNPs (ΔRGG+CTD or WT) in RRL that were kept on ice (**Figure 6E, Lanes 5 & 6**). In agreement with our prediction that only NT-hFMRP harboring the ncRBD would elicit puromycin-resistance, ribosome-bound nLuc•NT-hFMRP ΔRGG+CTD mRNP, but not nLuc•NT-hFMRP WT mRNP, was sensitive to puromycin (**Figure 6F, Lanes 1 & 2**). We were unable to detect a change in pelleted nLuc from the translated nLuc•NT-hFMRP WT mRNP when treated with puromycin (**Figure 6F, Lanes 3 & 4**). In total, these data support that FMRP uses a ncRBD formed by the RGG box motif and CTD to bind mRNA independent of mRNA G4s resulting in ribosome stalling and translational repression (**Figure 7**).

**Figure 7.**
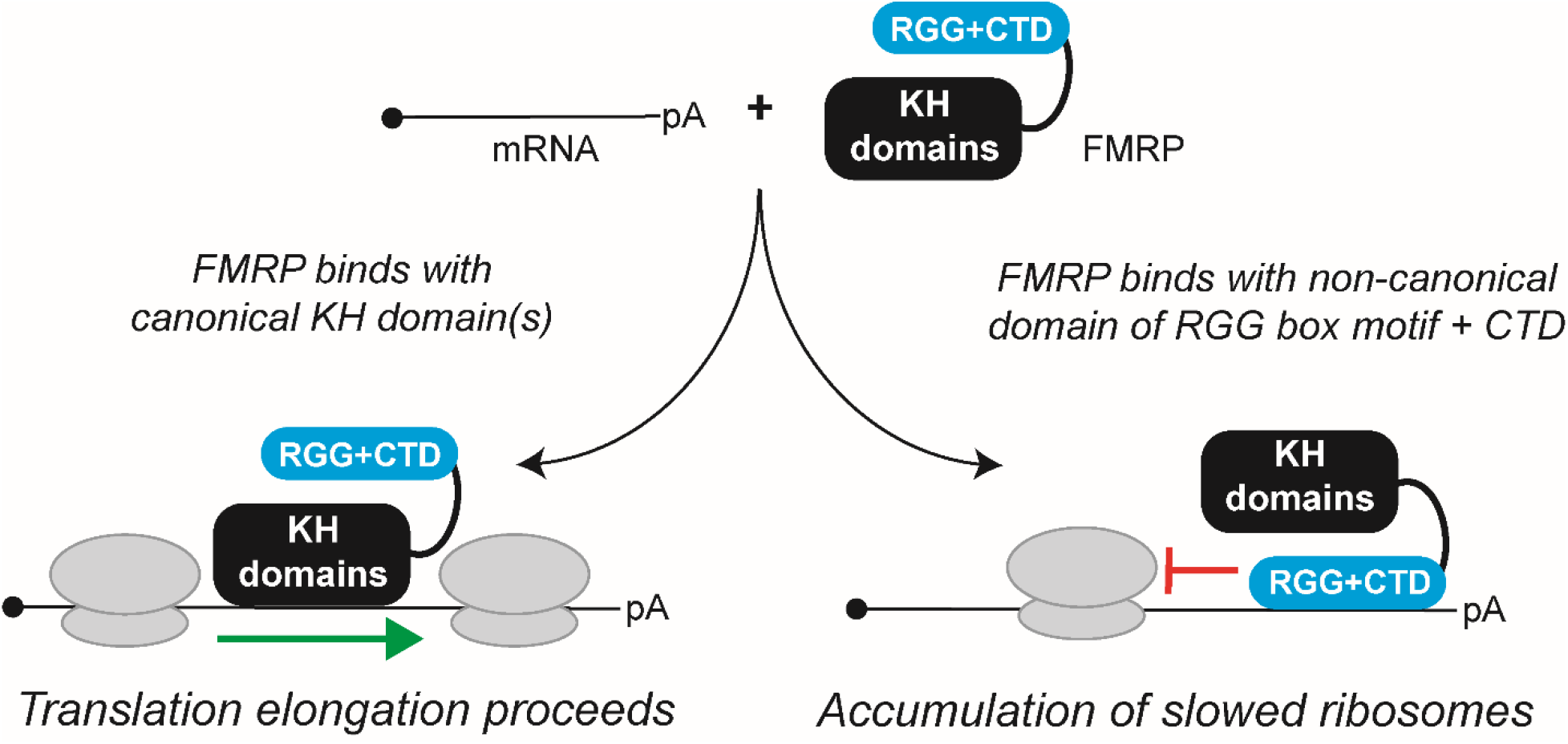
Model of FMRP-mediated translational repression via a ncRBD. FMRP can bind mRNA through canonical and non-canonical RBDs. When bound to mRNA through its canonical KH domain(s), FMRP does not robustly inhibit translation. The RGG box motif and CTD of FMRP together form a ncRBD that allows FMRP to bind multiple mRNAs and subsequently inhibit translation to cause accumulation of slowed/stalled ribosomes.

## DISCUSSION

The complexity of molecular and cellular phenotypes seen in FMRP-null neurons and model systems has led to much debate surrounding how FMRP targets transcripts and what functions it may possess once bound to mRNA (40). Most early data support FMRP as a translational repressor (21-25), but recent work in *Drosophila* and mouse models has provided compelling evidence that FMRP may in fact also act a translational activator (26-28). It remains largely unclear which RNA motifs or structures FMRP targets as few common enriched sequences were found across multiple transcriptome-wide FMRP binding studies (reviewed in 40). A single consensus sequence for FMRP may be an oversimplification since FMRP has multiple RBDs and motifs.

Here we refine the RNA-binding element that contains the RGG box motif and provide evidence that FMRP inhibits translation after binding mRNA via the ncRBD comprised of the RGG box motif and the CTD. The ncRBD is essential and sufficient for repression (24,41) (**Figure 2**) and required for FMRP to bind mRNA to inhibit translation (**Figures 2 & 3**). Scanning deletion analysis of the ncRBD identified that the residues required for mRNA binding reside within the region critical for translational repression (**Figures 4 & 5**). We and others (24,41) have found the KH1 and KH2 domains dispensable for translational repression, yet cryo-EM determination of *Drosophila* FMRP•80S ribosome complexes shows that KH1 and KH2 domains partially overlap with where the anticodon stem of a bound P-site tRNA would be located. In this structure, the RGG+CTD region of FMRP is near the A site and leading edge of the ribosome (22). Chen *et al*. proposed that the RGG box motif binds to the mRNA via G4s within the CDS and that the KH domains bind directly to the ribosome near the A site to sterically hinder delivery of charged tRNAs (22). Early studies identified FMRP-bound polysomes from mouse brain lysate as puromycin-resistant, a characteristic we demonstrate is reproduced in our translation assays (**Figure 6E, F**). Puromycin is a substrate for peptidyl transferase when ribosomes have an empty A site and are in an unrotated state during the decoding step of elongation. Puromycin-resistant FMRP-stalled ribosomes are thus thought to be inhibited during translocation (21). However, the fly FMRP•80S ribosome structure was solved using empty 80S ribosomes in a unrotated state unbound to mRNA. To our knowledge, our report is the first to provide *in vitro* biochemical data that demonstrate FMRP creates puromycin-resistant ribosome complexes on mRNA, but in agreement with what was previously shown in brain tissue lysates (21). The abundance of inhibited transcripts on heavier polysomes confirms translational repression is not due to decreased initiation (**Figure 6B-D; Supplemental Figure S9**). Our approach reported here provides valuable samples and tools to enrich for these complexes, which is not easily achievable from brain tissue lysates. Future structural determination of FMRP-inhibited ribosomes on mRNA will be critical for fully understanding FMRP-mediated translational repression.

Although the RGG box motif within the FMRP ncRBD has high affinity to the G4-containing Sc1 RNA (17), this interaction is not required for translational repression, nor does it provide enhanced translational repression (**Figure 1, Supplemental Figure S2**). In fact, structural determinations of the RGG box motif and Sc1 RNA complex show that the RGG box motif actually binds to the major groove of the duplexed RNA region of the Sc1 RNA and not to the G4 itself (32,33). Moreover, G4s are enriched in the UTRs of mRNAs but not in the CDS (42), while FMRP predominately binds to the CDS but not to the UTRs (21). If FMRP is recruited to the CDS via RGG box-G4 interaction, one would predict the distributions of FMRP binding events and G4s to be positively correlated, but they are instead negatively correlated. Recent ribosome profiling studies that incorporated RNA G4 prediction analyses show that transcripts that are derepressed in FMRP knockout cell lines are not enriched for RNA G4s (43). Goering *et al*. conclude that RNA G4s were instead correlated with FMRP-mediated mRNA localization, not translational repression. A more recent study from Darnell and colleagues found that G4s were not enriched in dendritic FMRP targets in CA1 pyramidal neurons (44).

Consistent with our finding that repression occurs independent of RNA G4s in the CDS (**Figure 1**), the ncRBD has a broad ability to bind RNA (**Figure 3, Supplemental Figure S6**). In further support of this G4-independent mode of translational repression by FMRP, two other reports have identified that FMRP binds to a stem loop-containing RNA (devoid of a G4) to inhibit translation. First, Maurin *et al*. identified mammalian FMRP inhibited expression of a reporter mRNA that contained the SL1 and SL2 stem-loops when inserted into the open reading frame but not either UTR. Which RNA-binding domain (KH1, KH2, or ncRBD) was responsible for this inhibition was not reported (45). Second, while this report was in preparation, Edwards *et al*. reported that the RGG+CTD region bound to a stem-loop in a short peptide reporter mRNA to inhibit translation (41). For both reports, further studies are warranted to determine if the stem-loops in the open reading frame cause FMRP to inhibit initiation and/or elongation. Nevertheless, the stem-loop sequences identified by Maurin *et al*. and Edwards *et al*. are absent in our nLuc reporter mRNA. Our finding that the ncRBD can bind all four homopolymeric RNAs and multiple G4-less reporter mRNAs suggests that FMRP binding is more promiscuous than previously appreciated. Future iCLIP studies in cells expressing the ncRBD alone will be important to determine, if any, sequence or RNA secondary structure preferences for binding and subsequent translational repression.

The structure of the ncRBD in FMRP bound or unbound to RNA is not known, but is predicted to be disordered and flexible (**Supplemental Figure S7**). A small fragment of the RGG box motif was used in determining the RGG box•Sc1 RNA complex and was found to be highly flexible. Recent reports have also shown that small repetitive regions (46) (e.g., poly-K patches, poly-R patches, and YGG box motifs) and other larger flexible regions have previously undefined RNA-binding abilities. For example, the disordered and flexible N-terminal half of *C. elegans* MEG-3 elicits RNA-binding (yet, does not cause phase transition that is commonly reported for such disordered regions) (47). Additionally, Xu *et al*. recently characterized human nuclear hormone receptor estrogen receptor α (ERα) as an RNA-binding protein that contains a functional arginine-rich RBD with binding preference to A-U rich sequences in 3′ UTRs (48) to control post-transcriptional regulation. Further assessment in AlphaFold and IUPred predicts the arginine-rich RBD in ERα is centered in a large highly flexible region (data not shown), similar to the FMRP ncRBD we define in this report. Our understanding of how flexible RNA-binding domains mechanistically associate with RNA is increasing but whether they contain sequence specificity is still unclear.

FMRP also facilitates the transport of dendritic mRNAs to synapses *in vivo* and this transport can be dependent on its ability to bind mRNA (43,49). Given that the translationally repressive ncRBD binds mRNA more promiscuously than previously appreciated (**Figure 3, Supplemental Figure S6**) and the lack of a shared consensus sequence that has been found throughout multiple transcriptome-wide FMRP binding screens (40), the data supports a model where FMRP could regulate translation of dendritic mRNAs primarily based on local concentration and not mRNA sequence. Since FMRP and dendritic mRNAs are transported to the synapses, often together, both are found at relatively high concentrations compared to other translationally active sites of neurons. Future studies of directly shuttling FMRP from one region of neuron to another and monitoring its ability to repress translation of proximal transcripts would prove beneficial in further defining a concentration-dependent model.

## ACKNOWLEDGEMENTS

Experiments were conceived by MRS and MGK and were performed by MRS, JEW, ELY, and MGK. The manuscript was written by MRS and MGK with input from JEW and ELY. We thank members of the Kearse lab and Aaron Goldstrohm for critically reading the manuscript. We also thank Brian Scarpitti for his help with image analyses and Christine Daugherty at the OSUCCC GSR.

## SUPPLEMENTAL DATA

**Supplemental Table S1.**
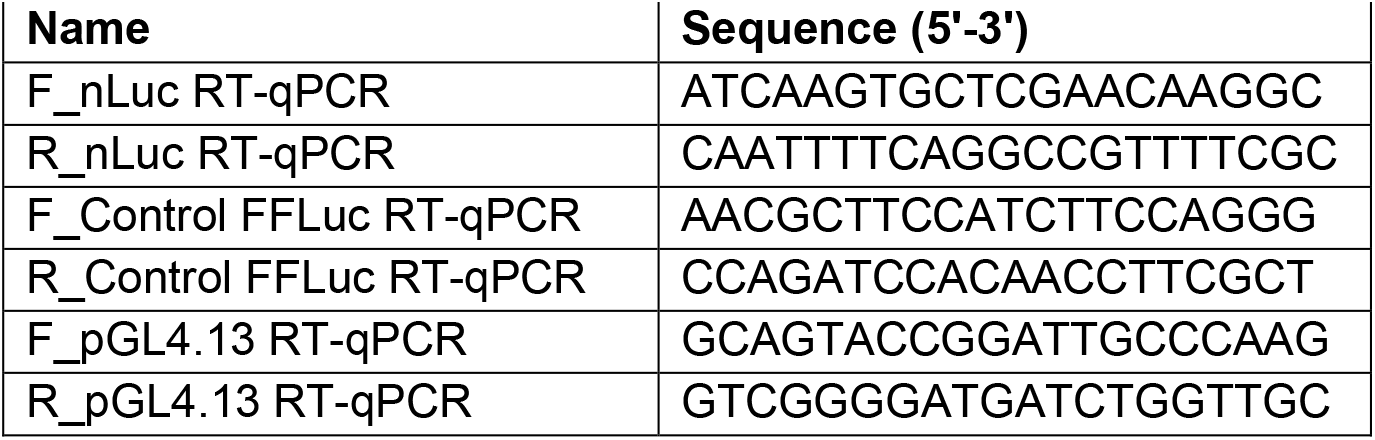
Oligonucleotides used in this study.

**Supplemental Table S2.**
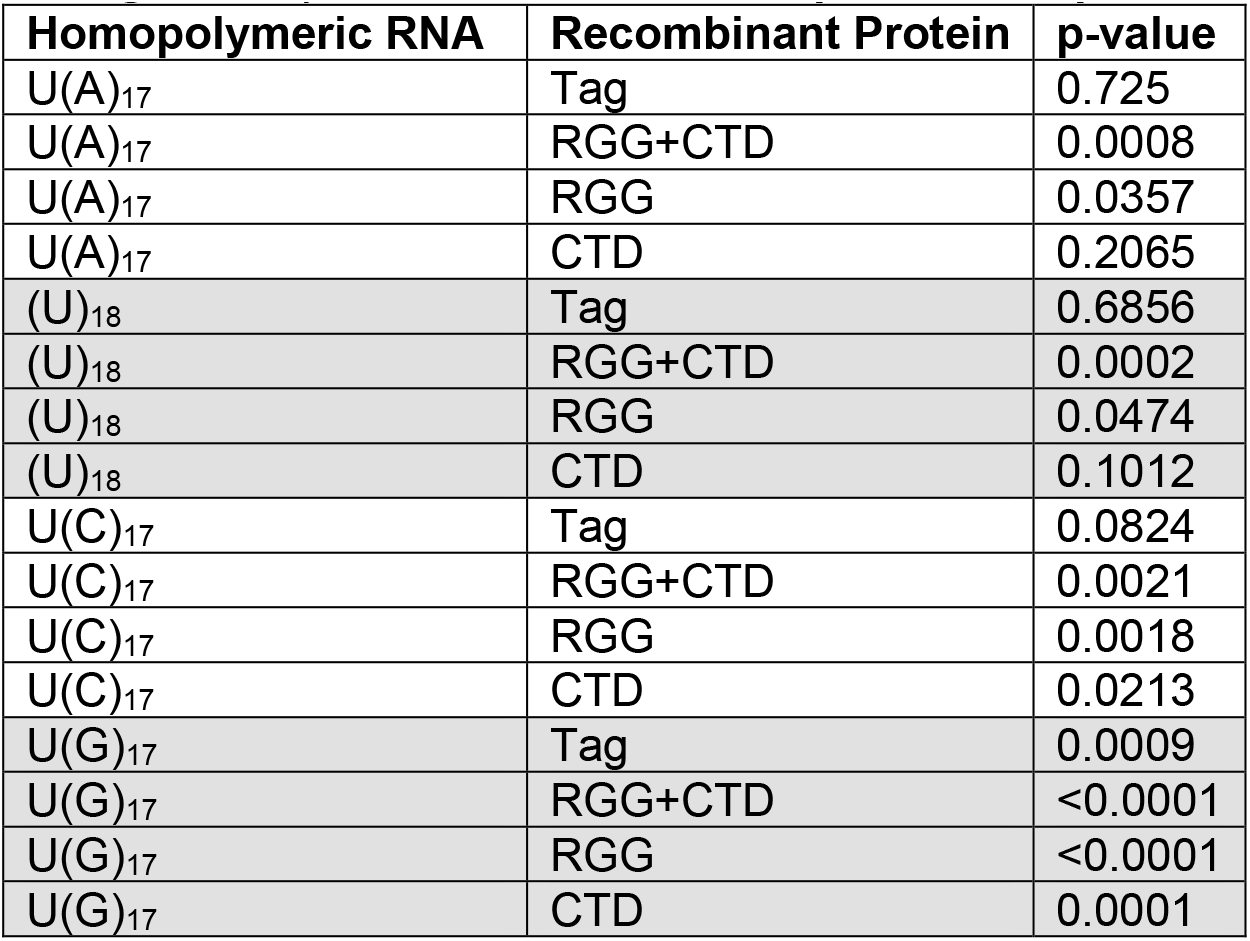
P-values of two-tailed unpaired t tests with Welch’s correction for Figure 3F (indicated recombinant protein compared to Buffer samples).

| Homopolymeric RNA | Recombinant Protein | p-value |
| --- | --- | --- |
| U(A)<sub>17</sub> | Tag | 0.725 |
| U(A)<sub>17</sub> | RGG+CTD | 0.0008 |
| U(A)<sub>17</sub> | RGG | 0.0357 |
| U(A)<sub>17</sub> | CTD | 0.2065 |
| (U)<sub>18</sub> | Tag | 0.6856 |
| (U)<sub>18</sub> | RGG+CTD | 0.0002 |
| (U)<sub>18</sub> | RGG | 0.0474 |
| (U)<sub>18</sub> | CTD | 0.1012 |
| U(C)<sub>17</sub> | Tag | 0.0824 |
| U(C)<sub>17</sub> | RGG+CTD | 0.0021 |
| U(C)<sub>17</sub> | RGG | 0.0018 |
| U(C)<sub>17</sub> | CTD | 0.0213 |
| U(G)<sub>17</sub> | Tag | 0.0009 |
| U(G)<sub>17</sub> | RGG+CTD | <0.0001 |
| U(G)<sub>17</sub> | RGG | <0.0001 |
| U(G)<sub>17</sub> | CTD | 0.0001 |

**Supplemental Figure S1.**
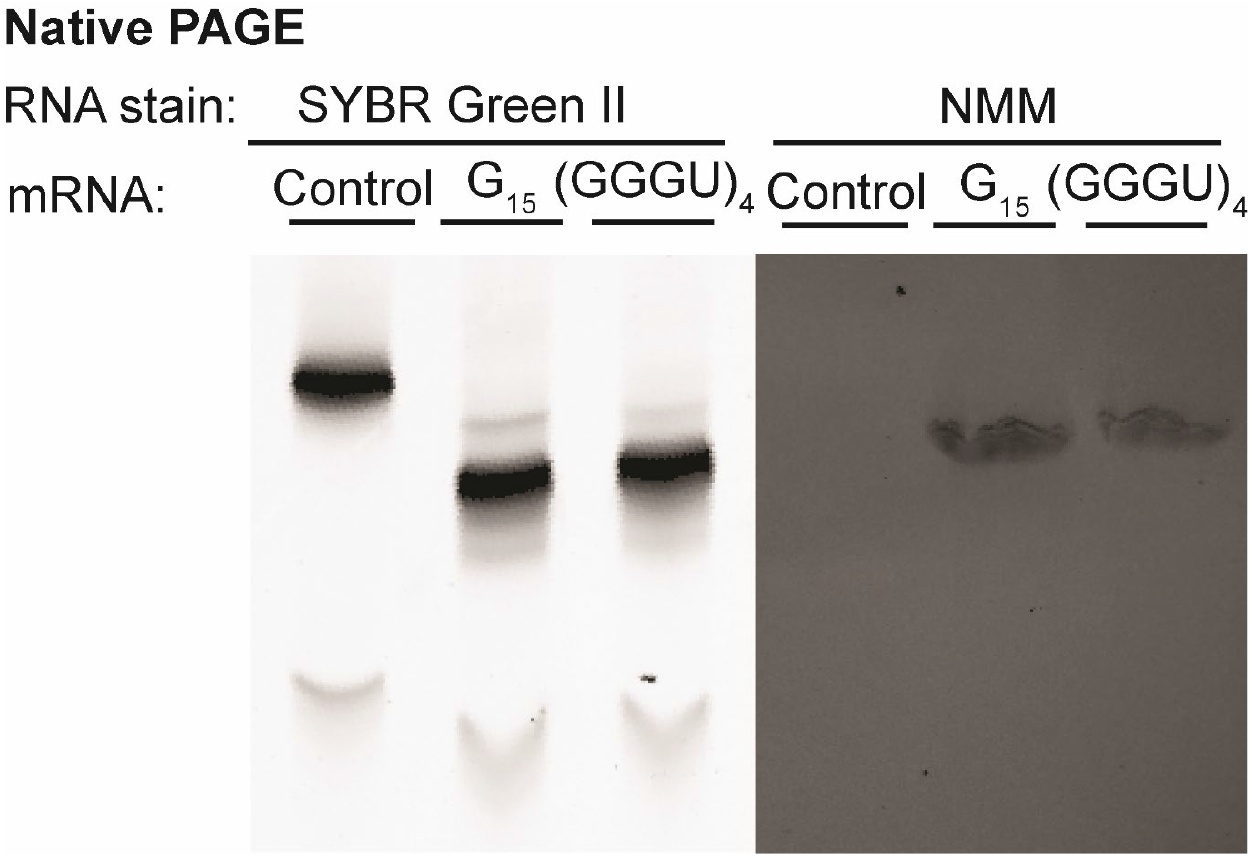
G_15_ and (GGGU)_4_ reporter mRNA, but not control reporter mRNA, harbor G-quadruplexes. Native PAGE of control, G_15_, and (GGGU)_4_ reporters stained for total RNA with SYBR Green II or for G4 structures with NMM.

**Supplemental Figure S2.**
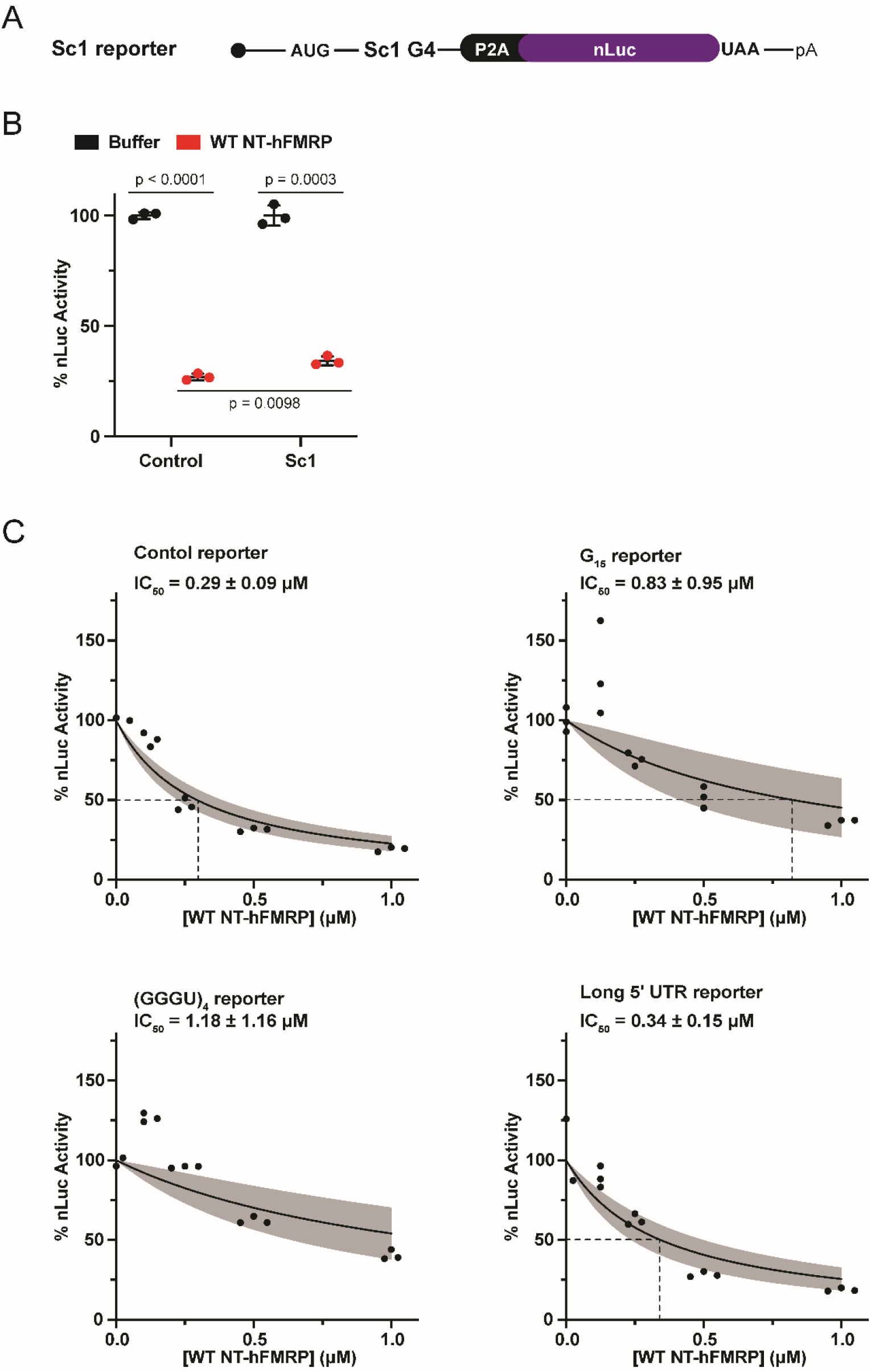
The Sc1 G4 does not enhance translational repression by FMRP. A) Schematic of custom nLuc reporter harboring the Sc1 G4 in the coding sequence. A P2A ribosome skipping motif was included immediately upstream of the nLuc coding sequence to ensure equal nLuc function between reporters. B) *In vitro* translation of Sc1 reporter Mrna with protein storage buffer or WT NT-hFMRP. Data are shown as mean ± SD. n = 3 biological replicates. Comparisons were made using an unpaired t test with Welch’s correction. C) *In vitro* translation of control, G_15_, (GGGU)_4_, and long 5′ UTR nLuc mRNA with a titration of recombinant wildtype NT-hFMRP. IC_50_ values were determined for control nLuc (0.29 ± 0.09 μM), G_15_ nLuc (0.83 ± 0.95 μM), (GGGU)_4_ nLuc (1.18 ± 1.16 μM), and long 5′ UTR nLuc (0.34 ± 0.15 μM) reporter mRNAs. n=3 biological replicates. A non-linear regression was used to calculate the IC_50_ and is shown as the line with the 95% confidence interval (CI) included as a watermark. The IC_50_ is reported ± 95% CI.

**Supplemental Figure S3.**
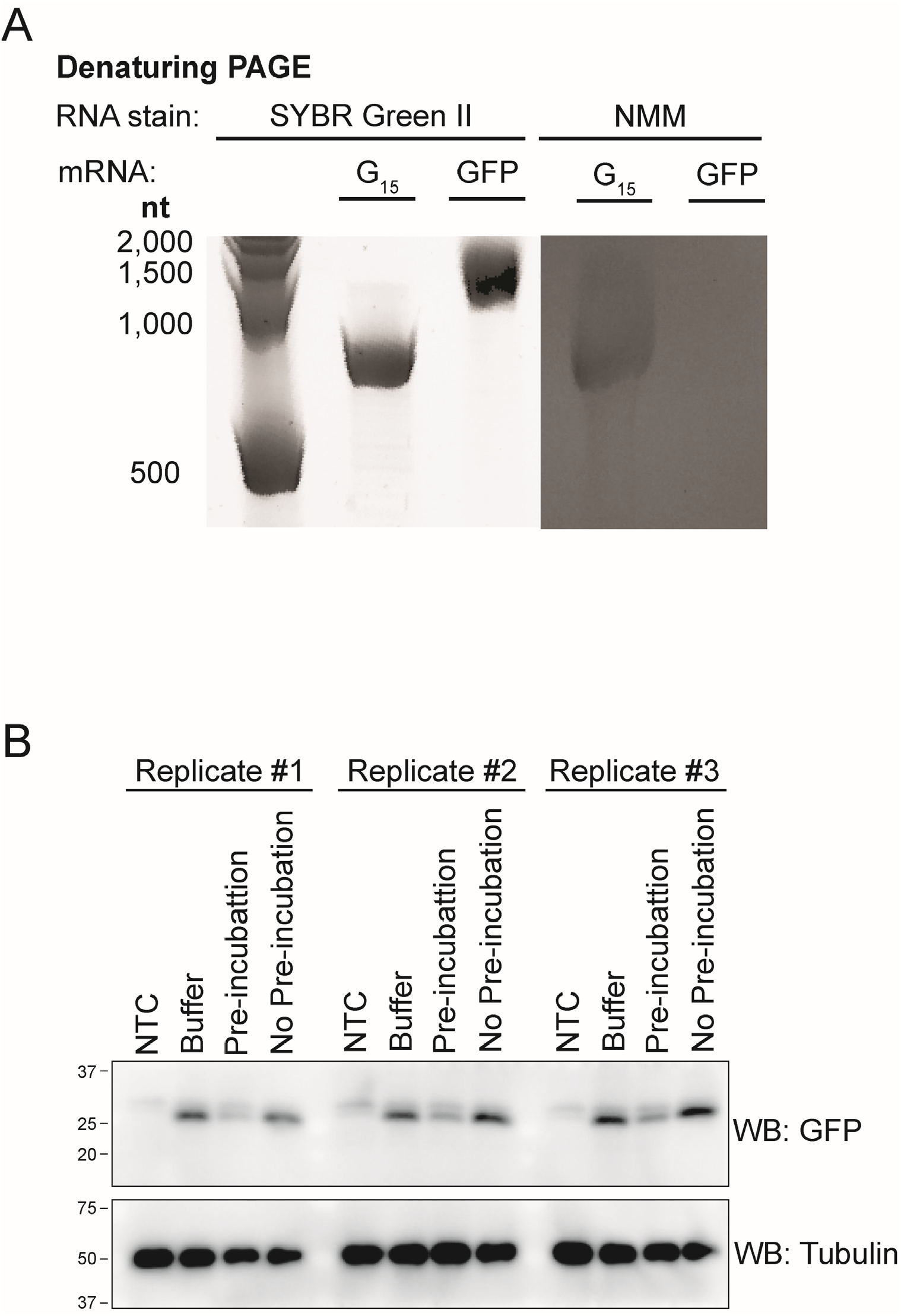
NT-hFMRP inhibits translation of G4-less mEGFP mRNA. A) Denaturing PAGE of G_15_ nLuc reporter and mEGFP mRNAs stained for total RNA with SYBR Green II or for G4 structures with NMM. B) Anti-GFP Western blot of *in vitro* translation reactions of G4-less mEGFP reporter mRNA with an no template control (NTC), protein storage buffer as an additional negative control, with 1 μM WT NT-hFMRP and G4-less mEGFP mRNA pre-incubated together, and with 1 μM WT NT-hFMRP and G4-less mEGFP without a pre-incubation step. Tubulin was used as a loading control.

**Supplemental Figure S4.**
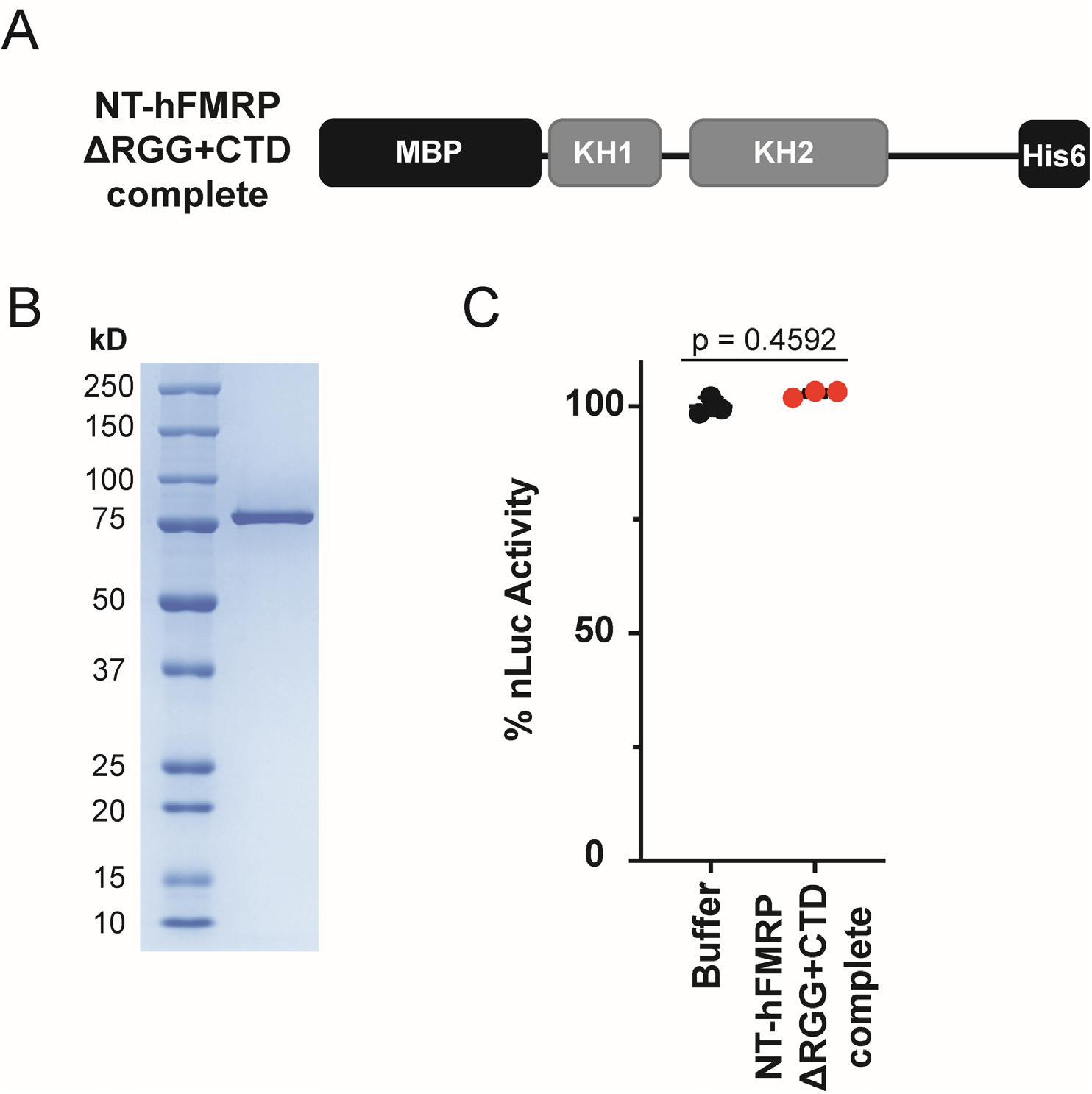
NT-hFMRP ΔRGG+CTD complete does not inhibit translation. A) Schematic of MBP- and His6-tagged NT-hFMRP ΔRGG+CTD complete. B) Coomassie stain of recombinant NT-hFMRP ΔRGG+CTD complete. C) *In vitro* translation of nLuc control mRNA with protein storage buffer or NT-hFMRP ΔRGG+CTD complete. Data are shown as mean ± SD. n = 3 biological replicates. Comparisons were made using an unpaired t test with Welch’s correction.

**Supplemental Figure S5.**
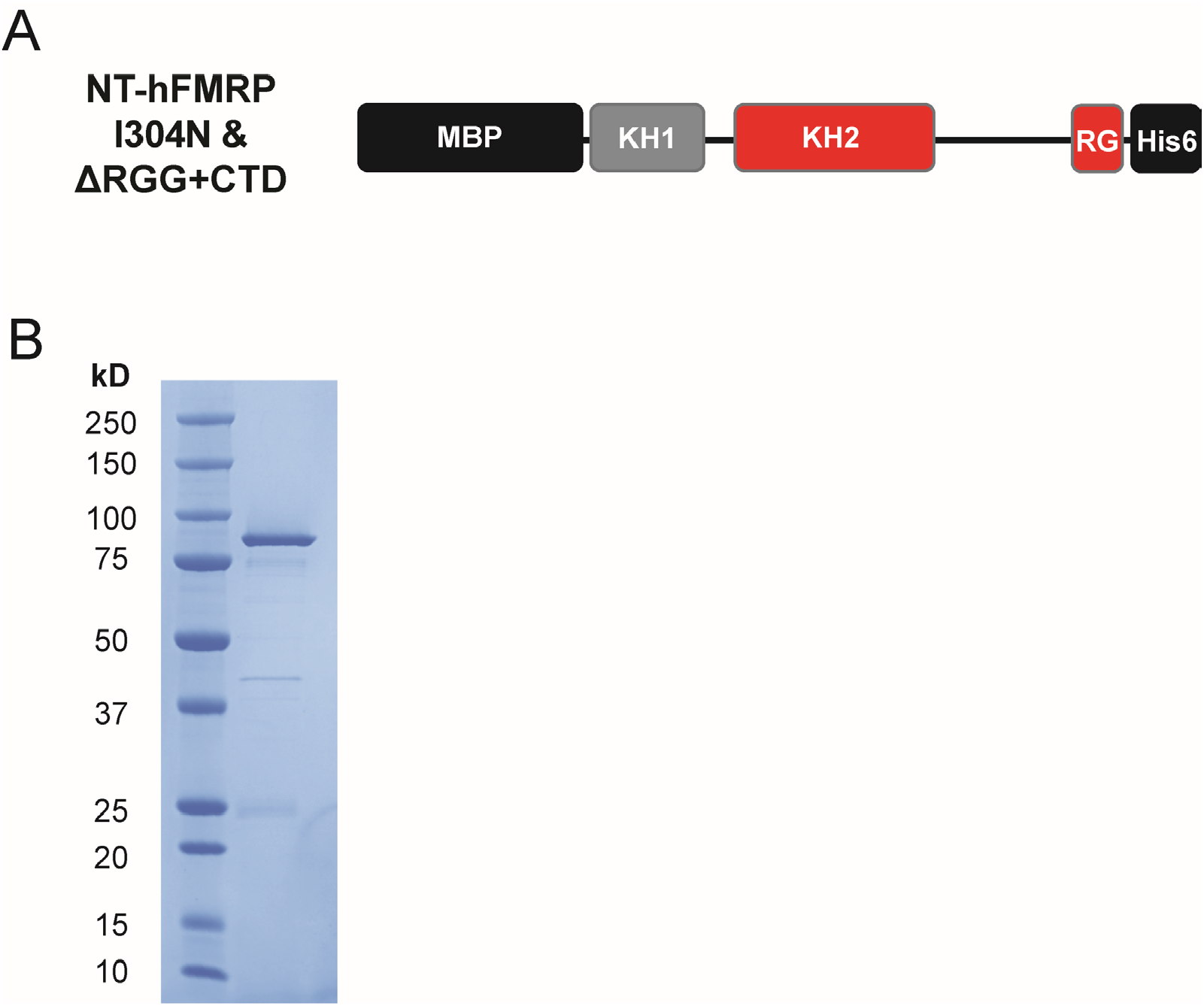
NT-hFMRP I304N & ΔRGG+CTD double mutant. A) Schematic of MBP- and His6-tagged NT-hFMRP I304N & ΔRGG+CTD. Mutated/truncated domains are highlighted in red. B) Coomassie stain of recombinant NT-hFMRP I304N & ΔRGG+CTD.

**Supplemental Figure S6.**
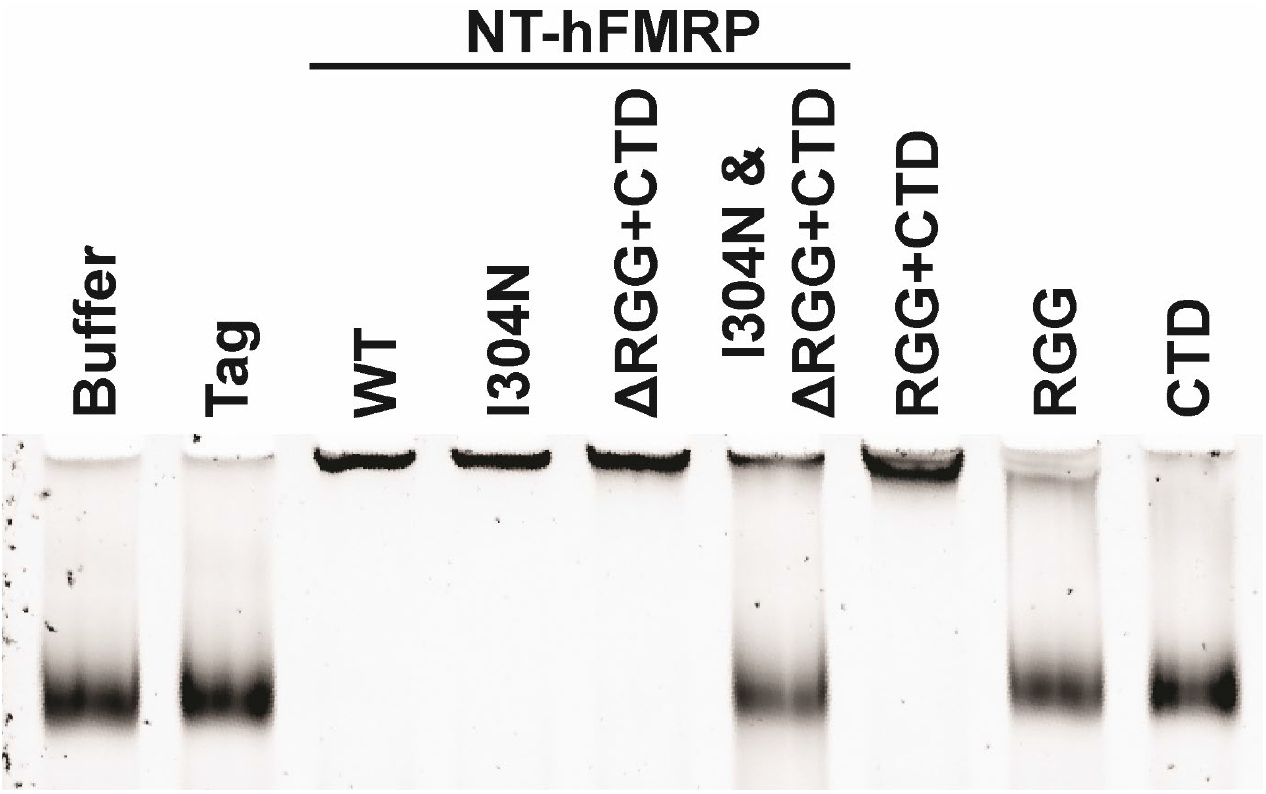
The RGG+CTD ncRBD binds G4-less mEGFP mRNA. EMSA of G4-less mEGFP mRNA with the indicated recombinant protein resolved on a native PAGE and subsequently stained with SYBR Green II.

**Supplemental Figure S7.**
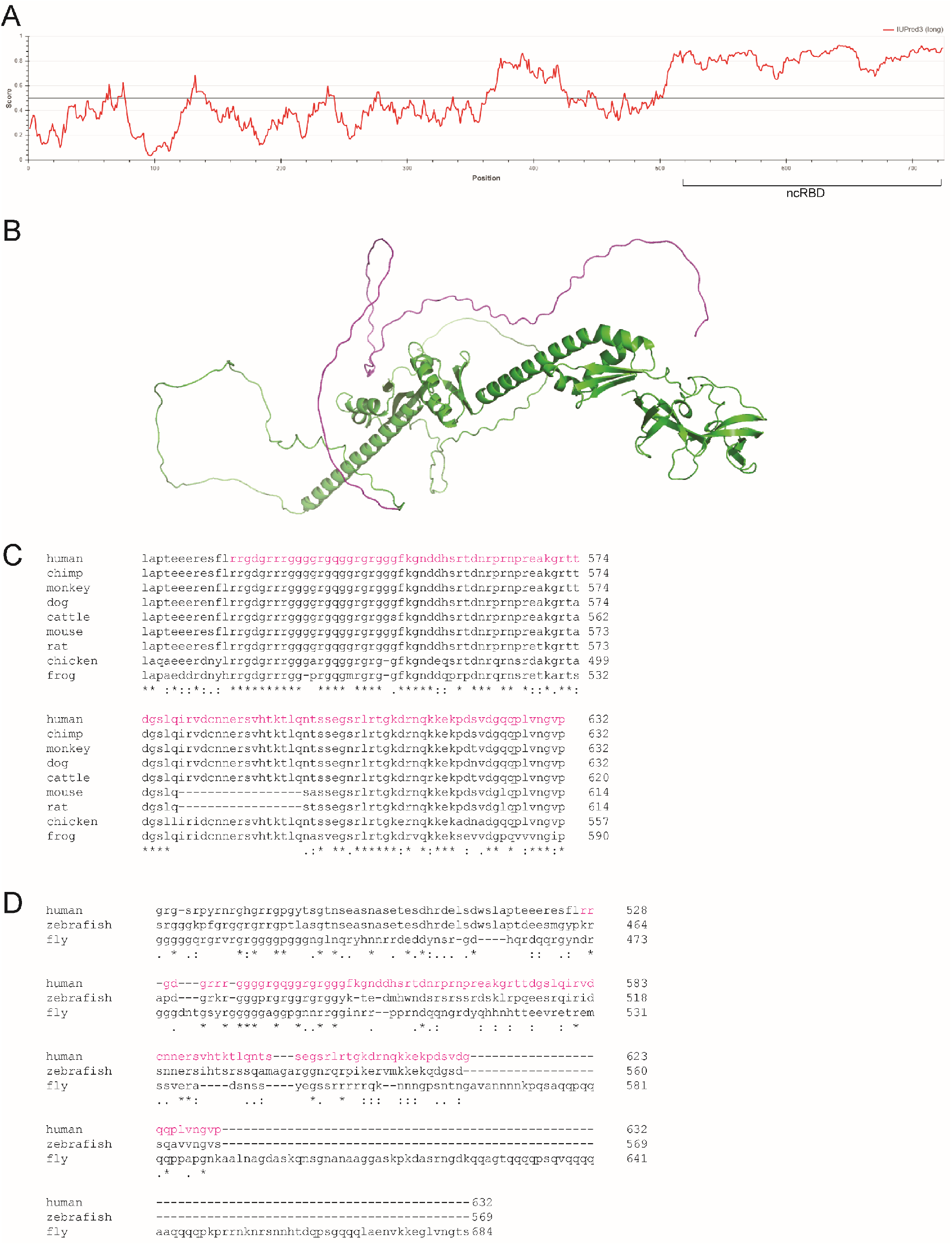
The FMRP ncRBD is predicted to be flexible and is conserved among most vertebrates. A) IUPred3 analysis (https://iupred.elte.hu/) of the full-length human FMRP. The ncRBD is bracketed. Scores above 0.5 (solid horizontal line) signal predicted unstructured flexible regions. B) AlphaFold structural prediction of full-length human FMRP with the ncRBD highlighted in magenta. C) Clustal Omega (1.2.4) alignment of vertebrate FMRP orthologs. The human FMRP ncRBD is highlighted in magenta. The residue numbering corresponds to the full-length protein sequence. D) Clustal Omega (1.2.4) alignment of human, zebrafish, and fruit fly FMRP. The human FMRP ncRBD is highlighted in magenta. The residue numbering corresponds to the full-length protein sequence.

**Supplemental Figure S8.**
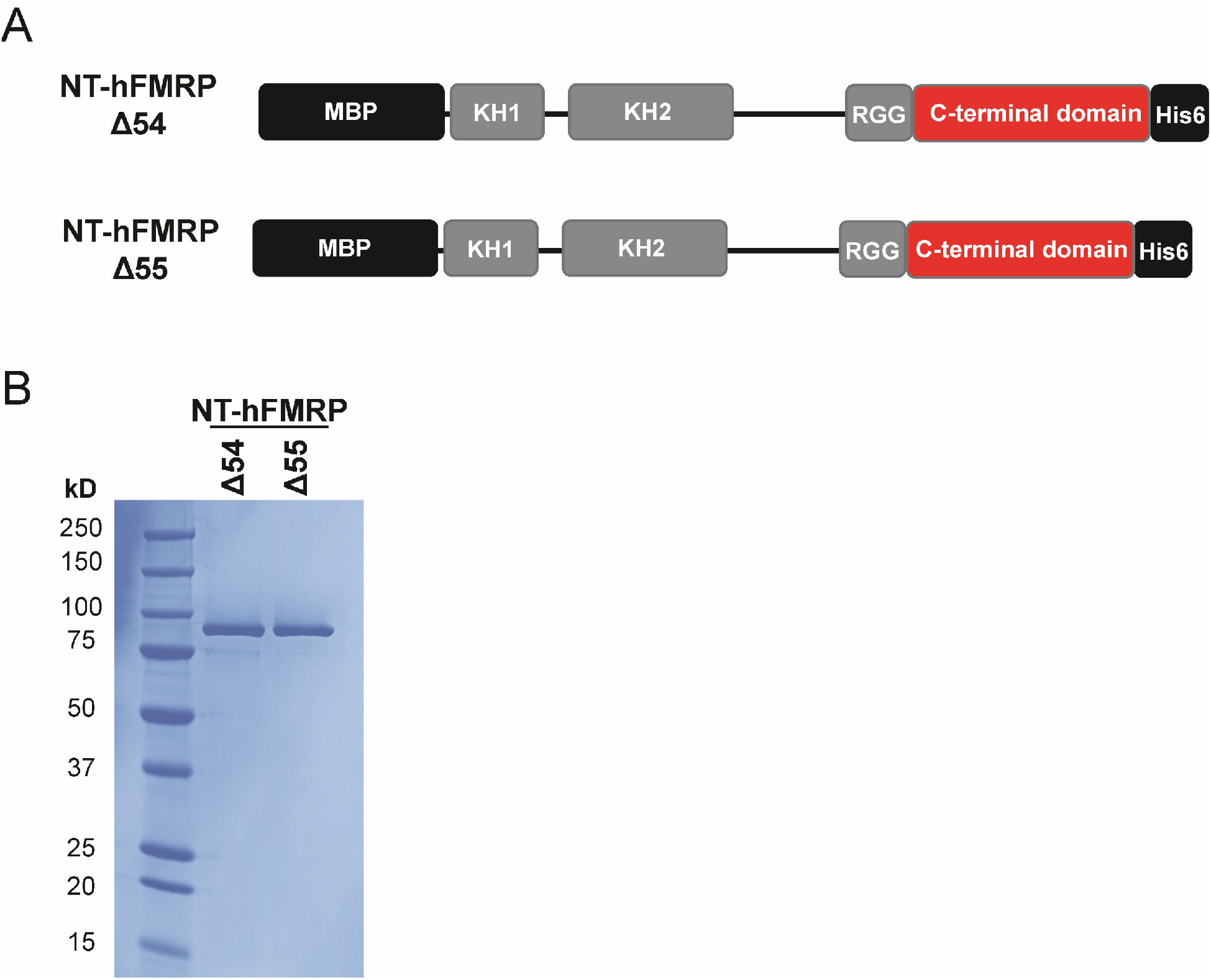
NT-hFMRP Δ54 and Δ55. A) Schematic of MBP- and His6-tagged NT-hFMRP Δ54 and Δ55. The truncated CTD is highlighted in red. B) Coomassie stain of recombinant NT-hFMRP Δ54 and Δ55.

**Supplemental Figure S9.**
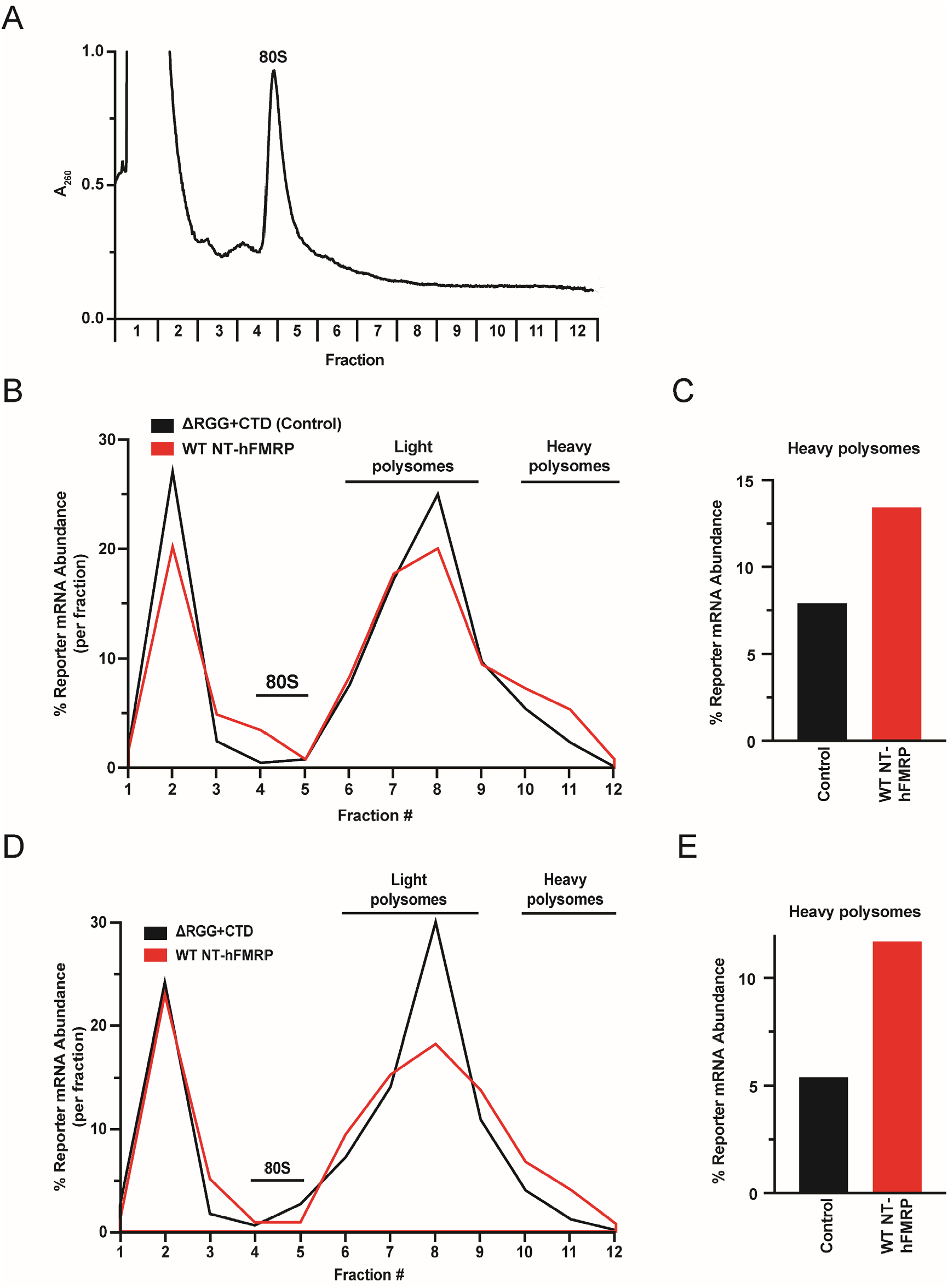
WT NT-hFMRP causes heavy polysomes to accumulate. A) A representative A_260 nm_ trace of sucrose gradient ultracentrifugation (polysome analysis) of *in vitro* translation reactions. The 80S monosome sediments in fractions 4 and 5. Thus, fractions 6-12 contain polysomes. Robust polysome curves are not visualized as mRNA input amounts were limited to ensure responses were in the linear dynamic range (see Materials and Methods). B-E) Separate biological replicates of experiments shown in **Figure 6B-D**. B & D) Distribution of reporter mRNA across sucrose gradients to assess polysome formation with ΔRGG+CTD (Control) or WT NT-hFMRP. Abundance of reporter mRNA in each gradient fraction was determined by RT-qPCR. C & E) Cumulative nLuc abundance in heavy polysomes in fractions 10-12 from ΔRGG+CTD (Control) and WT NT-hFMRP samples.

**Supplemental Figure S10.**
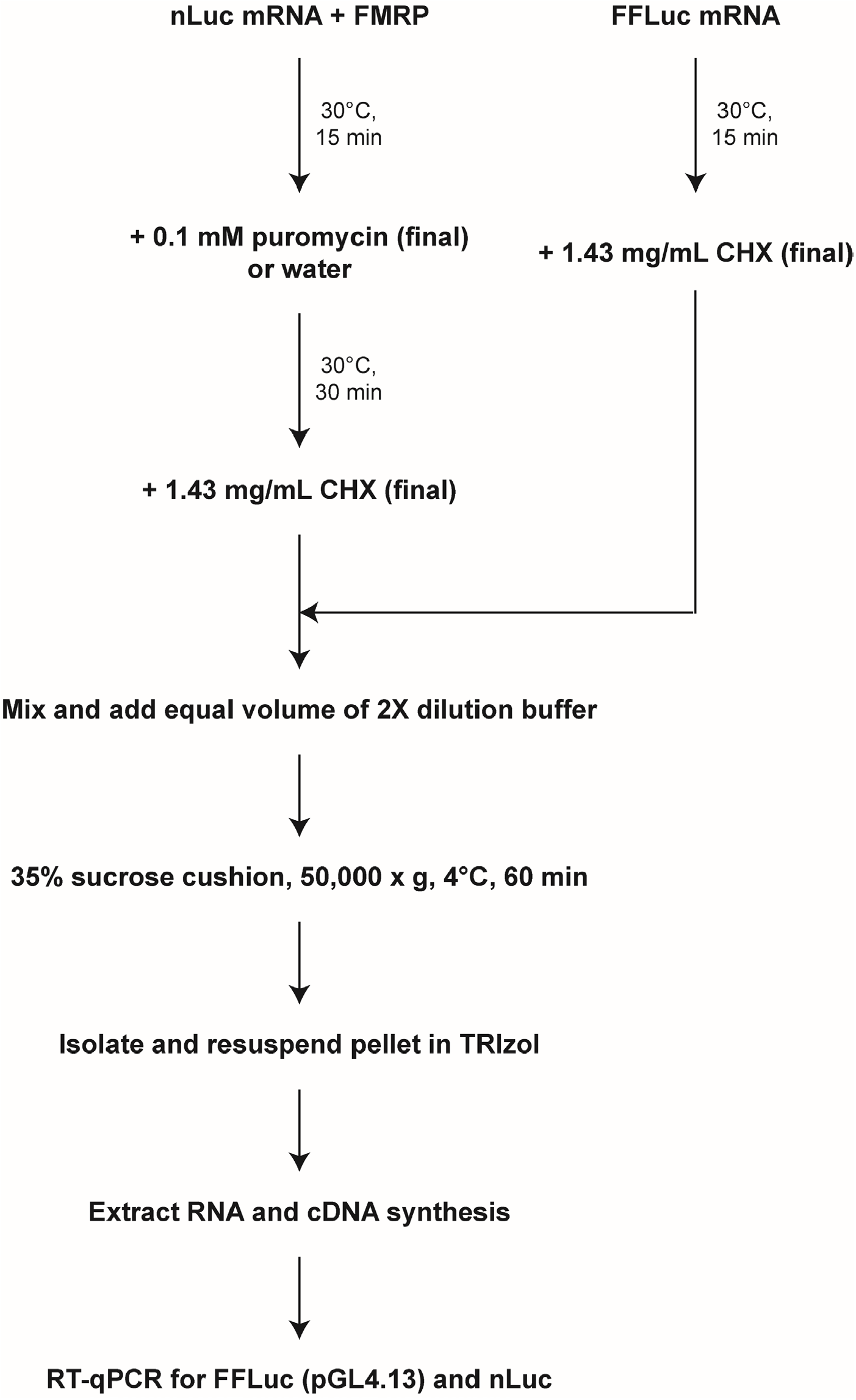
Workflow of puromycin-mediated ribosome dissociation and low-speed sucrose cushion assay. WT or mutant NT-hFMRP was allowed to form an mRNP with nLuc mRNA as described in the Material and Methods. FFLuc (pGL4.13) mRNA was translated and used as an internal spike-in control for ribosome-bound mRNA pelleting and RT-qPCR. See details in the Material and Methods.

### Reporter Sequences

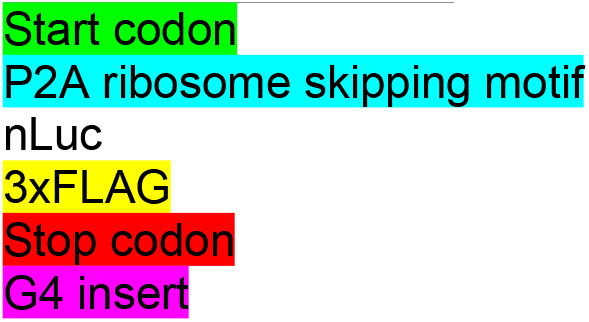

### Control nLuc reporter

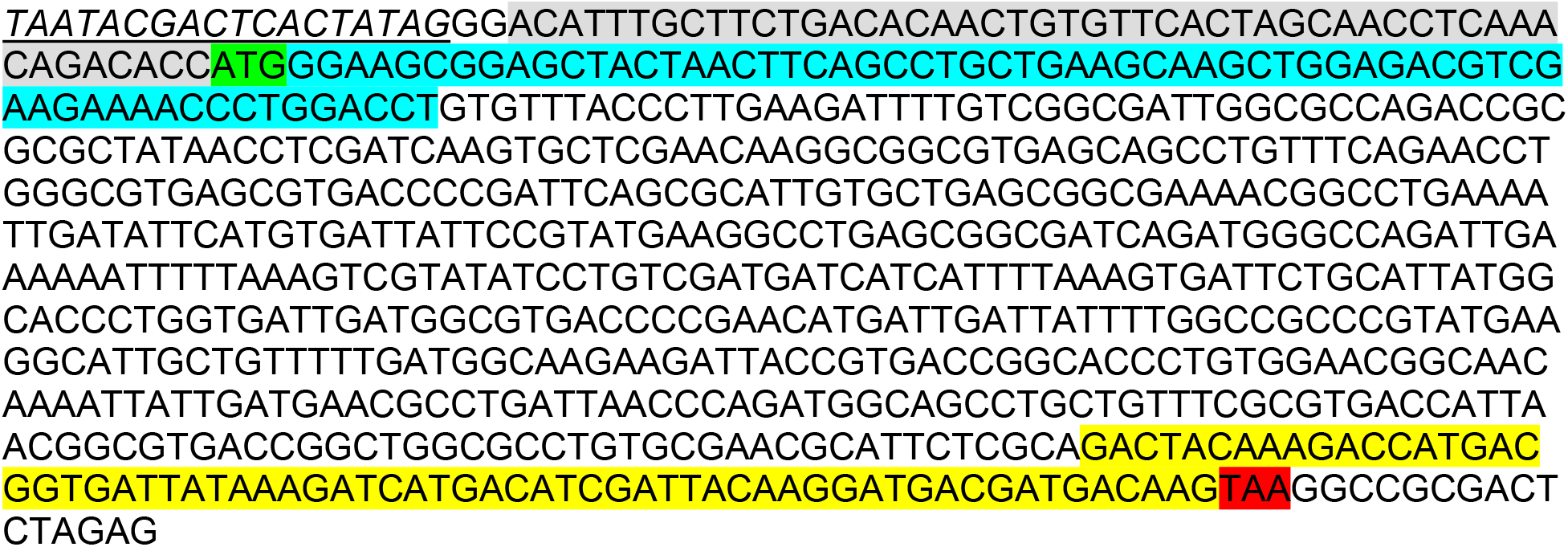

### G_15_ nLuc reporter

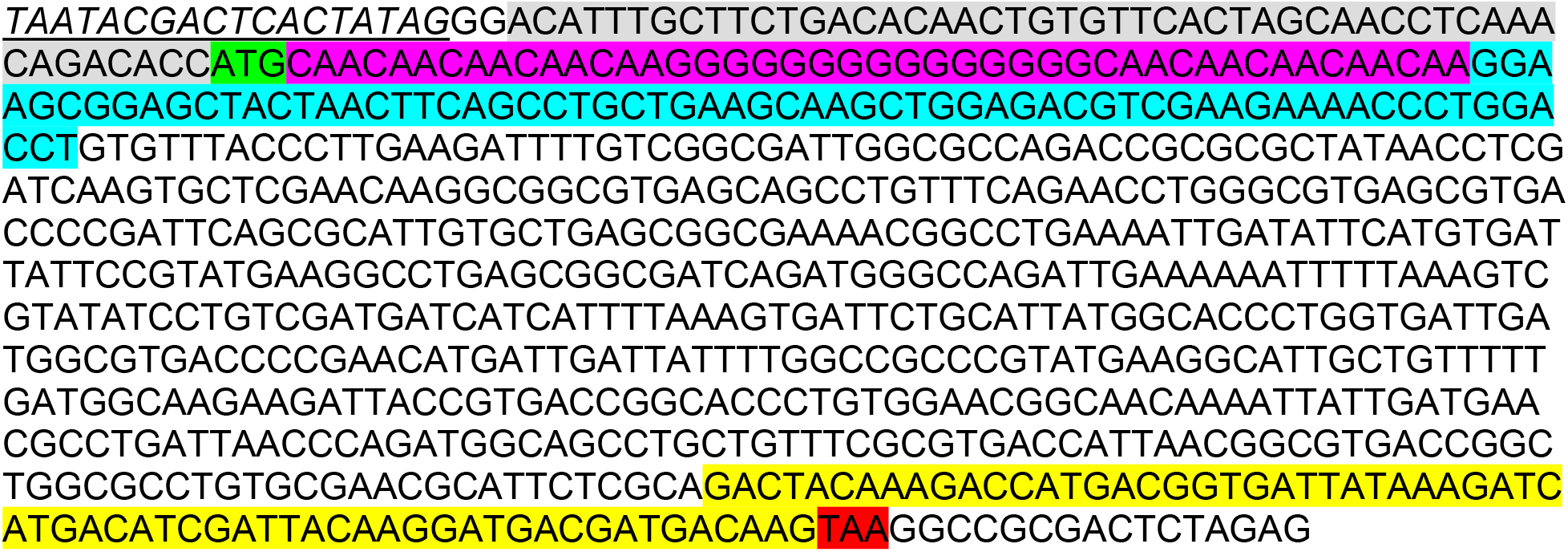

### (GGGU)_4_ nLuc reporter

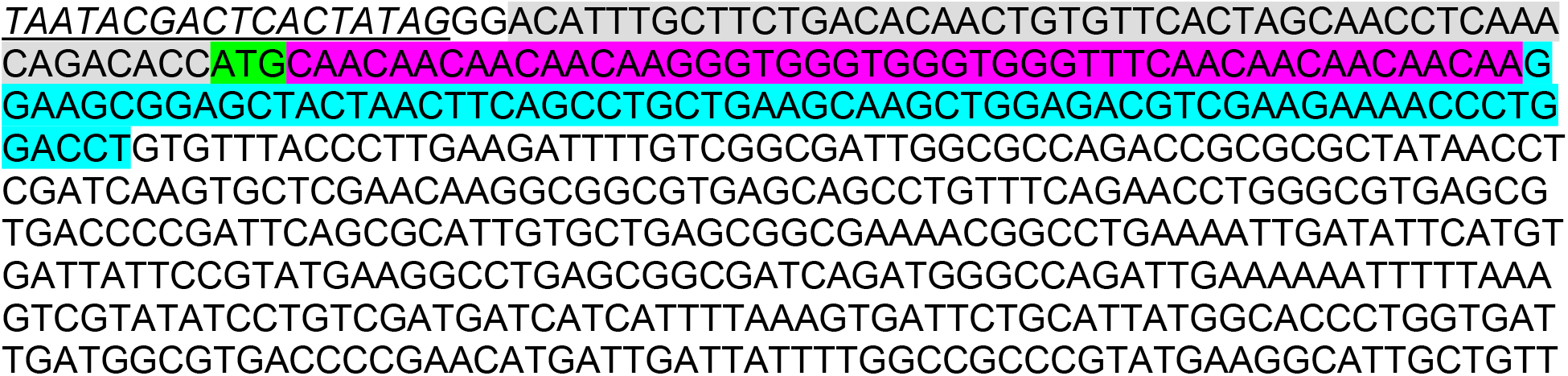
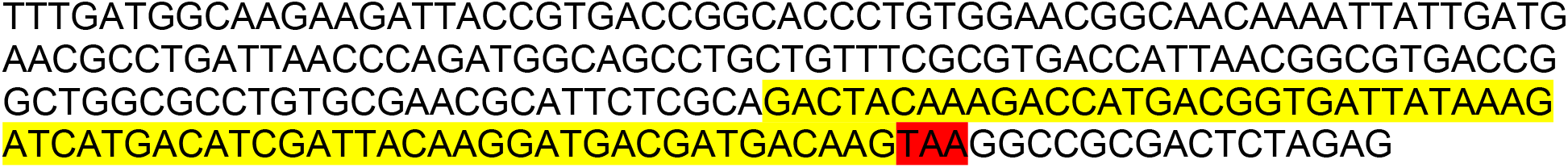

### Sc1 nLuc reporter

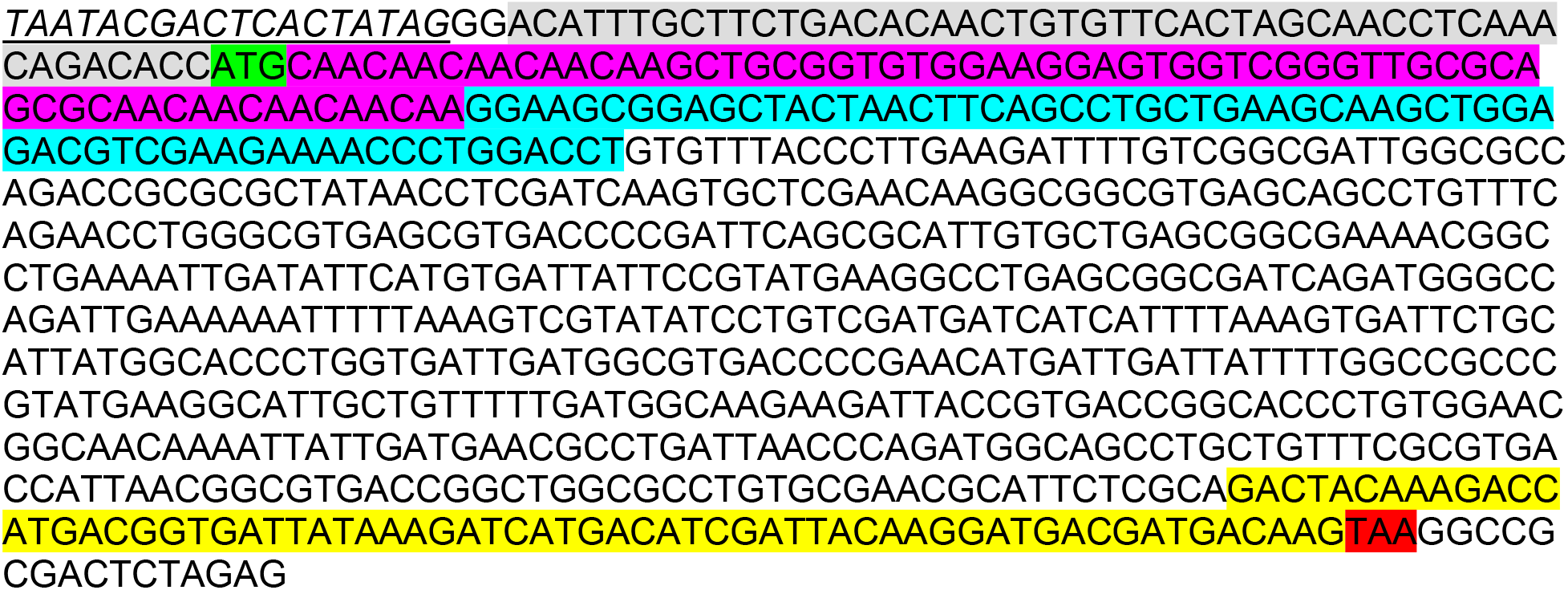

### Long 5′ UTR nLuc reporter

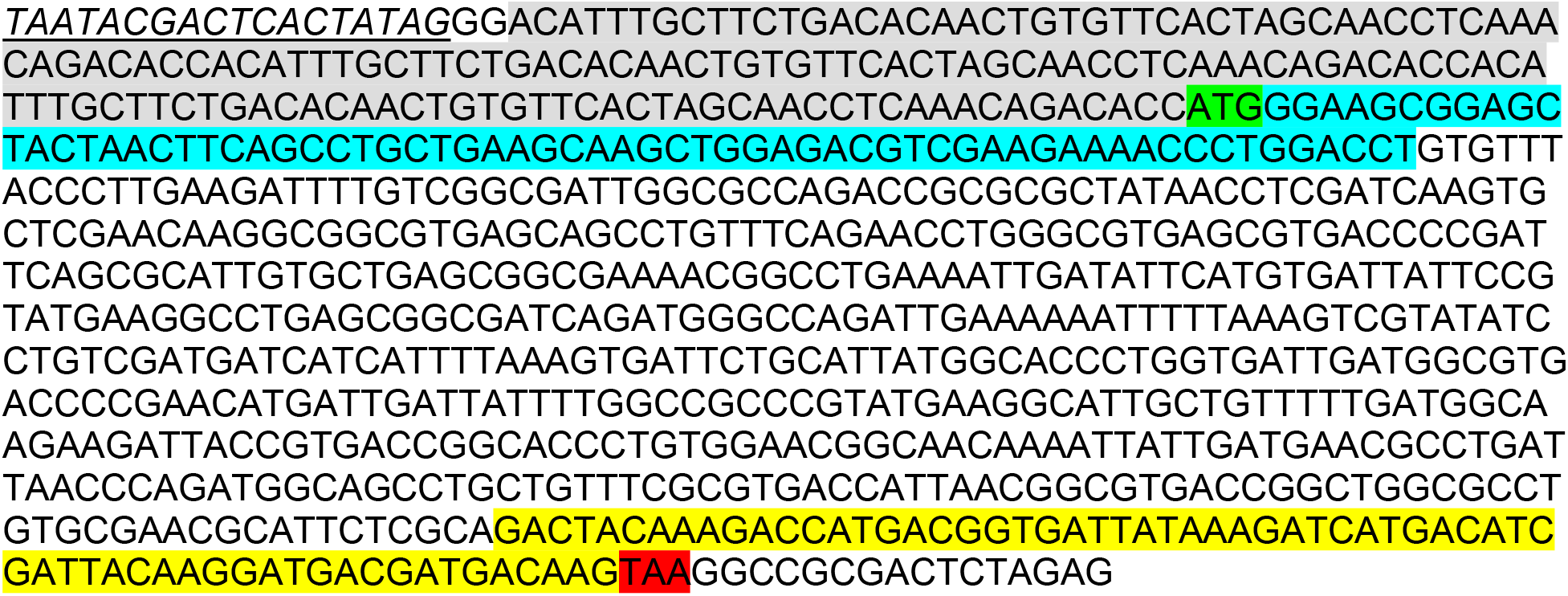

### G4-less mEGFP reporter

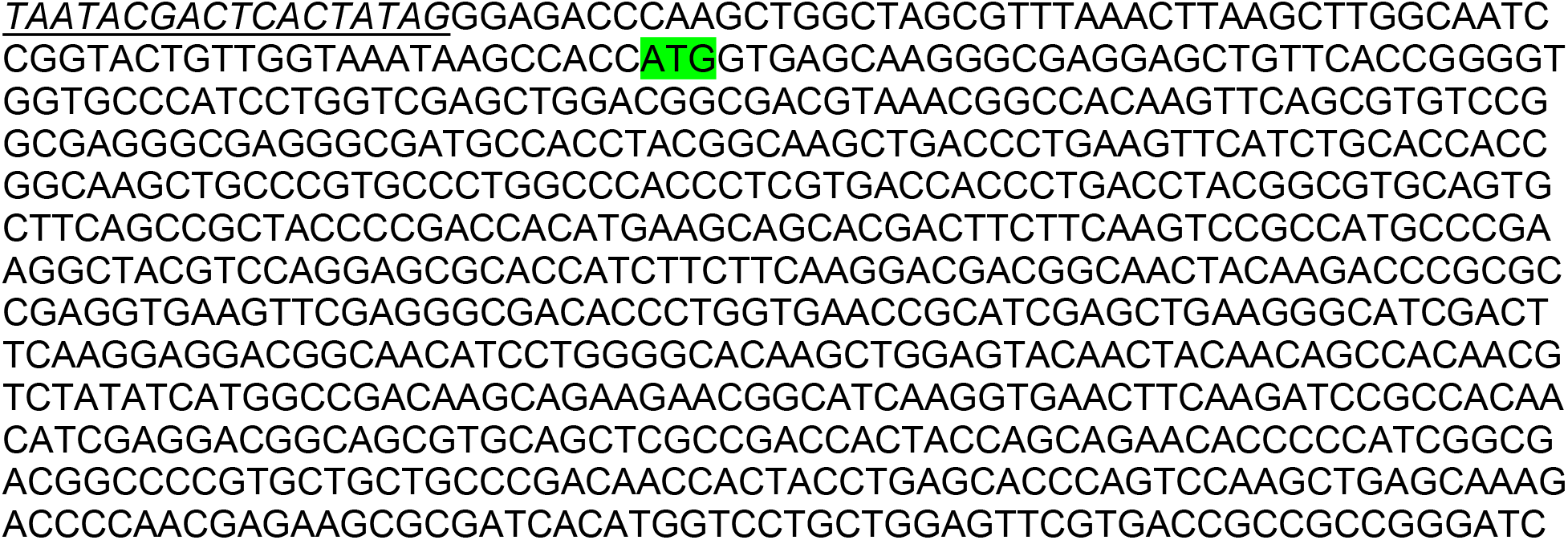
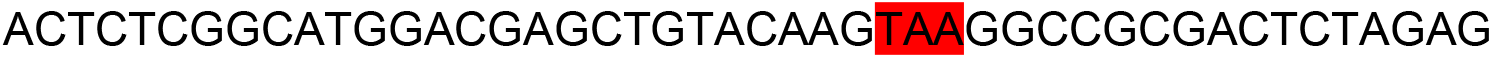

### Recombinant Protein Coding Sequences

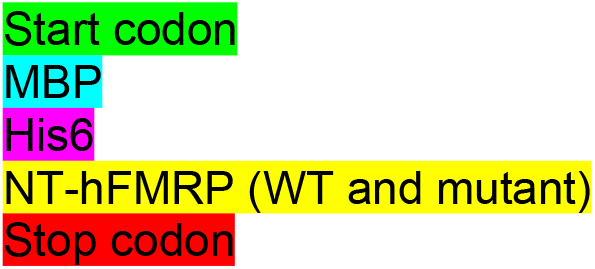

### His6-MBP

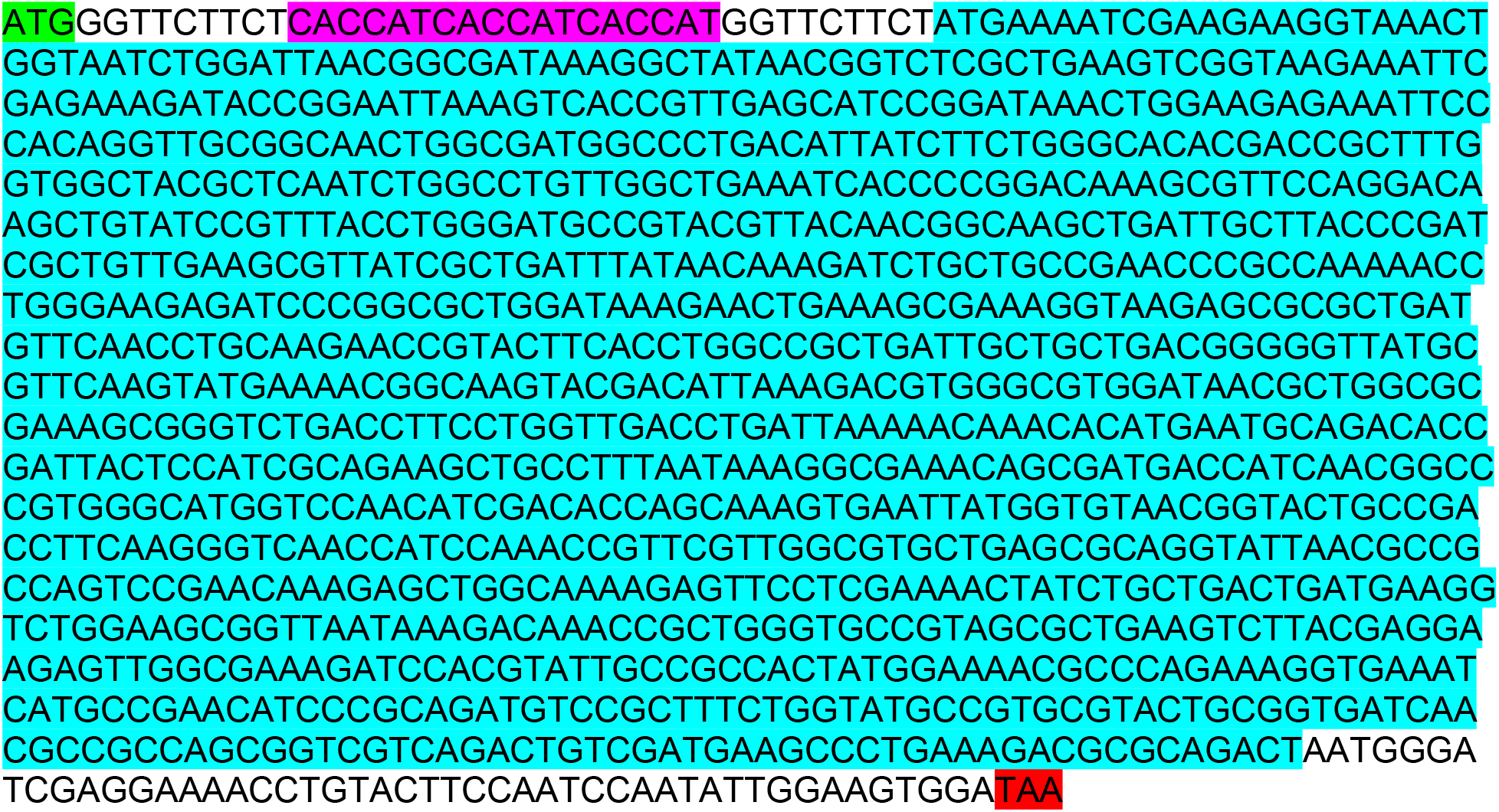

### MBP-(WT NT-hFMRP)-His6

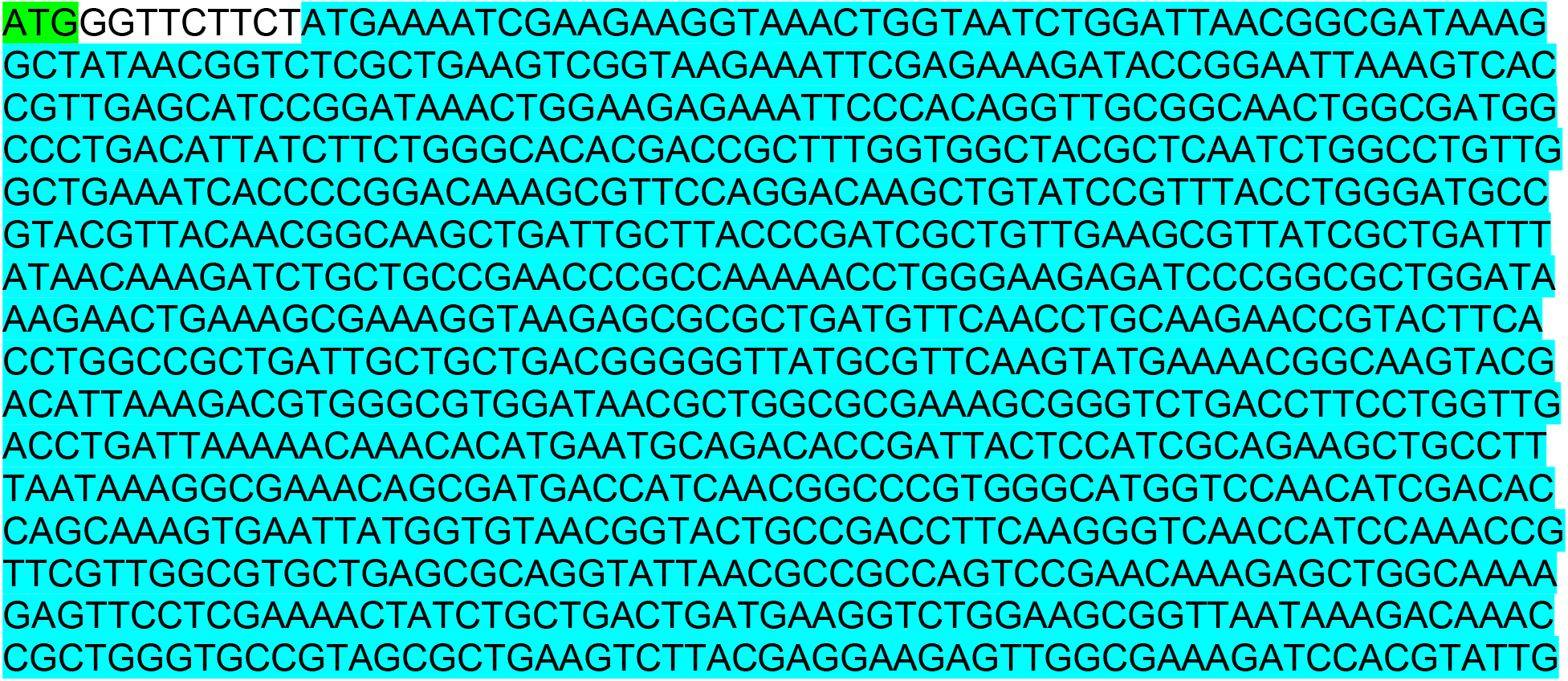
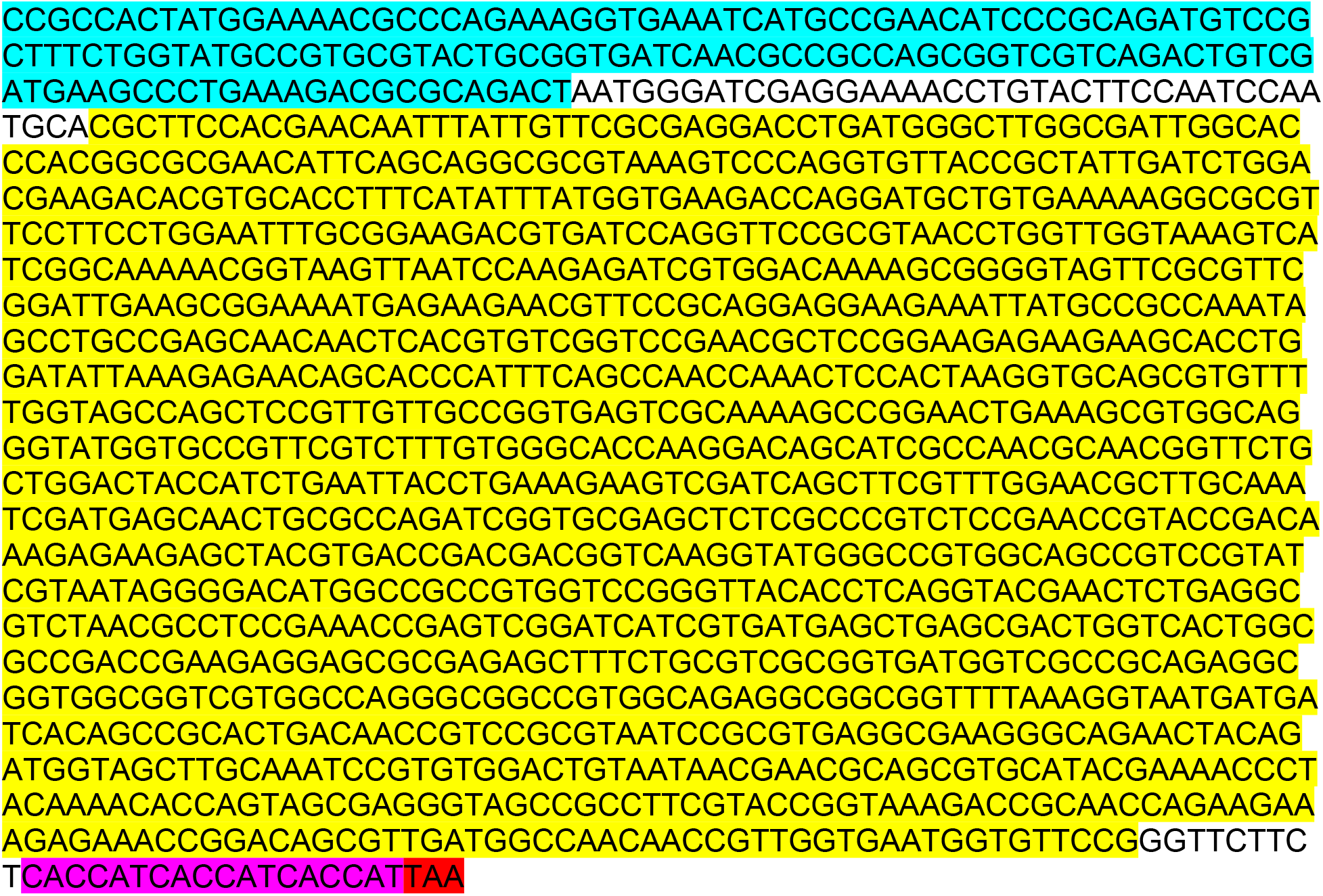

### MBP-(I304N NT-hFMRP)-His6

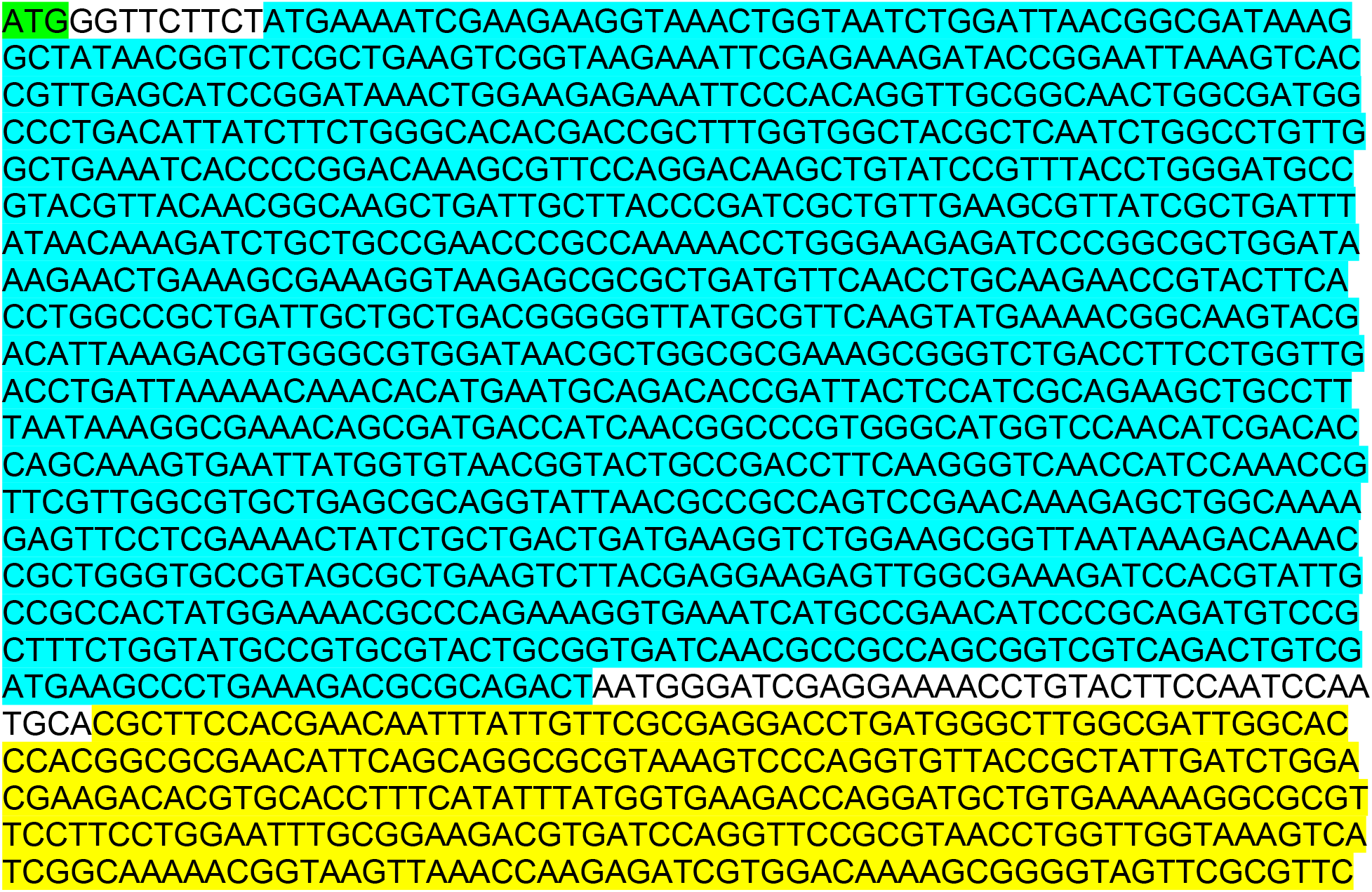
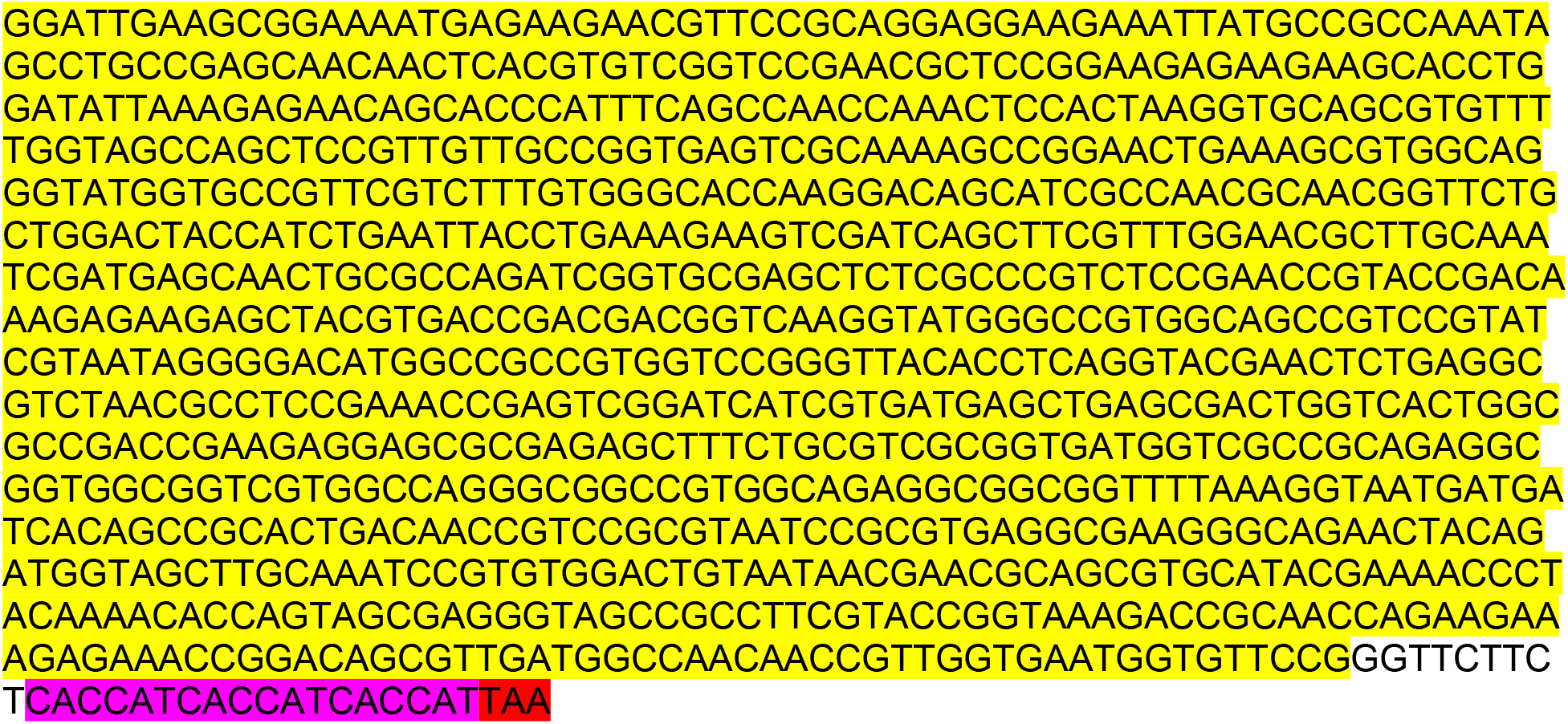

### MBP-(ΔRGG+CTD NT-hFMRP)-His6

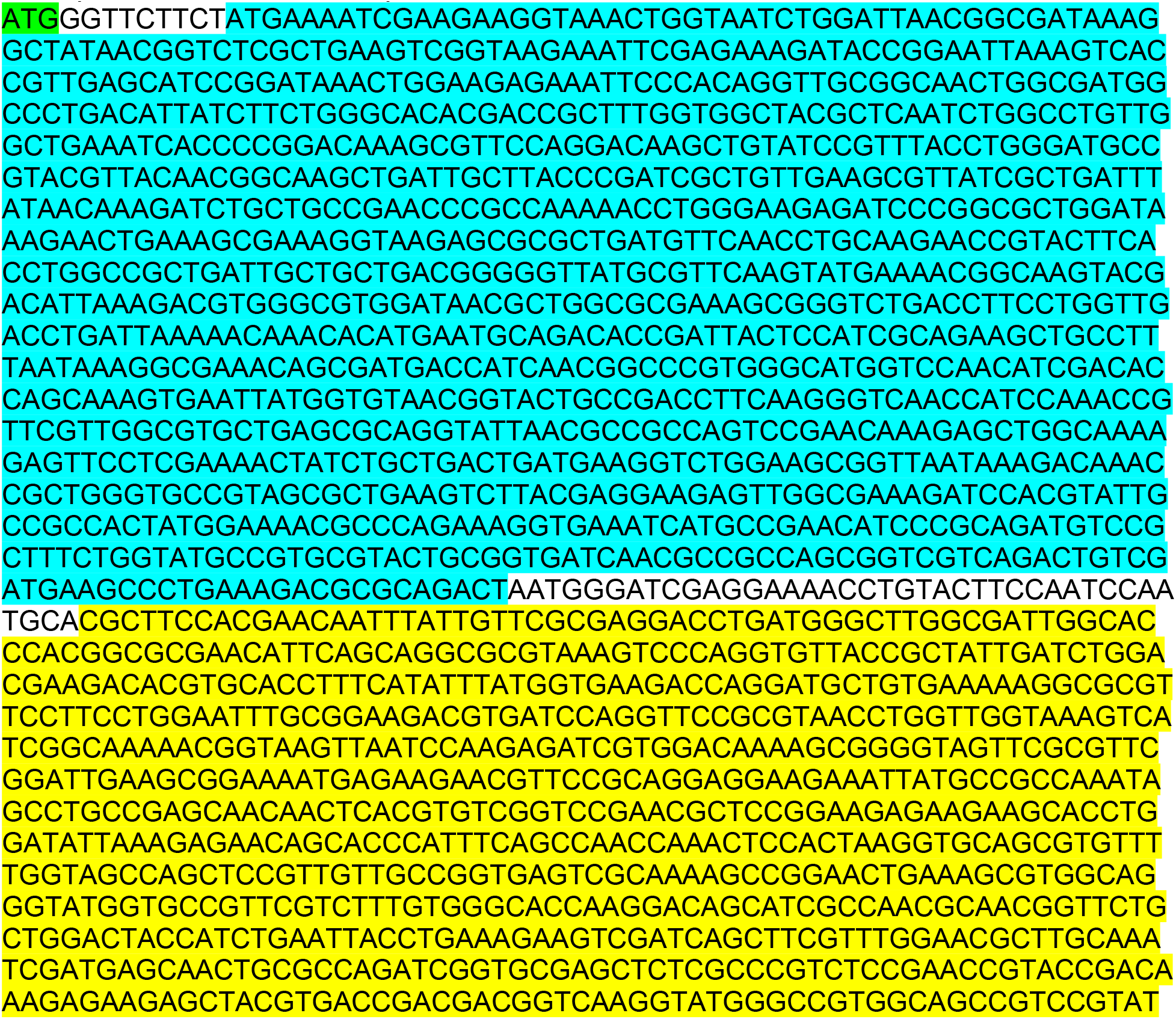
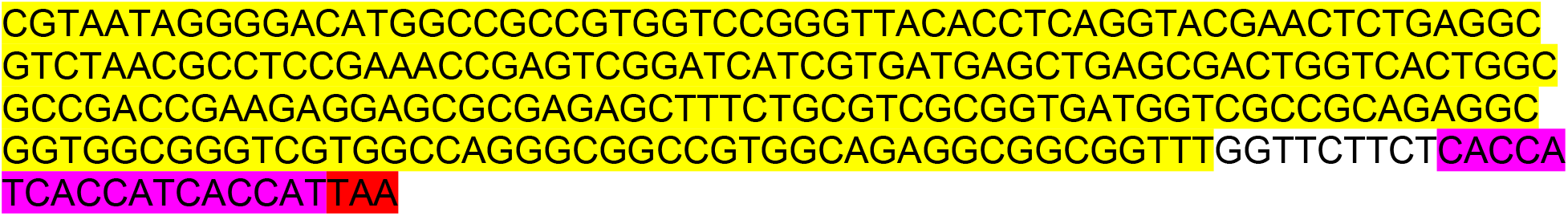

### MBP-(NT-hFMRP ΔRGG+CTD complete)-His6

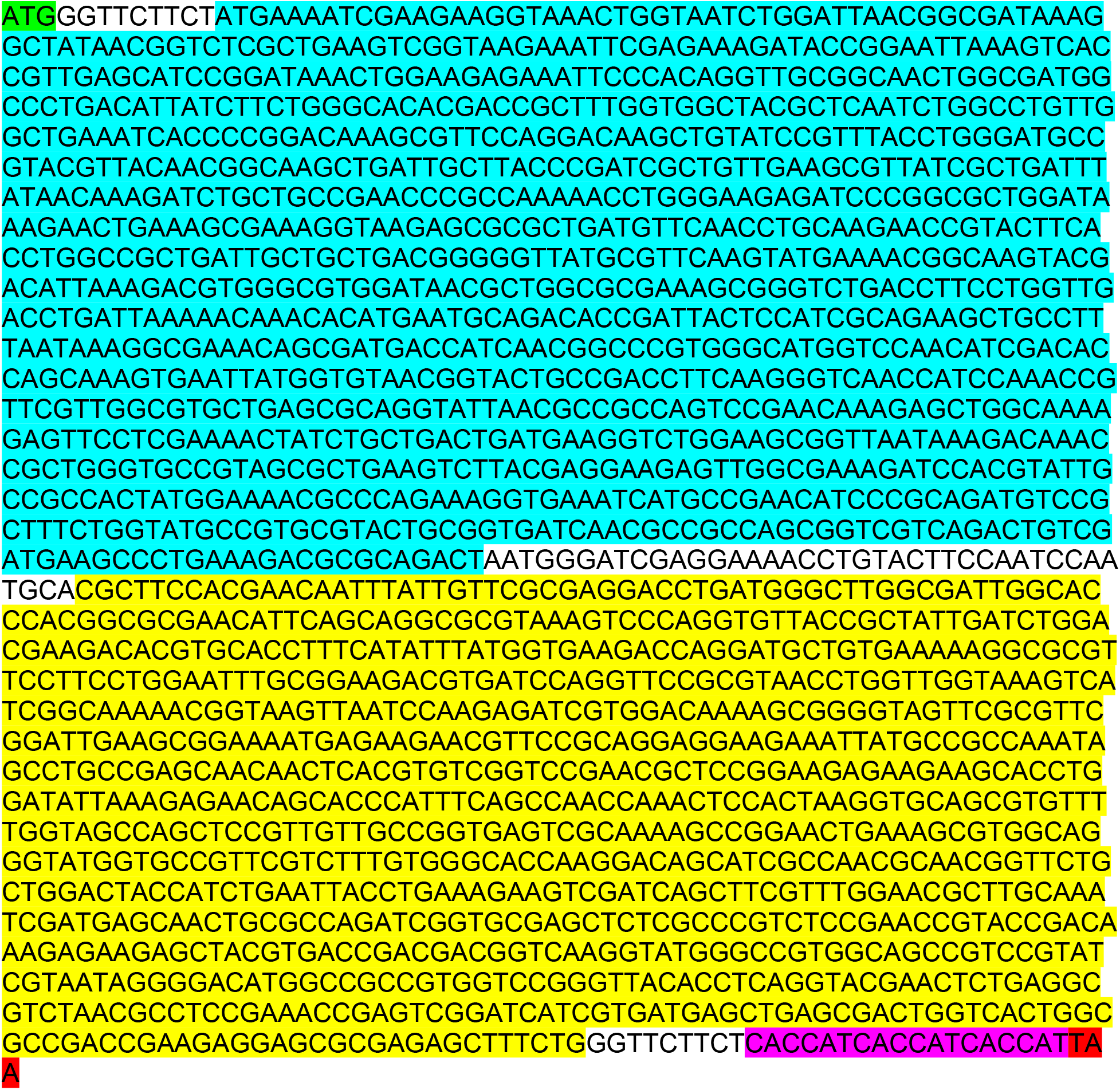

### MBP-(I304N & ΔRGG+CTD NT-hFMRP)-His6

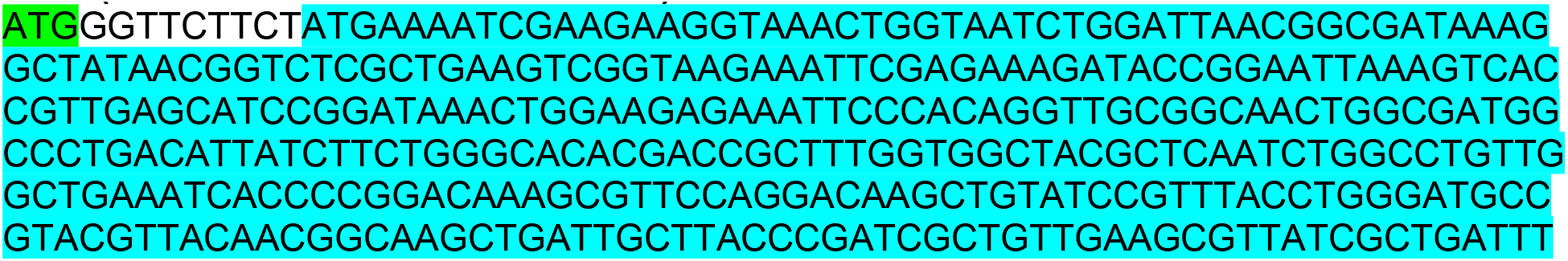
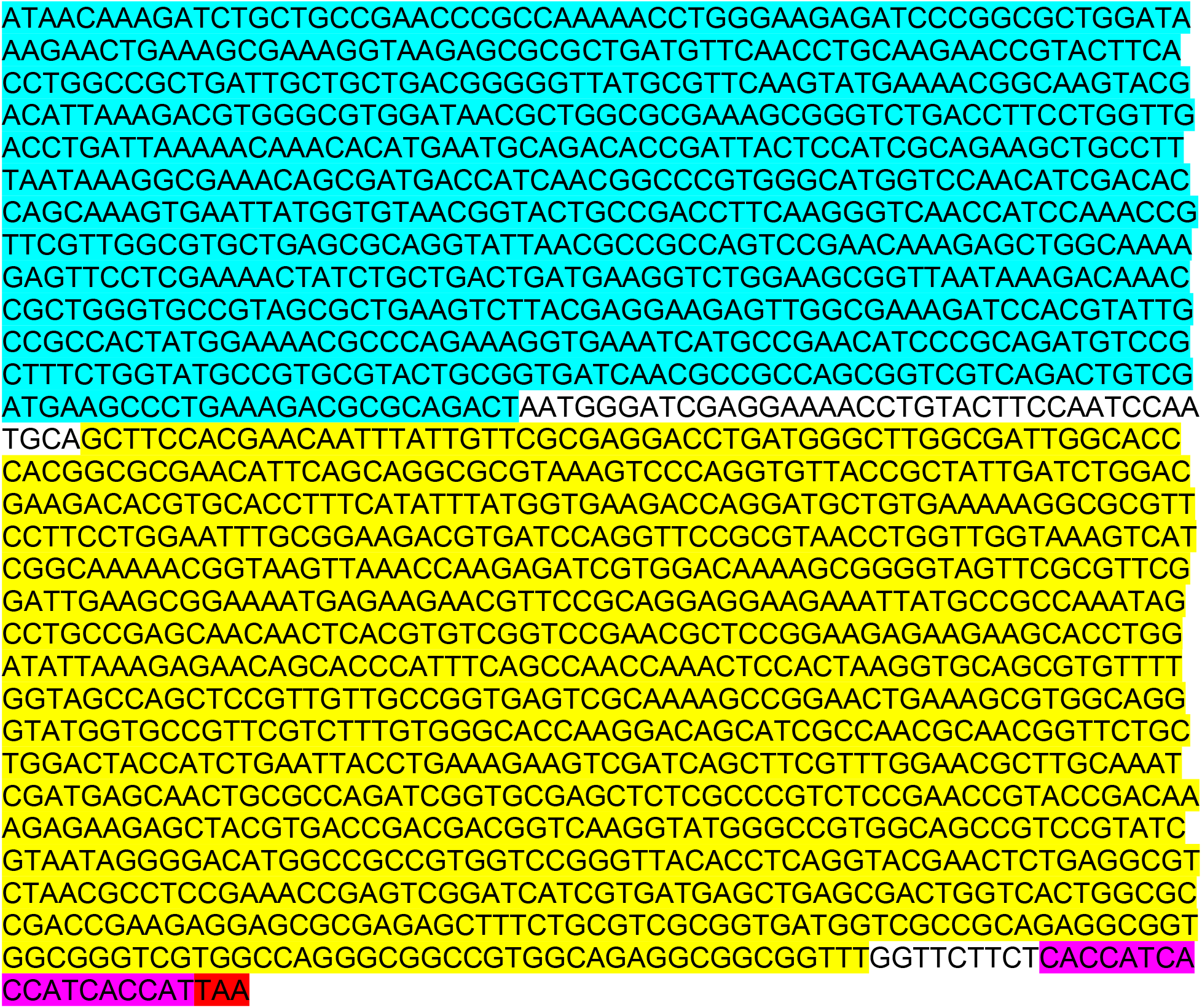

### MBP-(NT-hFMRP Δ54)-His6

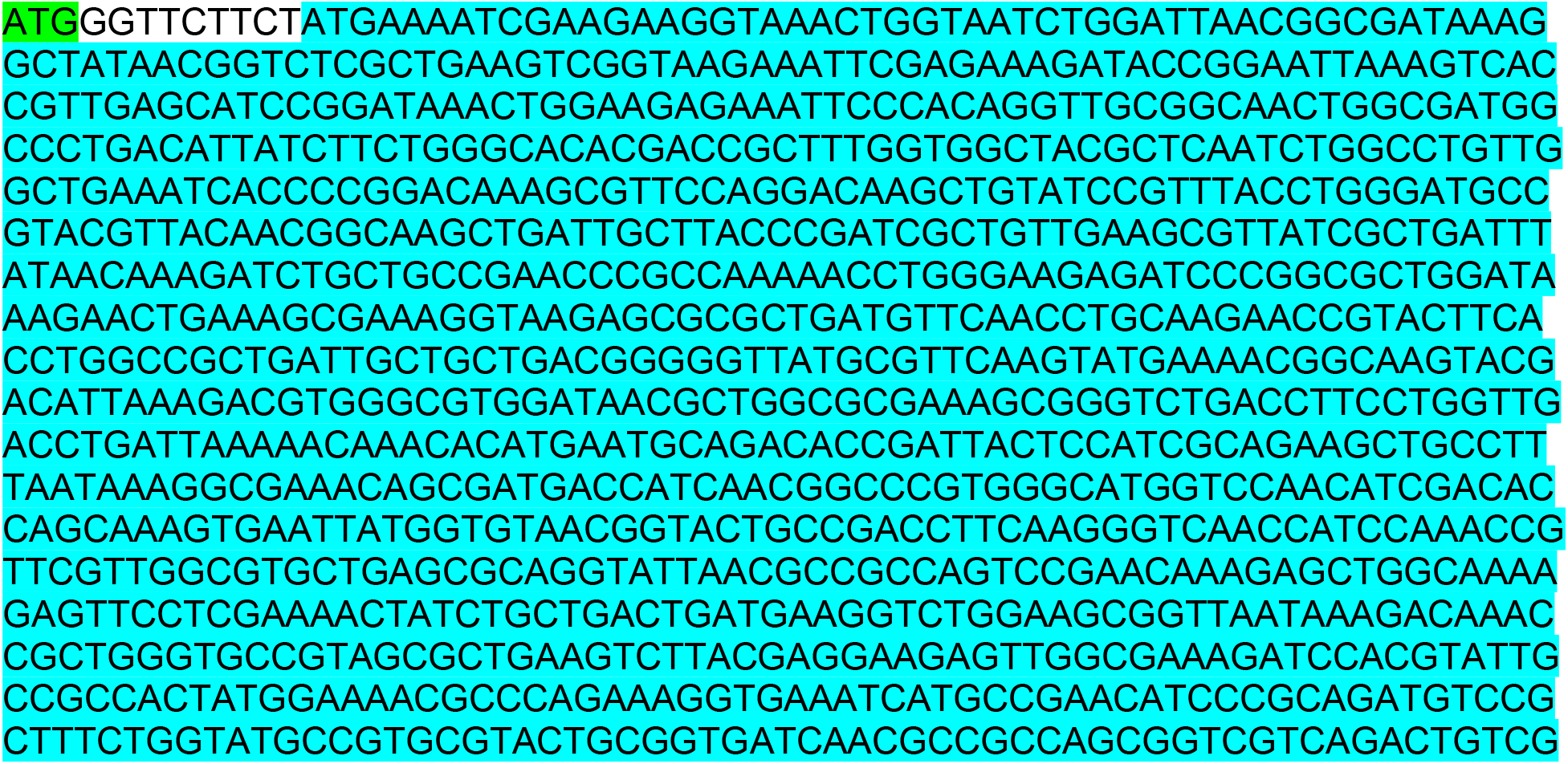
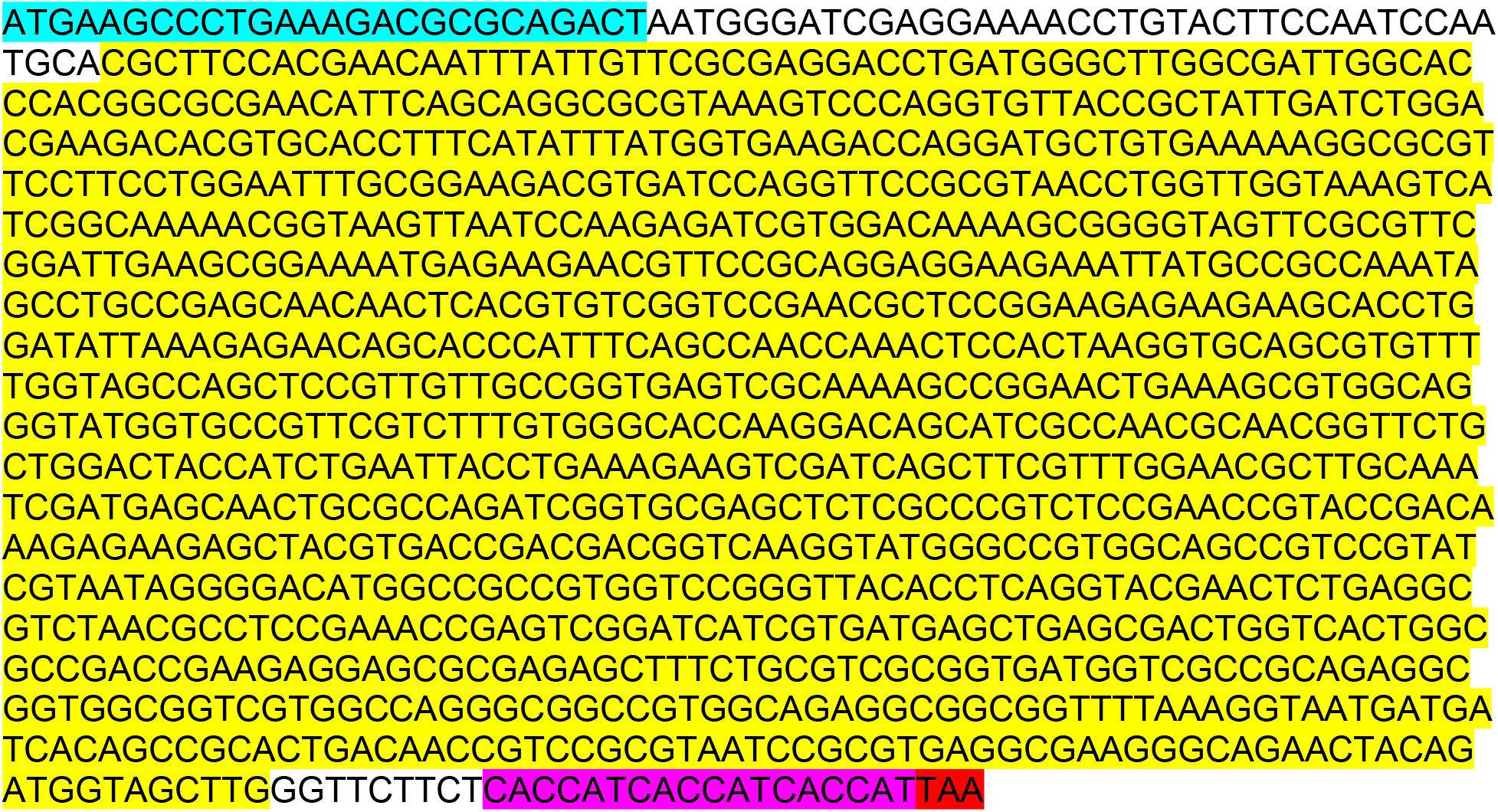

### MBP-(NT-hFMRP Δ55)-His6

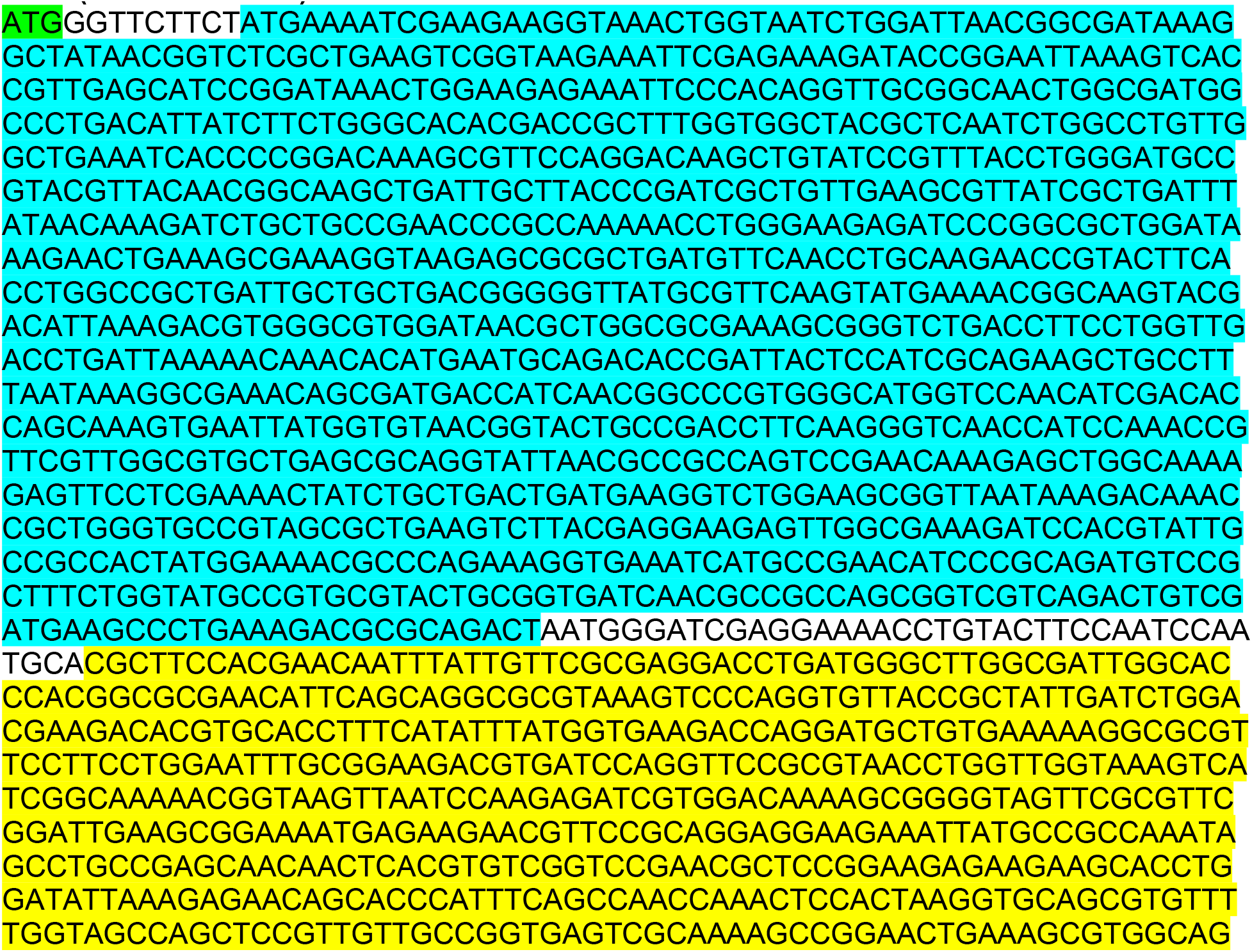
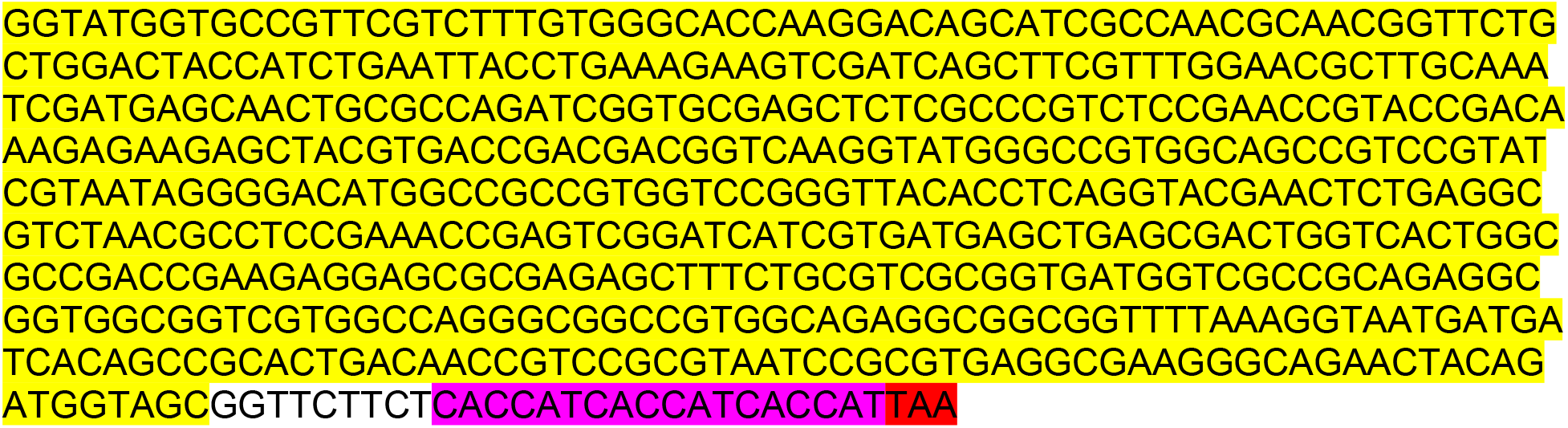

### MBP-(RGG+CTD)-His6

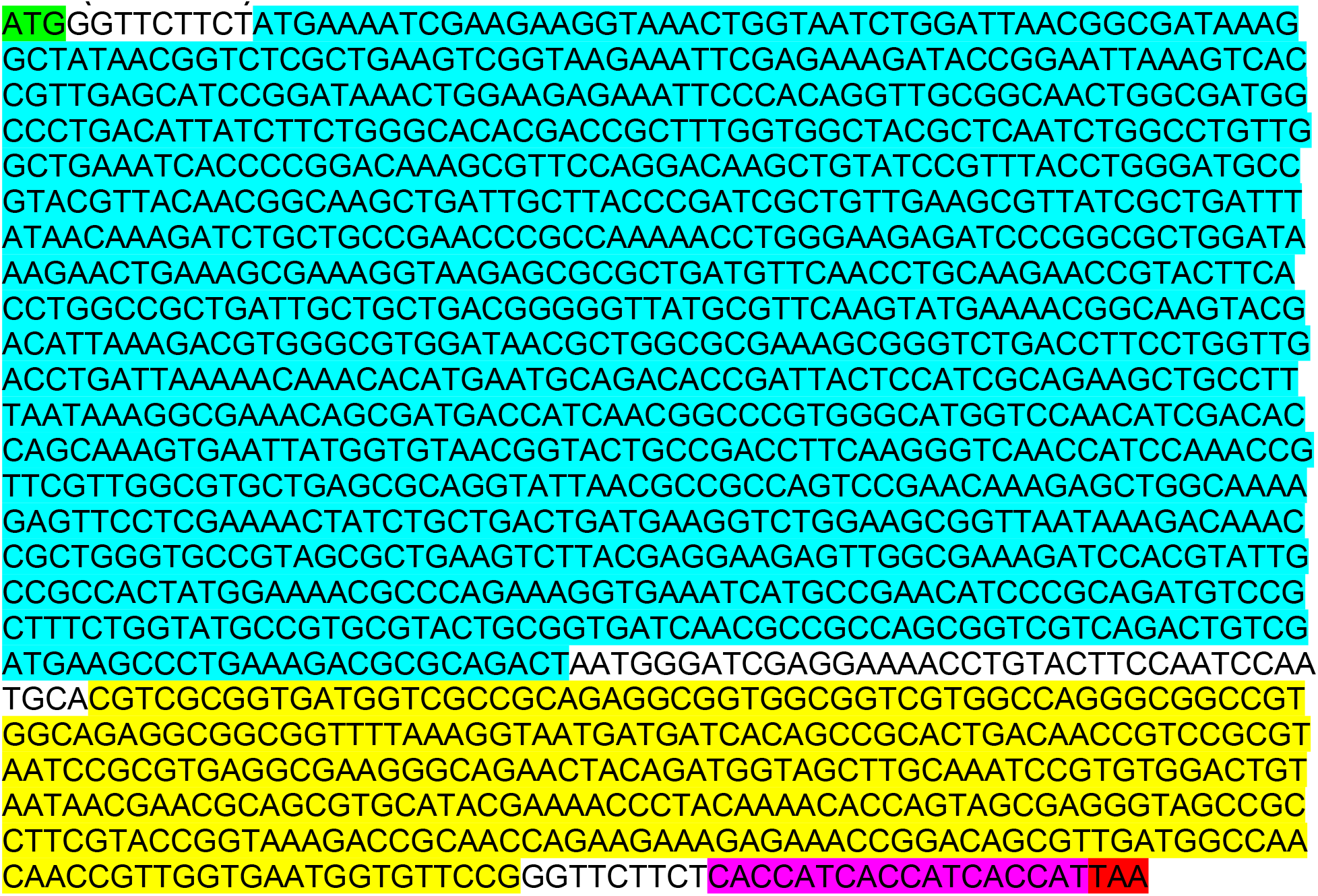

### MBP-(RGG)-His6

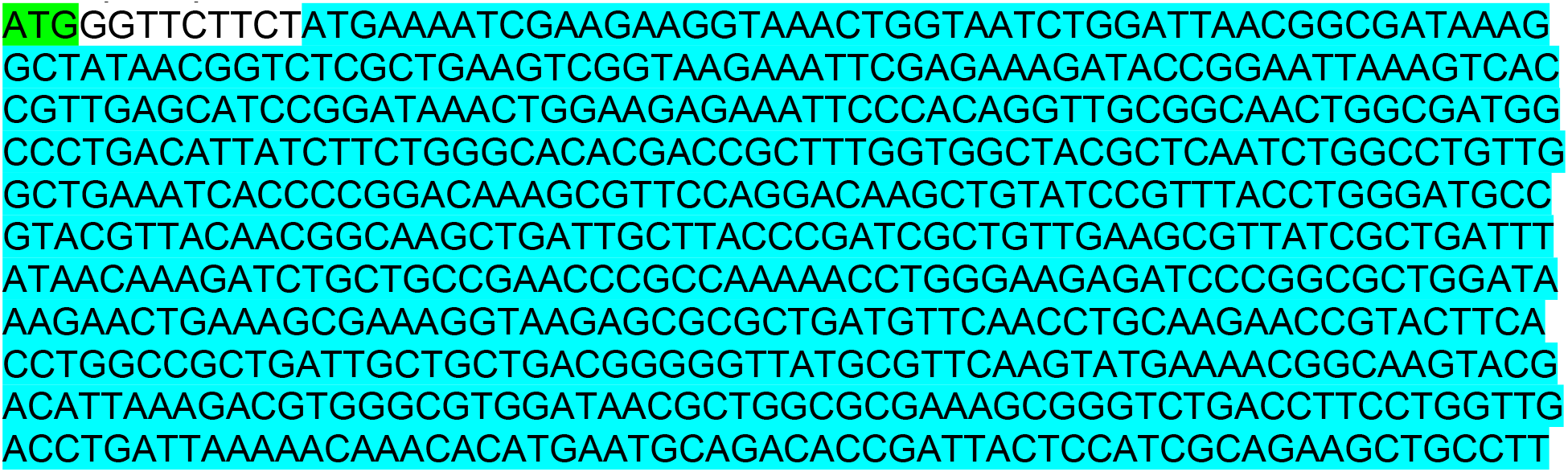
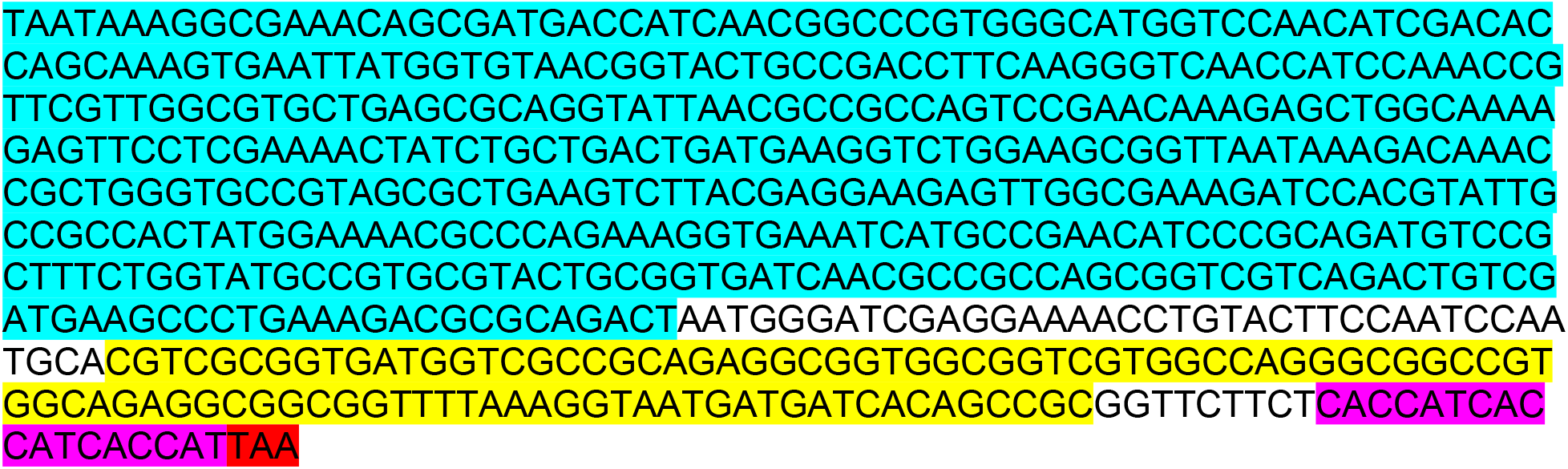

### MBP-(CTD)-His6

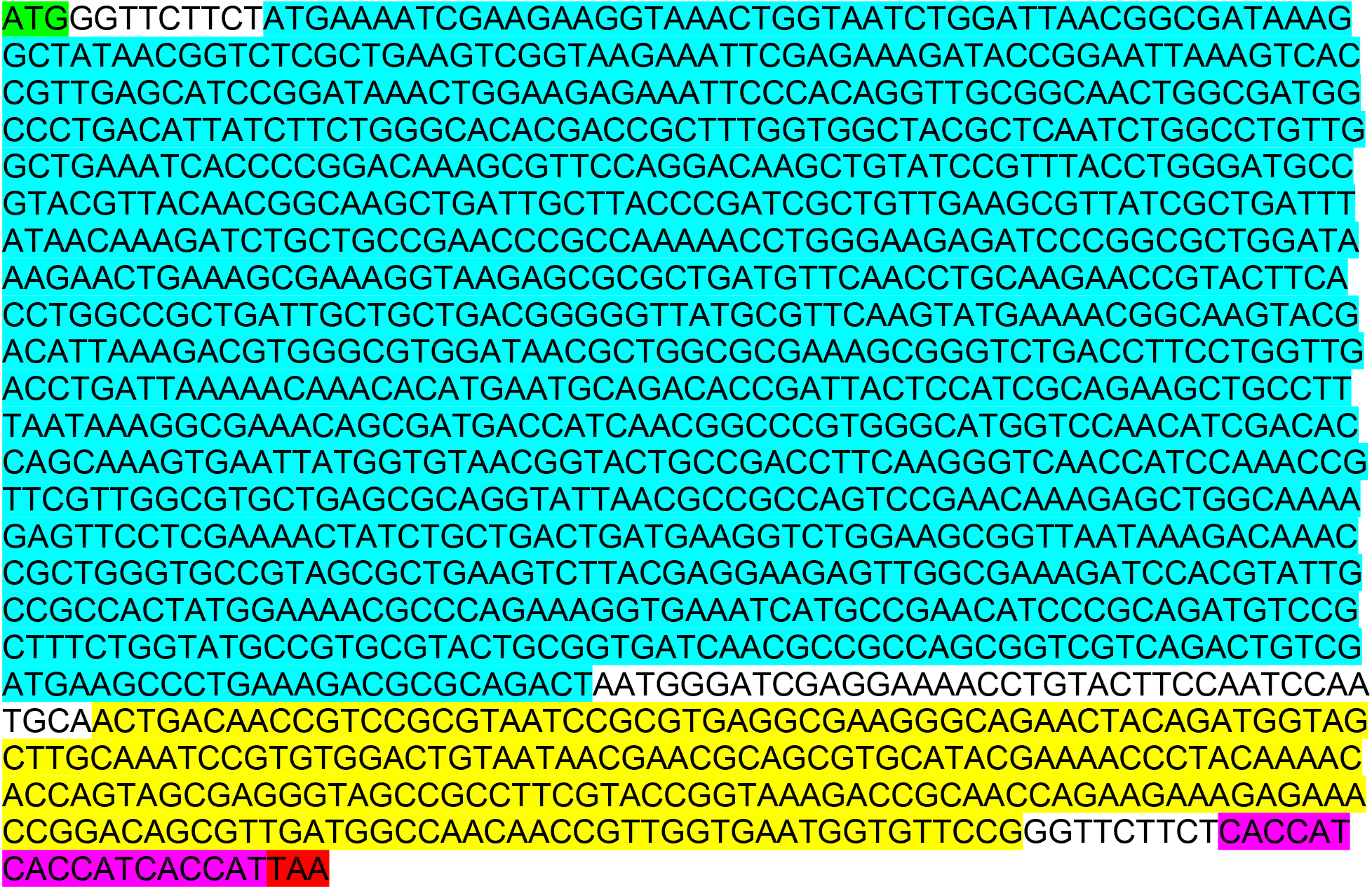

### MBP-(RGG+CTD Δ19)-His6

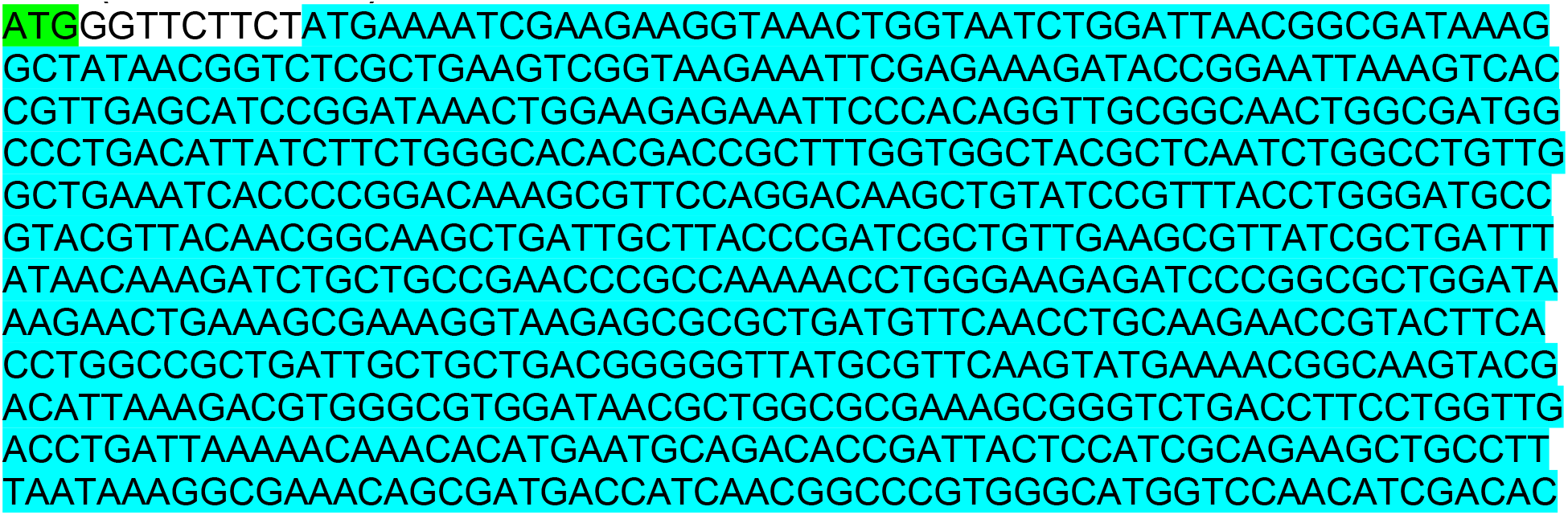
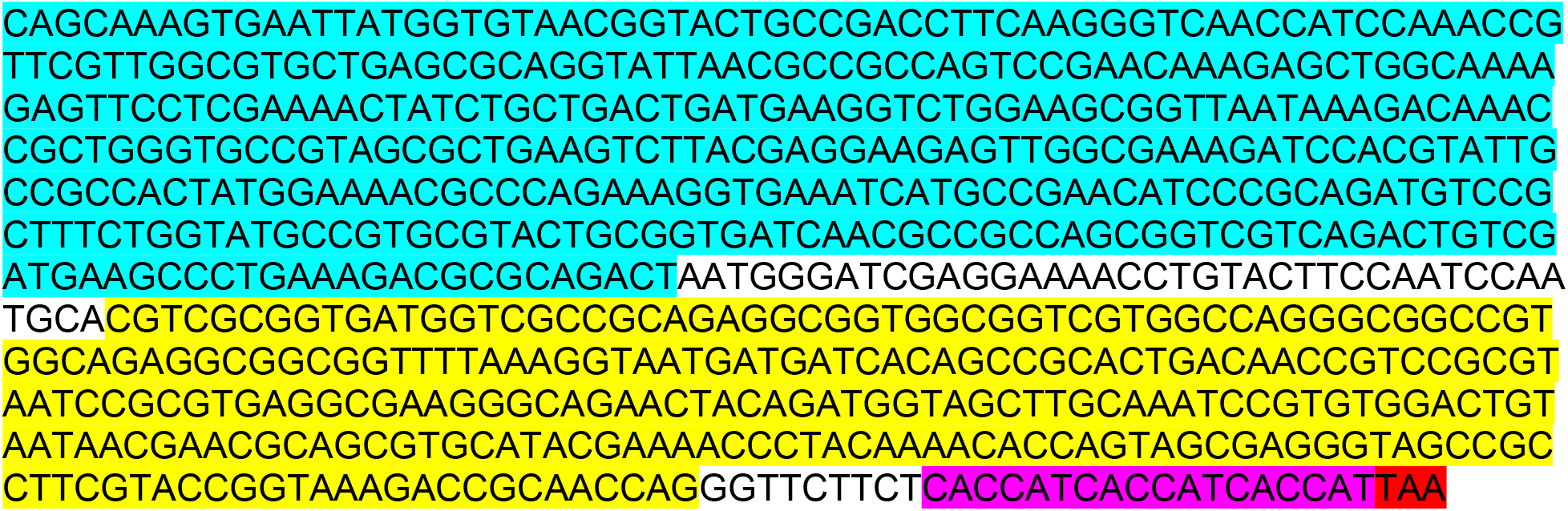

### MBP-(RGG+CTD Δ54)-His6

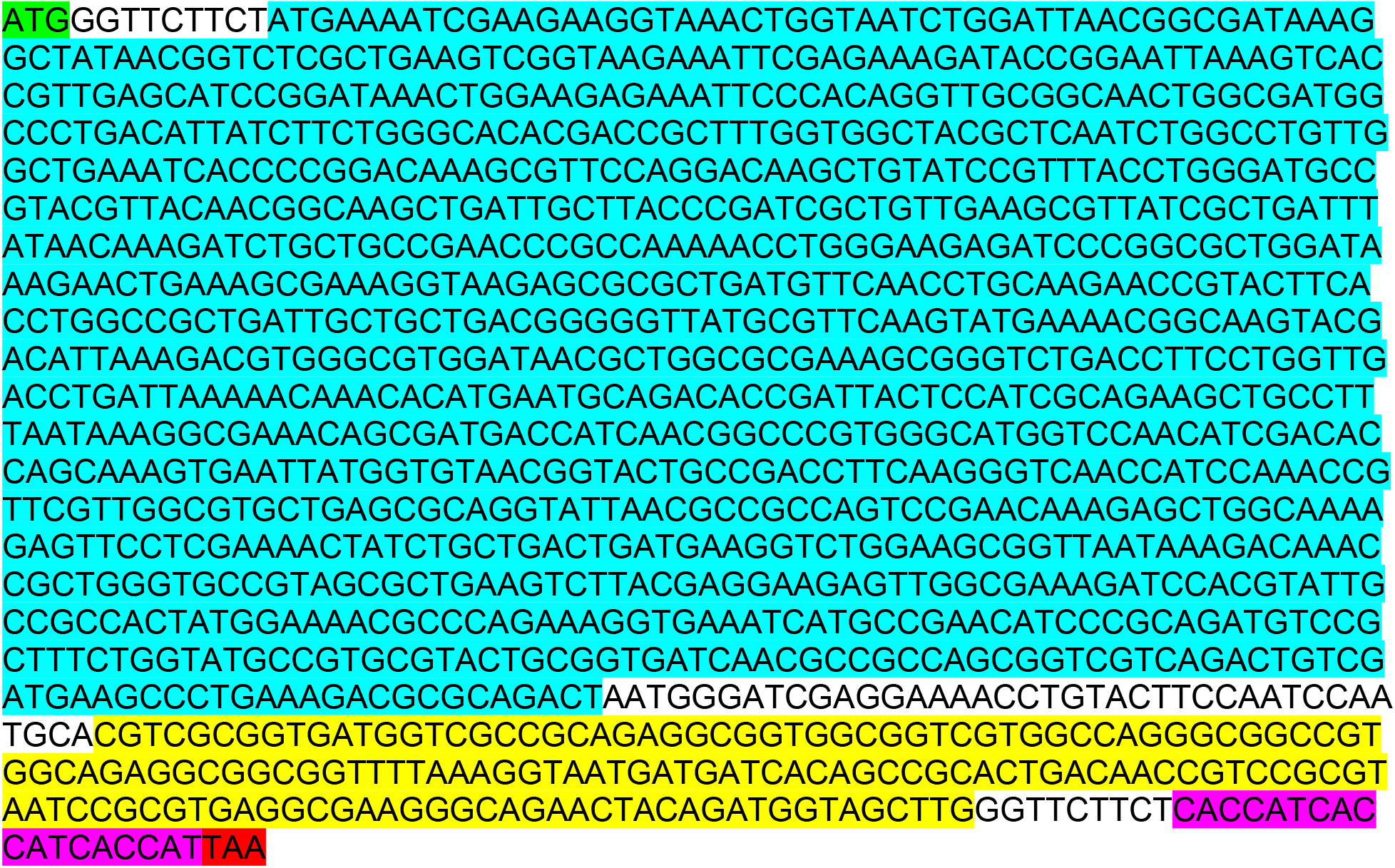

### MBP-(RGG+CTD Δ55)-His6

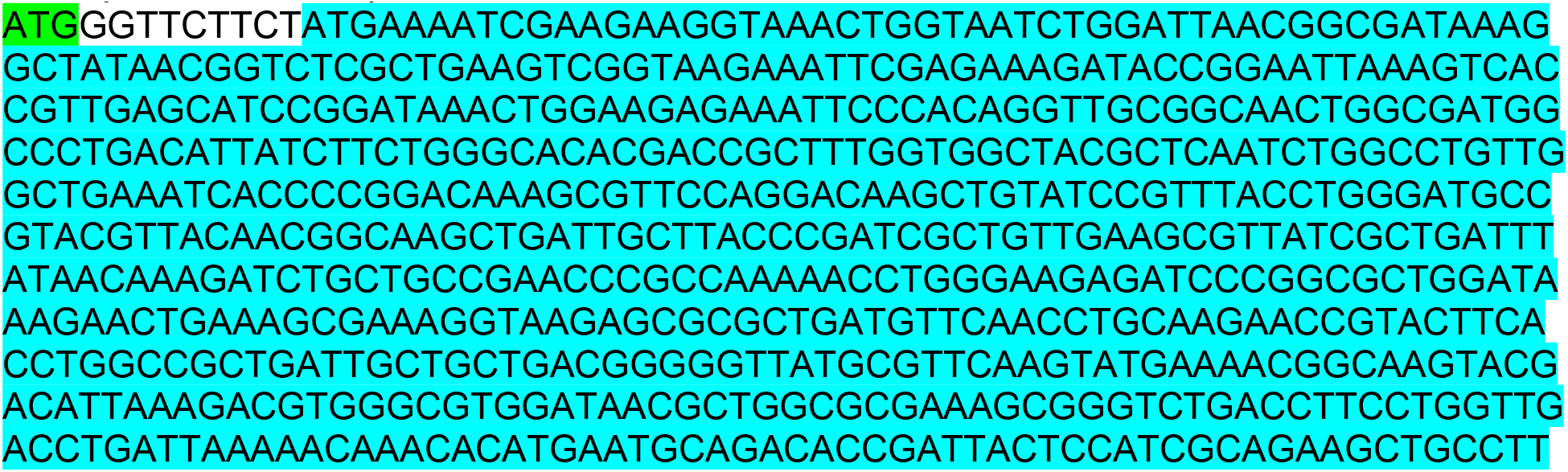
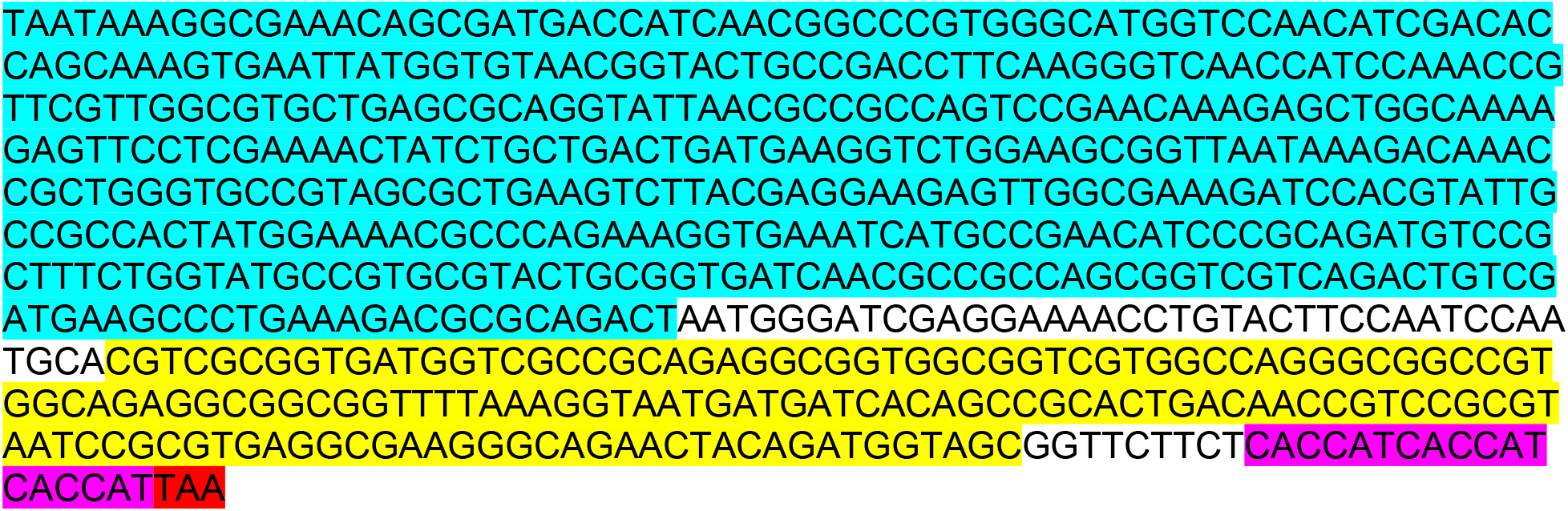

### MBP-(RGG+CTD Δ62)-His6

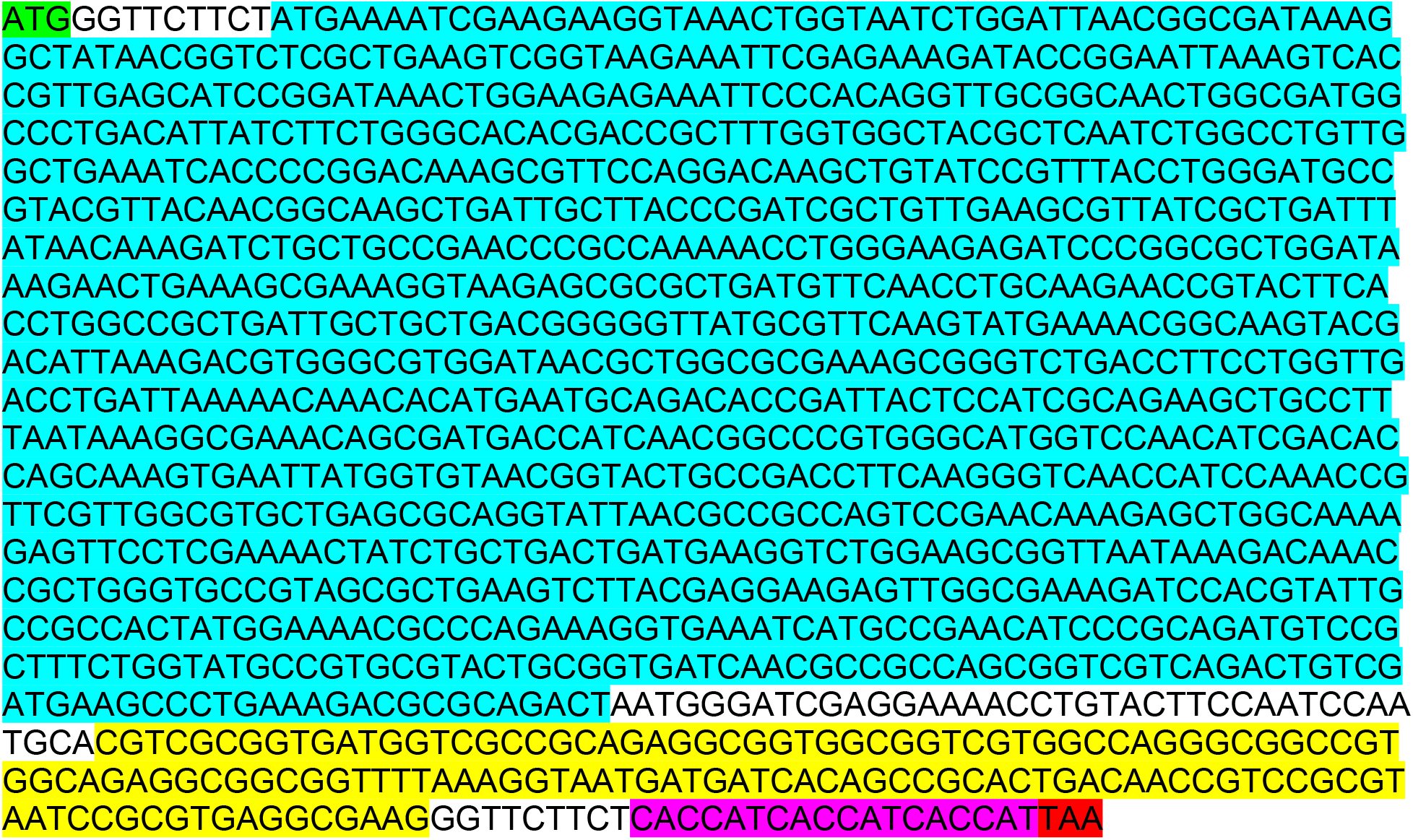

